# A toxin-antitoxin system ensures plasmid stability in *Coxiella burnetii*

**DOI:** 10.1101/2022.09.15.508156

**Authors:** Shaun Wachter, Diane C Cockrell, Heather E Miller, Kimmo Virtaneva, Kishore Kanakabandi, Benjamin Darwitz, Robert A Heinzen, Paul A Beare

## Abstract

*Coxiella burnetii* is the causative agent of Q fever. All *C. burnetii* isolates encode either an autonomous replicating plasmid (QpH1, QpDG, QpRS, or QpDV) or QpRS-like chromosomally integrated plasmid sequences. The role of the ORFs present on these sequences is unknown. Here, the role of the ORFs encoded on QpH1 was investigated. Using a new *C. burnetii* shuttle vector (pB-TyrB-QpH1ori) we cured Nine Mile Phase II of QpH1. The ΔQpH1 strain grew normally in axenic media but had a significant growth defect in Vero cells, indicating QpH1 was important for *C. burnetii* virulence. We developed an inducible CRISPR interference system to examine the role of individual QpH1 plasmid genes. CRISPRi of *cbuA0027* resulted in significant growth defects in axenic media and THP-1 cells. The *cbuA0028*/*cbuA0027* operon encodes CBUA0028 and CBUA0027, which are homologous to the HigB2 toxin and HigA2 anti-toxin, respectively, from *Vibrio cholerae*. Consistent with toxin-antitoxin systems, overexpression of *cbuA0028* resulted in a severe intracellular growth defect that was rescued by co-expression of *cbuA0027*. CBUA0028 inhibited protein translation. CBUA0027 bound the *cbuA0028* promoter (P*cbuA0028*) and CBUA0028, with the resulting complex binding also P*cbuA0028*. In summary, our data indicates *C. burnetii* maintains an autonomously replicating plasmid because of a plasmid-based toxin-antitoxin system.

## 1 Introduction

*Coxiella burnetii* is an obligate intracellular pathogen and the etiological agent of human Q fever. Other than New Zealand this bacterium has a worldwide distribution (Maurin and Raoult, 1999). Domestic animals, such as sheep, goats and cows, are the main reservoirs for human infection, and their contaminated byproducts provide the vehicle for release of the highly infectious and stable bacterium (Angelakis and Raoult, 2010). Human infections are often asymptomatic (∼60 %) but can present as an acute flu-like illness or under more rare instances manifest as a more serious chronic infection (Eldin et al., 2017). *C. burnetii* has a trophism for mononuclear phagocytes (Khavkin and Tabibzadeh, 1988, Stein et al., 2005) where the bacterium replicates within a large phagolysosomal-like parasitophorous vacuole termed the *Coxiella*-containing vacuole (CCV). During phagolysosomal maturation, *C. burnetii* converts from an environmentally stable small-cell variant (SCV) into a metabolically active large-cell variant (LCV) (Coleman et al., 2004). Replication within the CCV requires the Dot/Icm type IVB secretion system, which delivers essential effector proteins into the cytosol of infected cells. Secretion assays have identified greater than 100 Dot/Icm substrates have been identified in the *C. burnetii* genome (Larson et al., 2016).

The *C. burnetii* genome consists of a small chromosome (1.9 – 2.2 Mb) and either one of at least 4 different autonomously replicating plasmids (QpH1, QpDG, QpDV or QpRS) or a chromosomally integrated QpRS-like plasmid sequence (IPS). The *C. burnetii* plasmids range in size from 32.6 to 54.2 kb while the IPS consists of 17.5 kb (Beare et al., 2009). A comparison of the open reading frame (ORF) content of the autonomous plasmids indicated that 22 open reading frames (ORFs) are conserved (Beare et al., 2009) and of these only 5 ORFs (*cbuA0007a*, *cbuA0010*, *cbuA0013*, *cbuA0023* and *cbuA0027*) are strictly conserved (having an identical ORF size), while a further 3 ORFs (*cbuA0006*, *cbuA0011* and *cbuA0012*) have slightly different sizes when the IPS region is included in the comparison (Beare et al., 2009). Of these 8 ORFs, *cbuA0006*, *cbuA0013* and *cbuA0023* have been identified as encoding Dot/Icm substrates and were found to localize to small cytoplasmic puncta (CBUA0006), autophagosomes (CBUA0013) or displayed no specific localization (CBUA0023), respectively, when ectopically expressed in host cells (Voth et al., 2011). The conservation of these ORFs suggests that they are important for *C. burnetii* pathogenesis.

The recent creation of an axenic media (Omsland et al., 2009) (Sandoz et al., 2016), and its use in generating *himar1* transposon libraries in *C. burnetii* have allow a random method for identifying ORFs essential to bacterial virulence (Martinez et al., 2014, Newton et al., 2014, Weber et al., 2013). Using this approach Martinez *et_al* (Martinez et al., 2014) isolated 14 *himar1* mutants located in 8 different QpH1 ORFs (*cbuA0001*, *cbuA0003*, *cbuA0008b*, *cbuA0023*, *cbuA0032*, *cbuA0036*, *cbuA0039* and *cbuA0040*). Of these 14 mutants, 2 ORFs (*cbuA0036-parB* and *cbuA0039-repA*), with predicted roles in plasmid replication, had both strong replication and internalization defects in infected Vero cells, while two different mutants in *cbuA0023* had either a mild replication defect or no defect. The development of a method for creating targeted gene deletions has allowed further analysis of the role of ORFs in *C. burnetii* pathogenesis (Beare et al., 2012). Using this technique, Colonne *et_al* (Colonne et al., 2016) created gene deletions of *cbuA0015* and *cbuA0016* and showed these mutants had a small atypical CCV in infected THP-1 macrophages (Colonne et al., 2016). Recently a new shuttle vector (pQGK) was developed and was used to cure *C. burnetii* of the QpH1 plasmid (Luo et al., 2021). This QpH1-deficient strain of *C. burnetii* had a significant growth defect in bone marrow derived macrophages (Luo et al., 2021). Together these studies indicate that the ORF content of the *C. burnetii* plasmid is important for virulence.

In this current study, we used a *C. burnetii* shuttle vector containing a new tyrosine-based nutritional selection maker and QpH1 replication machinery (QRM; *cbuA0036*-*cbuA0039a*) to create a ΔQpH1 strain of Nine Mile RSA439 (NMII). The ΔQpH1 strain displayed a severe growth defect in Vero cells. To determine the roles of the ORFs on QpH1 we developed an inducible CRISPR interference (CRISRPi) system, containing a new proline-based nutritional selection marker. CRISPRi was used to individually knockdown expression of all the ORFs on the QpH1 plasmid. Knockdown of *cbuA0027* resulted in severe replication defects in axenic media and in THP-1 macrophage cells. Sequence analysis of *cbuA0027* indicated it was part of an operon with *cbuA0028* and that the products of these ORFs encoded an antitoxin and toxin, respectively. Overexpression analysis, pulldown assays and electrophoretic mobility shift assays (EMSA) confirm CBUA0028 and CBUA0027 are a functional non-canonical type II toxin-antitoxin system present on the QpH1 plasmid.

## 2 Results

### 2.1 Deletion of the QpH1 plasmid results in defective intracellular growth

Every *C. burnetii* strain sequenced to date has either an endogenous plasmid (QpH1, QpRS, QpDG, QpDV) or a QpRS-like chromosomal insertion. To examine the role of the plasmid in *C. burnetii* virulence we constructed a plasmid-less strain of Nine Mile phase II. To delete the endogenous plasmid, we constructed a *C. burnetii*-*Escherichia coli* shuttle vector termed pB-TyrB-QpH1ori that contains the QpH1 replication machinery (QRM), *E. coli* origin of replication (pMB1), selectable marker (CAT) for *E. coli* and a new *C. burnetii* nutritional selection marker. This marker is based on complementation of tyrosine auxotrophy, whereby *C. burnetii* expressing *tyrB* and grown in the presence of 4-hydroxyphenylpyruvate (4-HPP) will make L-tyrosine (Fig 1a and 1b). Introduction of pB-TyrB-QpH1ori into *C. burnetii* and selection in axenic media in the absence of tyrosine resulted in the curing of the endogenous QpH1 plasmid (ΔQpH1) (Fig 1c). Growth of ΔQpH1 was indistinguishable to that of wild-type NMII in axenic medium (Fig 2a), with both strains increasing in more than 2500-fold genome equivalents (GE) over 7 days. Replication in Vero cells following a 6-day infection was then examined (Fig 2b). The ΔQpH1 strain displayed a significant reduction in growth (21.9-fold GE increase) compared to NMII (577-fold GE increase) (Fig 2b). In infected Vero cells ΔQpH1 was contained within a small compressed CD63-positive vacuole (Fig 2c and 2d), which is like the phenotype seen for some Dot/Icm substrate mutants (Larson et al., 2015). These results are consistent with previous data showing transposon mutants in the *parB* (*cbuA0036*) and *repA* (*cbuA0039*) ORFs located on QpH1 that have strong replication defects in Vero cells (Martinez et al., 2014).

**Figure 1.**
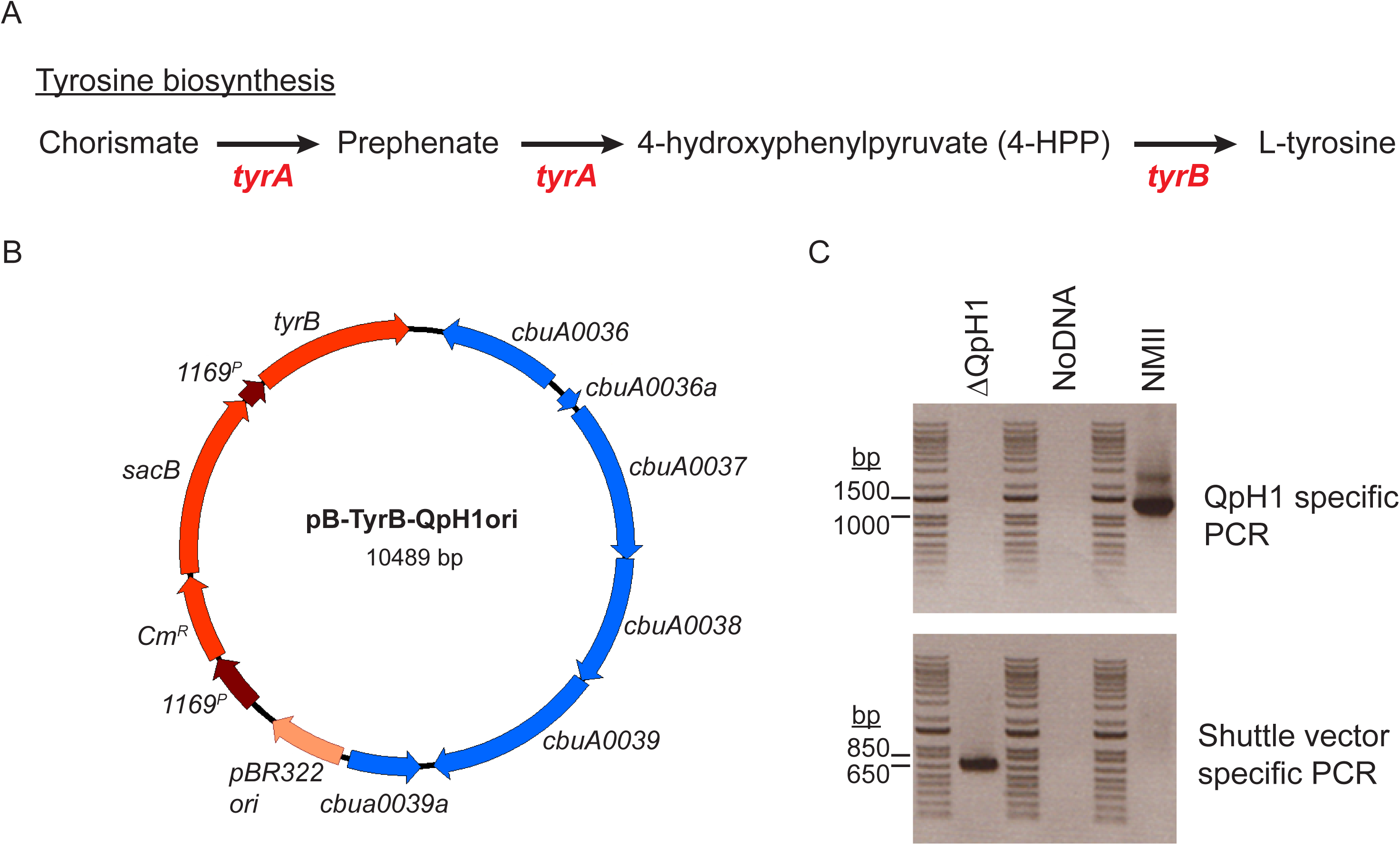
Curing NMII of the QpH1 plasmid. (a) Schematic of the tyrosine biosynthesis pathway. *C. burnetii* is auxotrophic for L-tyrosine and missing both *tyrA* and *tyrB* genes required to convert chorismate into L-tyrosine. (b) Plasmid map of the pB-TyrB-QpH1ori shuttle vector. (c) PCR detection of a QpH1 specific region (P*cbuA0023*-*cbuA0023*) and a shuttle specific gene (*cm^r^*). Oligonucleotide primer pairs used for PCR detection are listed in Table S2. PCR was carried out using ΔQpH1 and NMII gDNA as template DNA. A no DNA template control was also used. The 1 Kb Plus DNA ladder (Invitrogen) was used for size comparison (bands are sized at 15,000, 10,000, 8,000, 7,000, 6,000, 5,000, 4,000, 3,000, 2,000, 1,500, 1,000, 850, 650, 500, 400, 300, 200, 100 bp). The QpH1 specific PCR band (1,235 bp) was not detected in the ΔQpH1 strain but was present in NMII, while the shuttle vector specific PCR band (650 bp) was only seen in ΔQpH1. These data indicate that QpH1 has been cured from ΔQpH1.

**Figure 2.**
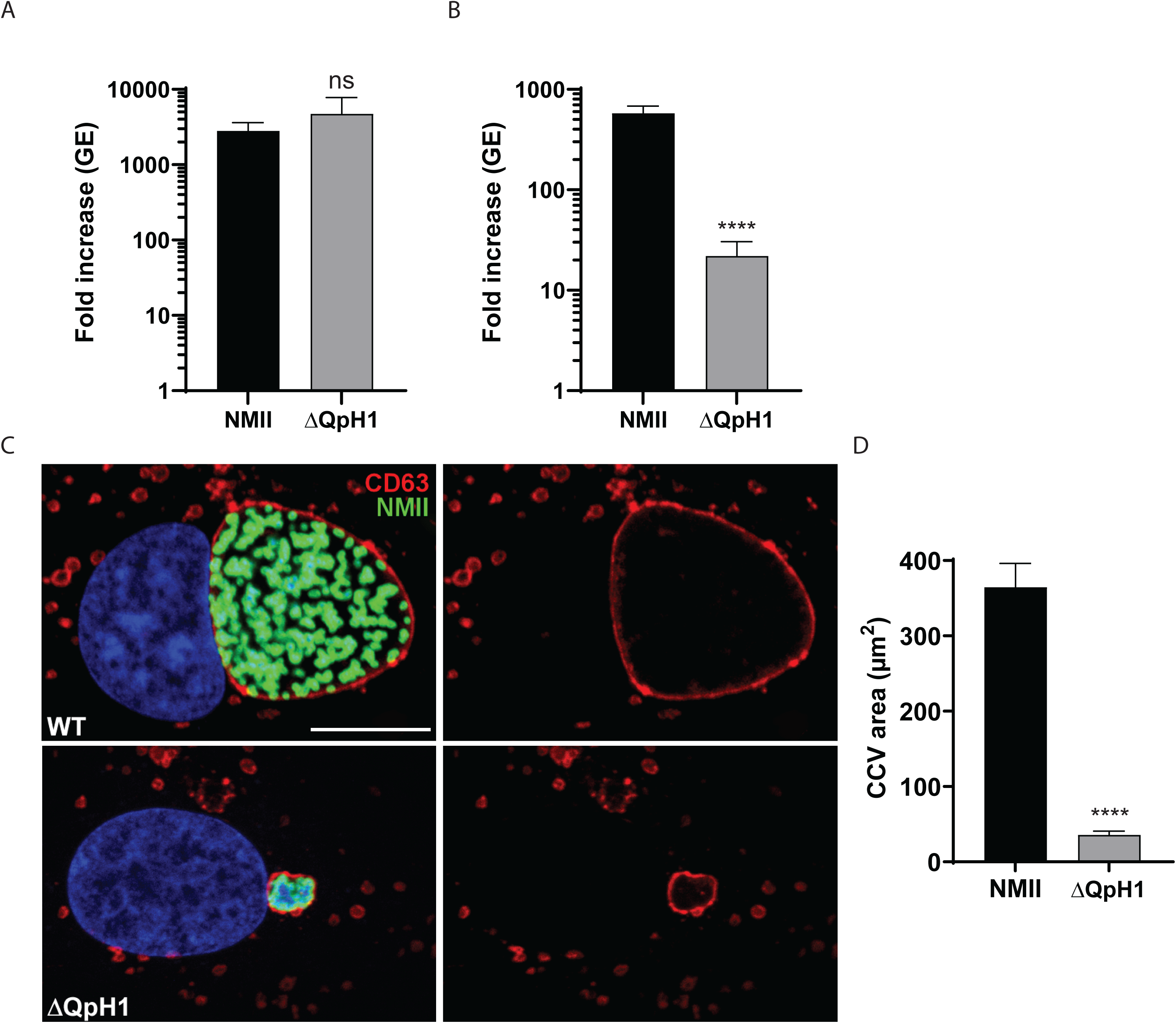
*C. burnetii* ΔQpH1 has a severe intracellular growth defect. Replication of wild-type NMII *C. burnetii* and the ΔQpH1 mutant in (a) ACCM-D and (b) Vero cells. Replication is shown as fold increase in *C. burnetii* genome equivalents (GE) in axenic media (7 days) and Vero cells (6 days). Results are expressed as the means of results from two biological replicates from three independent experiments. Error bars indicate the standard deviations from the means, and asterisks indicate a statistically significant difference (Student’s *t* test, **** - P < 0.0001) compared to values for wild-type *C. burnetii*. ns indicated no significant difference. (c) Confocal fluorescence micrographs of Vero cells infected for 3 days with NMII and ΔQpH1 mutant. CD63 (red) and *C. burnetii* (green) were stained by indirect immunofluorescence, and DNA (blue) was stained with Hoescht 33342. Bar, 10 μm. (d) The histogram depicts the mean intensity of CCV area +/- SD of >50 cells for at least three independent experiments. Statistical difference was determined using the Student’s *t* test (**** - P < 0.0001).

### 2.2 CRISPR interference of QpH1 ORF expression identifies a potential plasmid-based toxin-antitoxin system

To investigate the requirement of the plasmid ORFs for virulence, we developed a miniTn7T-based inducible CRISPR interference (CRISPRi) system for knockdown of *C. burnetii* ORFs (Fig 3a). The *C. burnetii* CRISPRi system contains isopropyl β-D-1-thiogalactopyranoside (IPTG) inducible expression of 3x-c-myc-tagged dCas9 and one or more targeting single guide RNA (sgRNA), while selection of CRISPRi transformants was achieved using complementation of the proline auxotrophy in *C. burnetii* (Fig 3b). The CRISPRi system was tested for its ability to repress expression of single (Fig S1) and multiple (Fig S2) *C. burnetii* ORFs. Complementation of the CRISPRi knockdown system was achieved using an anhydrotetracycline (aTc) inducible system expressing a codon optimized version of the knocked down ORF, incorporating silent mutations into the codons targeted by the sgRNA (Fig S1). CRISPRi was used to target each of the plasmid ORFs contained in QpH1. Three targeting sequences were chosen for each ORF, except for *cbuA0023a* that only had two nGG PAM sequences, and cloned upstream of the sgRNA sequence. The effect of CRISPRi-based QpH1 ORF knockdown on growth in axenic media (Fig S3) and in THP-1 macrophages (Fig S4) was examined following 6- and 5-day incubations, respectively. Induction of the CRISPRi system was analyzed by immunoblot analysis of dCas9 production (Fig S5). The Nine Mile QpH1 plasmid encodes 8 Dot/Icm type IVB secretion substrates (Maturana et al., 2013). Knockdown of the ORFs encoding these proteins resulted in no defect in either ACCM-D growth (Fig 4a) or in THP-1 macrophage growth (Fig 4b) suggesting that they are dispensable for growth under these conditions. Targeted knockdown was examined by qRT-PCR for one target for each type 4B secretion substrate ORF (Fig S6) and showed knockdown of *cbuA0006*, *cbuA0013*, *cbuA0014*, *cbuA0015*, *cbuA0023* and *cbuA0025* but not *cbuA0016* or *cbuA0034*. Although *cbuA0016*, the downstream part of an operon with *cbuA0015*, presumably would also be knocked down in the *cbuA0015* CRISPRi strain. Only one ORF, *cbuA0027*, displayed significant growth defects in both axenic media (Fig 5a and Fig S3) and THP-1 cells (Fig 5b and Fig S4) with all CRISPRi target sequences tested. Targeted knockdown of *cbuA0027* was confirmed by qRT-PCR (Fig S7). Complementation of *cbuA0027* CRISPRi knockdown was achieved using an aTc inducible codon-optimized version of *cbuA0027* (Fig 6a) that contained a modified DNA sequence in the region targeted by the *cbuA0027* CRISPRi constructs (Fig 6b). Complementation resulted in restoration of intracellular growth in THP-1 macrophages (Fig 6c). The *cbuA0027* ORF is located downstream of *cbuA0028* forming a two-gene operon (Fig 7a) (Mao et al., 2009, Mao et al., 2014). A structural-based I-Tasser analysis of CBUA0028 and CBUA0027 indicated that they are homologous to the HigB2 toxin and HigA2 antitoxin from *Vibrio cholerae*, respectively (Fig 7b and c). CBUA0028 displays 26% identity with HigB2 and conservation of the five amino acids (K49, R51, R64, Y82 and K84) previously shown to comprise the HigB2 active site (Hadzi et al., 2017). CBUA0027 has 30% identity with HigA2 and displays a conserved helix-turn helix motif that is essential for DNA binding (Park et al., 2020). These data suggested CBUA0028 and CBUA0027 function as a toxin and antitoxin, respectively.

**Figure 3.**
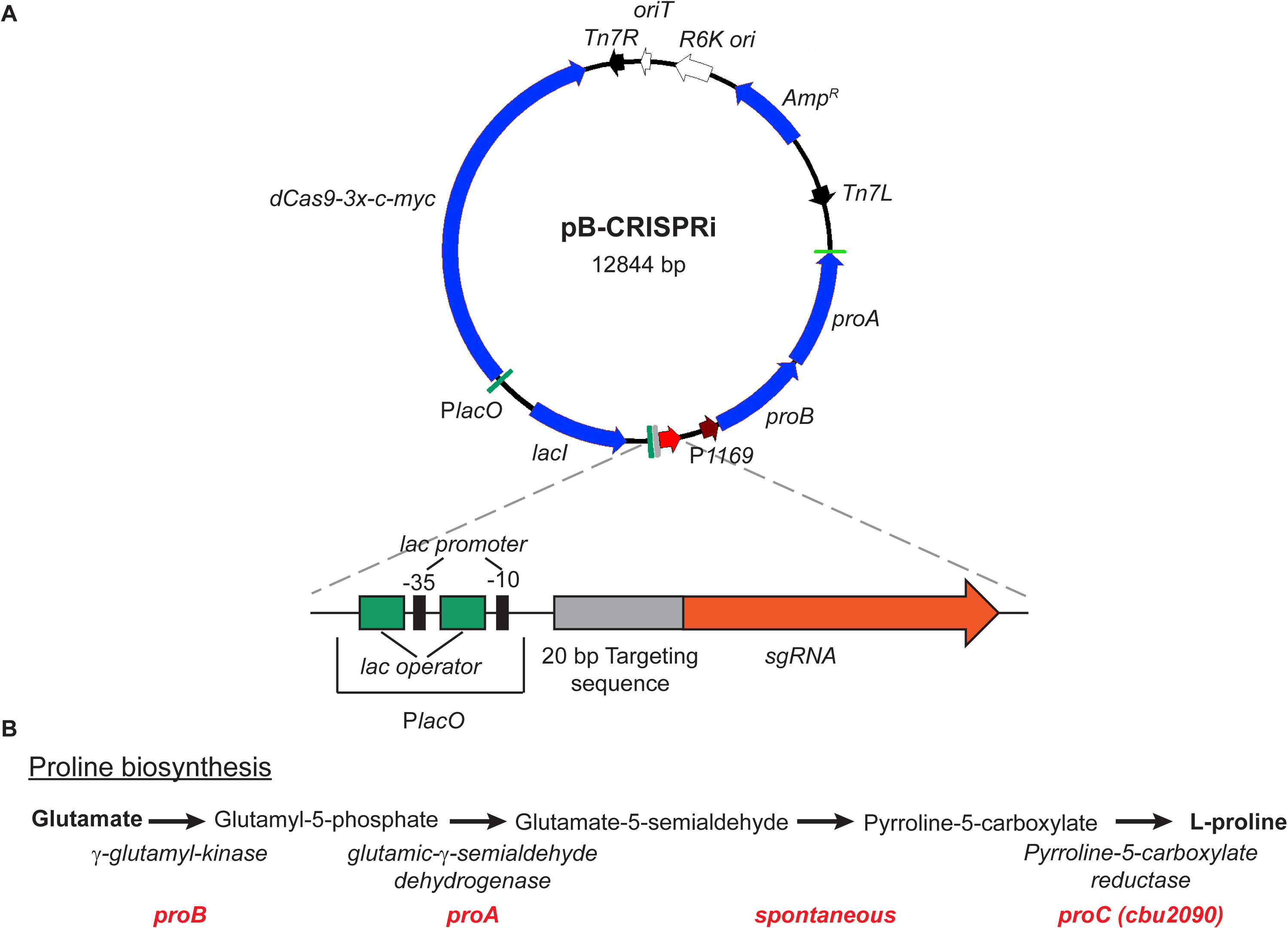
Components of the inducible *C. burnetii* CRISPRi system. (a) Plasmid map of pB-CRISPRi. Proline complementation was achieved by cloning the *proBA* operon from *Legionella pneumophilia* in front of the *cbu1169* promoter (P*1169*). Inducible expression of dcas9 and sgRNA is controlled by the *lac* promoter (P*lacO*). The PlacO-sgRNA region is enlarged and depicts two *lac* operators flanking the lac -35 promoter sequence upstream of the 20 bp targeting sequence fused to sgRNA. (b) Schematic of the proline synthesis pathway. The genes required to convert glutamate to proline are depicted in red. *C. burnetii* encodes *proC* (*cbu2090*) but is missing *proA* and *proB*.

**Figure 4.**
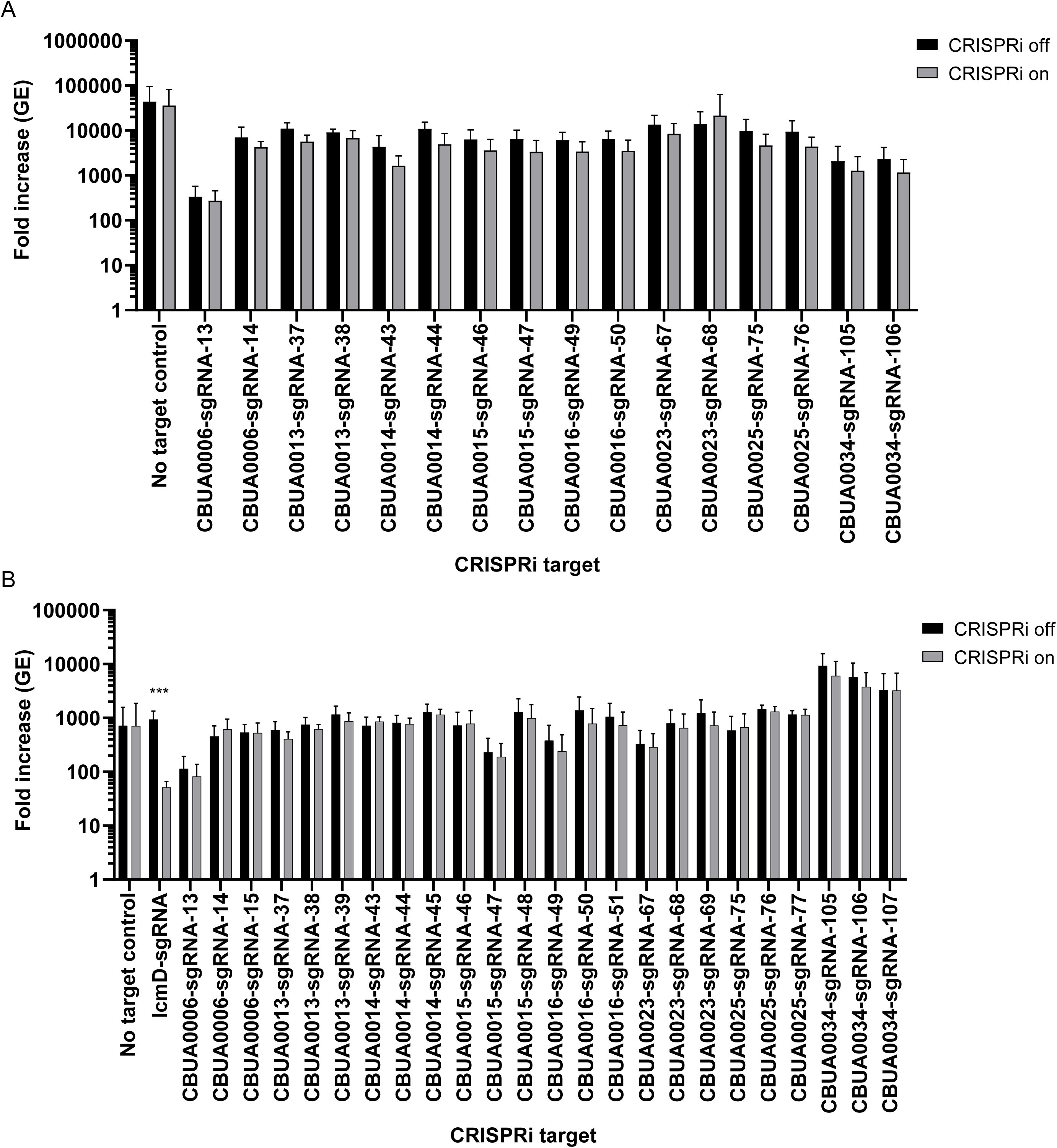
CRISPRi repression of QpH1 Dot/Icm subtrate expression does not affect axenic or THP-1 macrophage growth. Replication is shown as fold increase in *C. burnetii* genome equivalents (GE) of Dot/Icm substrates CRISPRi strains after 6 days in (a) ACCM-D or after a 5-day infection of (b) THP-1 macrophages in the absence (CRISPRi off) or presence (CRISPRi on) of CRISPRi system inducer molecule IPTG. Replication of a no sgRNA target control was also examined as a negative control. CRISPRi repression of the essential *icmD* Dot/Icm apparatus gene was used as a positive control in THP-1 macrophages. Results are expressed as the means of results from two biological replicates from three independent experiments. Error bars indicate the standard deviations from the means, and asterisks indicate a statistically significant difference (Student’s *t* test, *** - P < 0.001) compared to values for the CRISPRi off samples.

**Figure 5.**
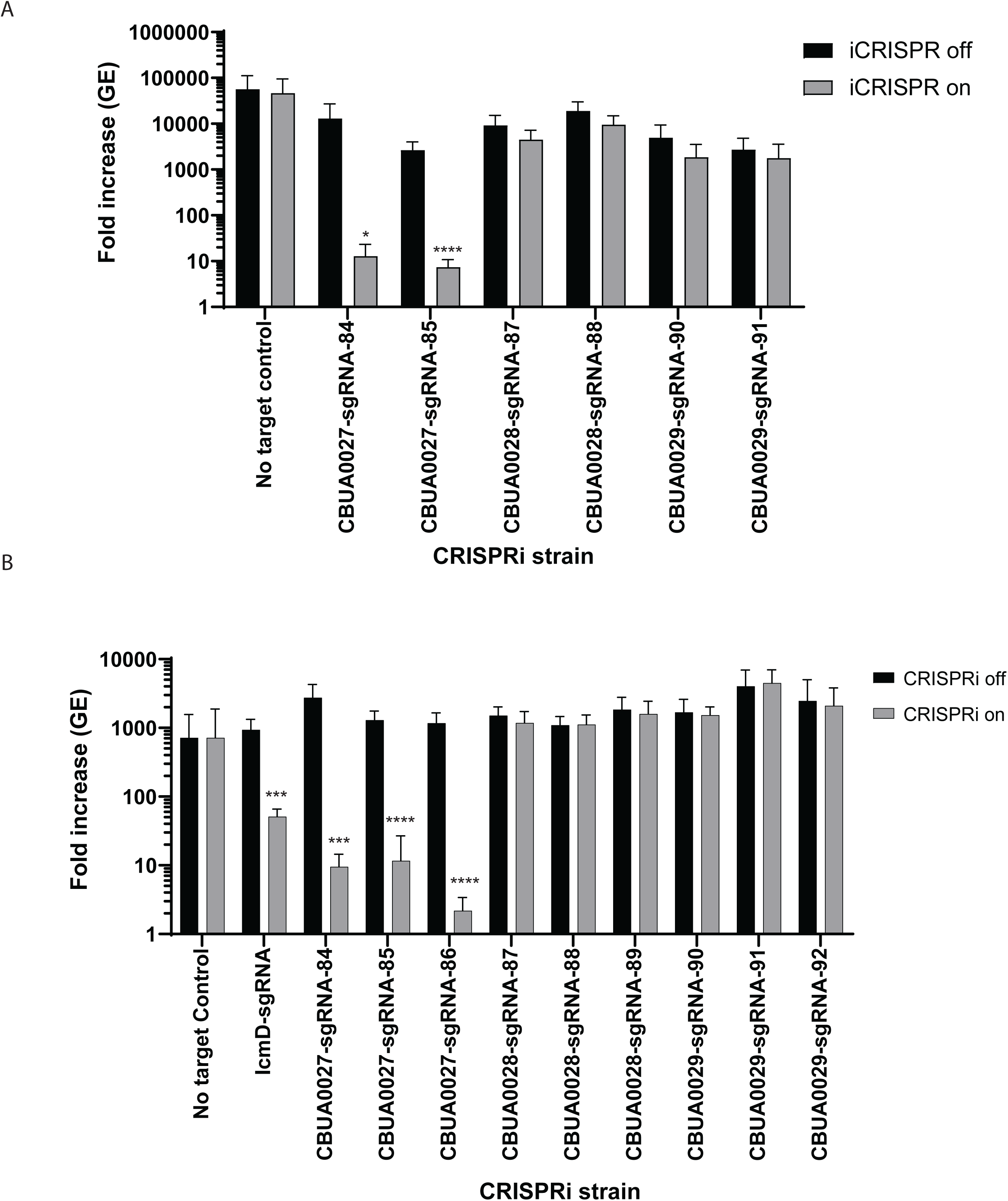
CRISRPi repression of *cbuA0027* expression causes a severe growth defect in axenic media and THP-1 macrophages. Replication is shown as fold increase in *C. burnetii* genome equivalents (GE) of CRISPRi strains targeting *cbuA0027*, *cbuA0028* and *cbuA0029* genes after 6 days in (a) ACCM-D or after a 5-day infection of (b) THP-1 macrophages in the absence (CRISPRi off) or presence (CRISPRi on) of CRISPRi system inducer molecule IPTG. Replication of a no sgRNA target control was also examined as a negative control. CRISPRi repression of the essential *icmD* Dot/Icm apparatus gene was used as a positive control in THP-1 macrophages. Results are expressed as the means of results from two biological replicates from three independent experiments. Error bars indicate the standard deviations from the means, and asterisks indicate a statistically significant difference (Student’s *t* test, * - P < 0.05, *** - P < 0.001, and **** - P < 0.0001) compared to values for the CRISPRi off samples.

**Figure 6.**
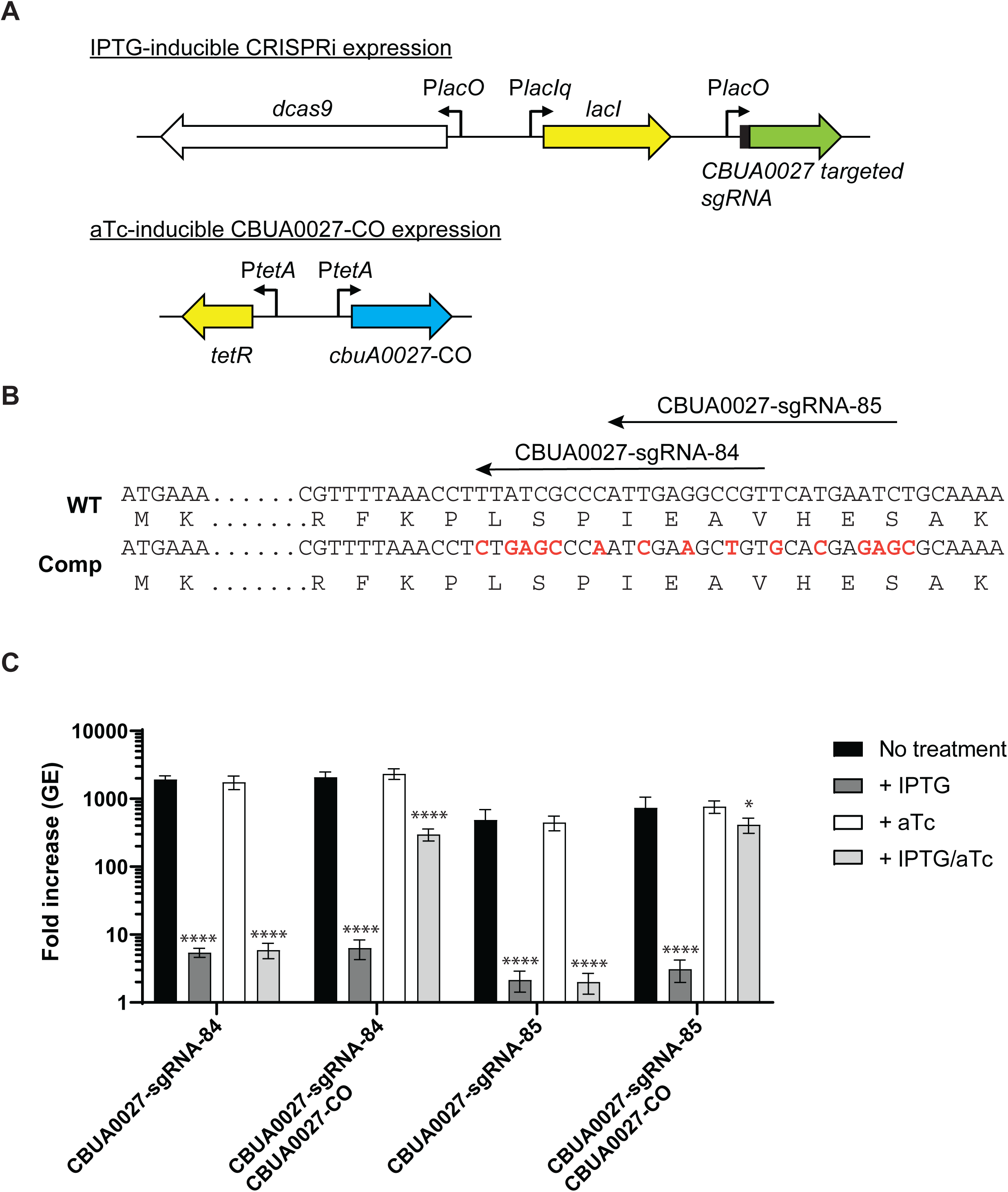
Complementation of *cbuA0027* CRISPRi repression allows restoration of intracellular growth. (a) Schematics of the IPTG-inducible *cbuA0027* CRISPRi repression system and aTc-inducible expression of the codon optimized *cbuA0027* gene. In the presence of IPTG expression of dCas9-3x-c-myc and *cbuA0027* targeted sgRNA occurs from the P*lac* promoters. In the presence of aTc expression of the codon optimized *cbuA0027* gene occurs from the P*tetA* promoter. (b) Partial sequences of the wild-type (WT) and codon optimized *cbuA0027* gene are depicted. The location of the two *cbuA0027* sgRNA targeting sequences are shown with arrows. Nucleotides that were changed are depicted in red. For *cbuA0028*-sgRNA-84 9 bp of the target sequence was changed and for *cbuA0028*-sgRNA-85 8 bp of the target sequence was changed. These changes did not affect the protein sequence. (c) Expression of *cbuA0027*-CO allows complementation of THP-1 macrophage growth in *cbuA0027* CRISPRi respression strains. Replication is shown as fold increase in *C. burnetii* genome equivalents (GE) of *cbuA0027* CRISPRi strains with or without the *cbuA0027*-CO complement plasmid (pJB-lysCA-TetRA-*cbuA0027*-CO) after a 5-day infection of THP-1 macrophages in the absence of any inducer (No treatment; CRISPRi off/Complement off), presence of IPTG (CRISPRi on/Complement off), presence of aTc (CRISPRi off/Complement on) or presence of both IPTG and aTc (CRISPRi on/Complement on). Results are expressed as the means of results from two biological replicates from three independent experiments. Error bars indicate the standard deviations from the means, and asterisks indicate a statistically significant difference (One way ANOVA, * - P < 0.05, **** - P < 0.0001) compared to values for the no treatment samples.

**Figure 7.**
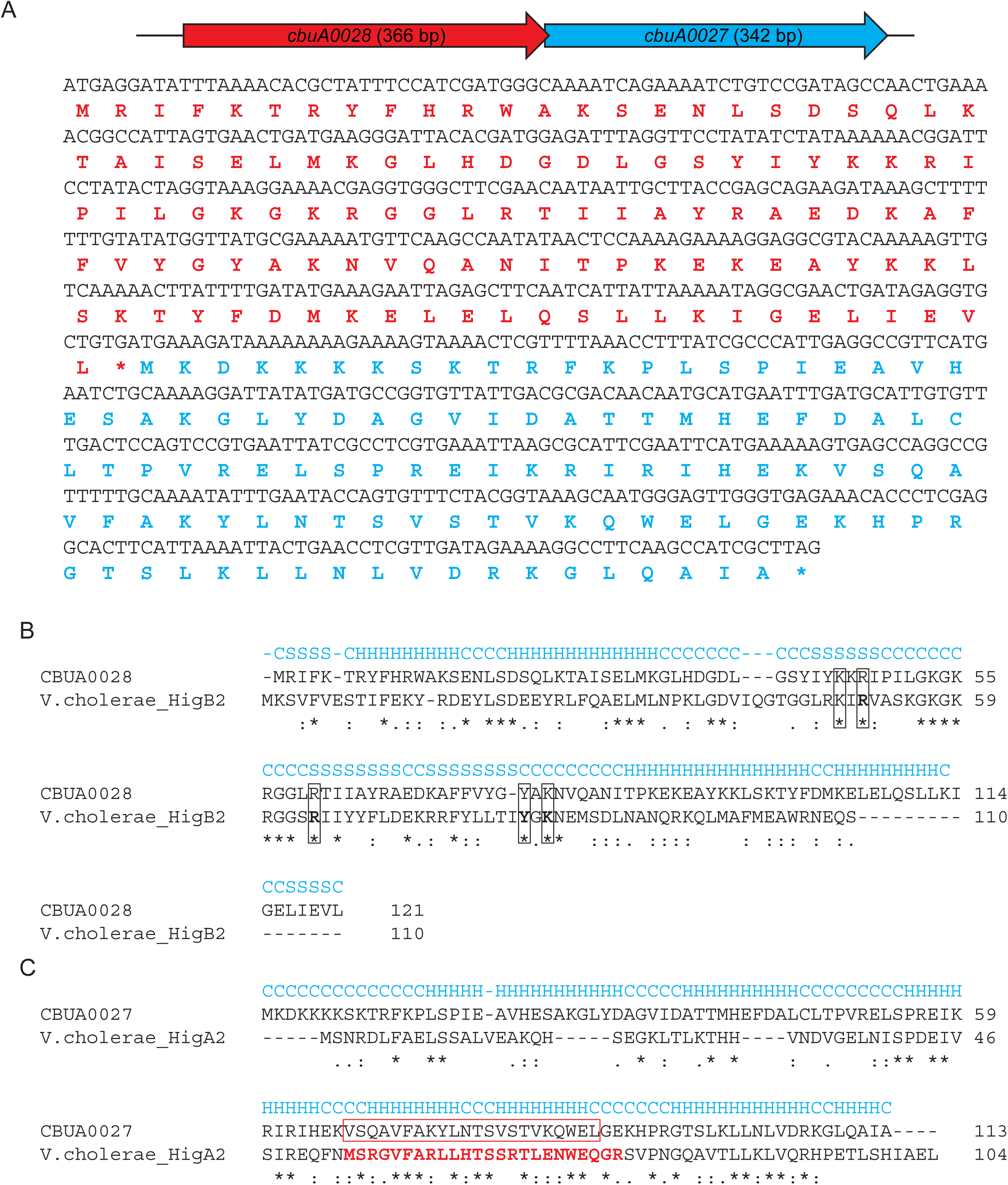
The CBUA0028/CBUA0027 TA modules retain essential regions of the Vibrio cholerae HigBA2 TA system. (a) Schematic of the *cbuA0028*/*cbuA0027* operon, showing their toxin-antitoxin gene orientation, and their corresponding DNA and amino acid sequences. The CBUA0028 and CBUA0027 amino acid sequences are depicted in red and blue. (b) I-Tasser structural-based alignment of CBUA0028 and HigB2 toxin of V. cholerae. Secondary structure prediction is denoted in blue (C – coil, H – helix, S – strand). Identical residues are depicted with an asterisk (*) while similar residues are denoted as a single dot (.) or colon (:). The location of essential active site residues in HigB2 (K49, R51, R64, Y82 and K84) are boxed and are conserved in CBUA0028. (C) I-Tasser structural-based alignment of CBUA0027 and HigA2 antitoxin of V. cholerae. Secondary structure prediction is denoted in blue (C – coil, H – helix, S – strand). Identical residues are depicted with an asterisk (*) while similar residues are denoted as a single dot (.) or colon (:). The residues comprising the essential helix-turn-helix DNA binding motif in HigA2 is colored red. The predicted helix-turn-helix DNA binding motif in CBUA0027 is highlighted by a red box and aligns with the same motif in HigA2.

### 2.3 Overexpression of *cbuA0028* inhibits growth of *C. burnetii*

To ascertain whether CBUA0028 functions as a toxin, *cbuA0028* was cloned into the low-copy miniTn7T transposon vector, whereby its expression was controlled by aTc-inducible promoter (Fig 8a; pMiniTn7T-proBA-TetRA-*cbuA0028*). Expression of *cbuA0028* using its own promoter was not possible in *E. coli* due to the toxic effects of CBUA0028, resulting in mutations to the cloned *cbuA0028* ORF (data not shown). One mutation of note, resulted in a lysine to glutamic acid change at residue 79 in CBUA0028 that had no toxic effect on *E. coli* growth (data not shown), presumably because K79 is predicted to be one of the active site residues in CBUA0028 (Fig 7b). The pMiniTn7T-proBA-TetRA-*cbuA0028* expression vector was transformed into ΔQpH1, containing a clean background absent of the native *cbuA0028* and *cbuA0027* ORFs. Transformation was only possible using ΔQpH1 containing an IPTG-inducible *cbuA0027* construct (ΔQpH1/CBUA0027) (Fig 8b), with prior induction of *cbuA0027* necessary for survival of the pMiniTn7T-proBA-TetRA-*cbuA0028* transformants. The *cbuA0028* ORF is conserved in all *C. burnetii* except genomic group V (Beare et al., 2006), the group containing the chromosomally integrated QpRS-like sequences. Within this genomic group the *cbuA0028* ORF is N-terminally truncated (Fig S8a) and the *cbuA0028* promoter is missing while the complete *cbuA0027* ORF is intact (Fig S8b). We decided to test if this truncated ORF (*cbuA0028G*) could also function as a toxin (Fig 8c). The effect of *cbuA0028* and *cbuA0028G* overexpression was investigated by examining *C. burnetii* growth in THP-1 macrophages in the presence and absence of the putative anti-toxin CBUA0027 (Fig 8d). Induction of *cbuA0027* alone had no effect on *C. burnetii* growth. Overexpression of *cbuA0028* resulted in a severe growth defect in THP-1 macrophages, with an approximate 2-fold GE increase over a 5-day infection (Fig 8d). This growth defect was alleviated by co-expression of *cbuA0027*. Overexpression of *cbuA0028*G had no effect on *C. burnetii* growth in THP-1 macrophages. In the absence of any treatment less growth was seen in the ΔQpH1/CBUA0027/CBUA0028 strain suggesting that a background level of *cbuA0028* expression was occurring (Fig 8d). This data shows *cbuA0028* expression prevents *C. burnetii* growth in the absence of CBUA0027 and suggests the N-terminus of CBUA0028 is essential to its function.

**Figure 8.**
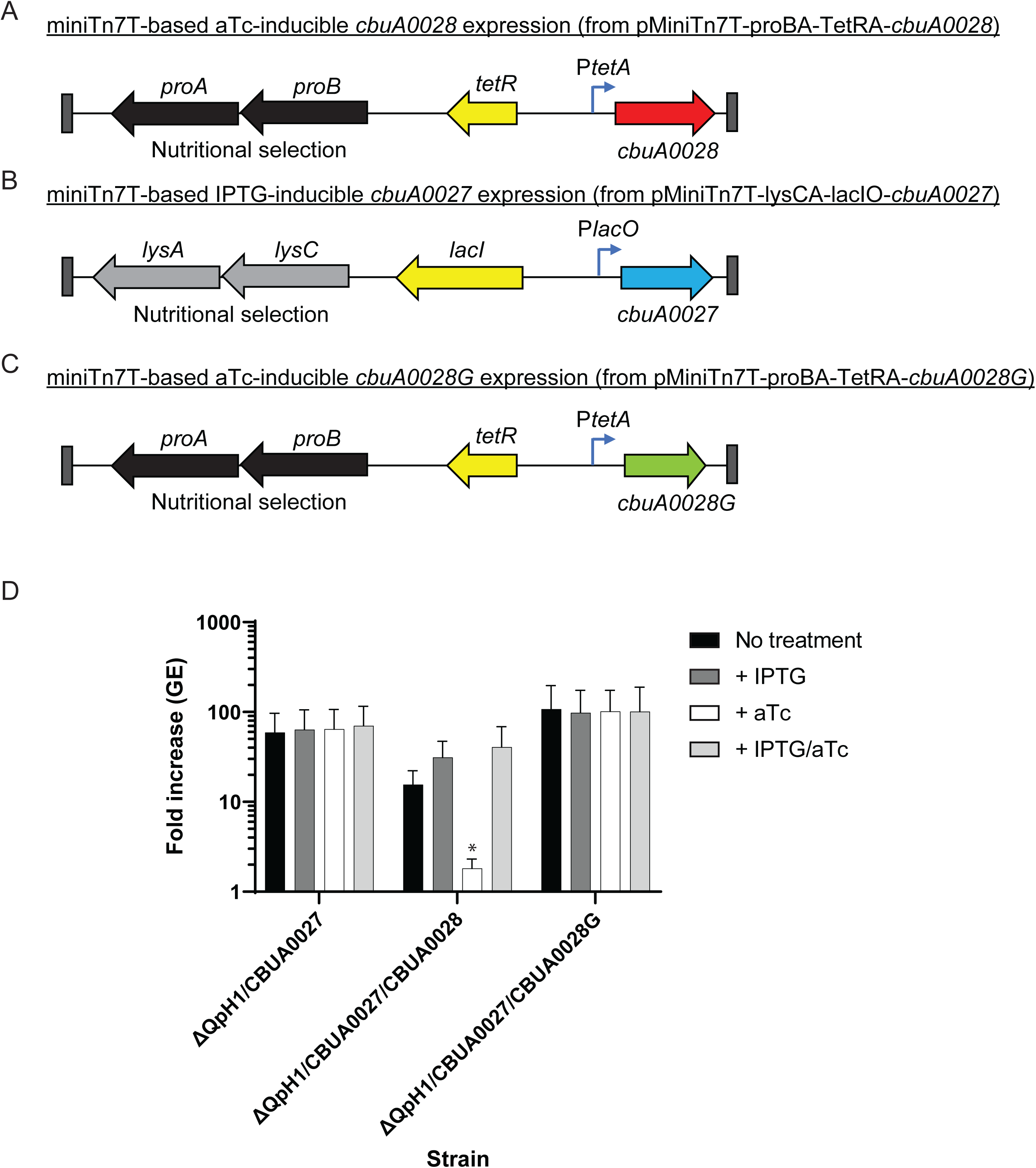
Expression of *cbuA0028* in the absence of CBUA0027 prevents *C. burnetii* replication. Schematic of miniTn7T transposons designed to allow (a) aTc-inducible expression of *cbuA0028* (from pMiniTn7T-proBA-TetRA-*cbuA0028*) or (b) IPTG-inducible expression of *cbuA0027* (from pMiniTn7T-proBA-lacIO-*cbuA0027*) or (c) aTc-inducible expression of the N-terminally truncated version of *cbuA0028* from G Q212 (*cbuA0028G*) (from pMiniTn7T-proBA-TetRA-*cbuA0028G*). The ΔQpH1 mutant was first transformed with the pMiniTn7T-lysCA-lacIO-*cbuA0027* using the lysine-based nutritional selection to integrate the IPTG-inducible into the *C. burnetii* genome. The resulting strain ΔQpH1/CBUA0027 was treated with IPTG to induce *cbuA0027* expression then transformed with miniTn7T transposons containing either aTc-inducible *cbuA0028* or *cbuA0028G* expression using proline-base nutritional selection to create ΔQpH1/CBUA0027/CBUA0028 and ΔQpH1/CBUA0027/CBUA0028G, respectively. (d) Expression of *cbuA0028* caused a severe growth defect in THP-1 macrophages that could be alleviated by co-expressing *cbuA0027*. Replication is shown as fold increase in *C. burnetii* genome equivalents (GE) of the ΔQpH1/CBUA0027 strain with or without the CBUA0028 or CBUA0028G miniTn7T transposons after a 5-day infection of THP-1 macrophages in the absence of any inducer (No treatment), presence of IPTG, presence of aTc or presence of both IPTG and aTc. Results are expressed as the means of results from two biological replicates from three independent experiments. Error bars indicate the standard deviations from the means, and asterisks indicate a statistically significant difference (One way ANOVA, * - P < 0.05) compared to values for the no treatment samples.

### 2.4 CBUA0028 production prevents translation of co-produced proteins

The HigB2 toxin has a RelE-like active site and when bound to the ribosome A-site cleaves translating mRNA molecules without a preference for specific sequences (Hadzi et al., 2017). Due to the conservation of the HigB2 active site residues in CBUA0028 we wanted to explore whether CBUA0028 is functioning in the same manner. To investigate this phenomenon, a T7 polymerase-based cell-free expression system containing a control gene expression plasmid, expressing the type 4B secretion substrate *cbu0665* (pEXP1-*cbu0665*), and decreasing amounts of *cbuA0028* expression plasmid (pET28a-*cbuA0028*) as DNA templates was used. Production of N-terminally XpressT-tagged CBU0665 and C-terminally V5-tagged CBUA0028 protein was then examined by immunoblot (Fig 9a and b). The results show a dose-responsive increase in the level of CBU0665 when decreasing amounts of CBUA0028 were present. Co-production of CBUA0027 rescued the effect of CBUA0028 on CBU0665 production. These data confirm that CBUA0028 prevents production of an unrelated protein (CBU0665) and that CBUA0027 can alleviate this effect, results that are consistent with CBUA0028 and CBUA0027 being a toxin and antitoxin, respectively.

**Figure 9.**
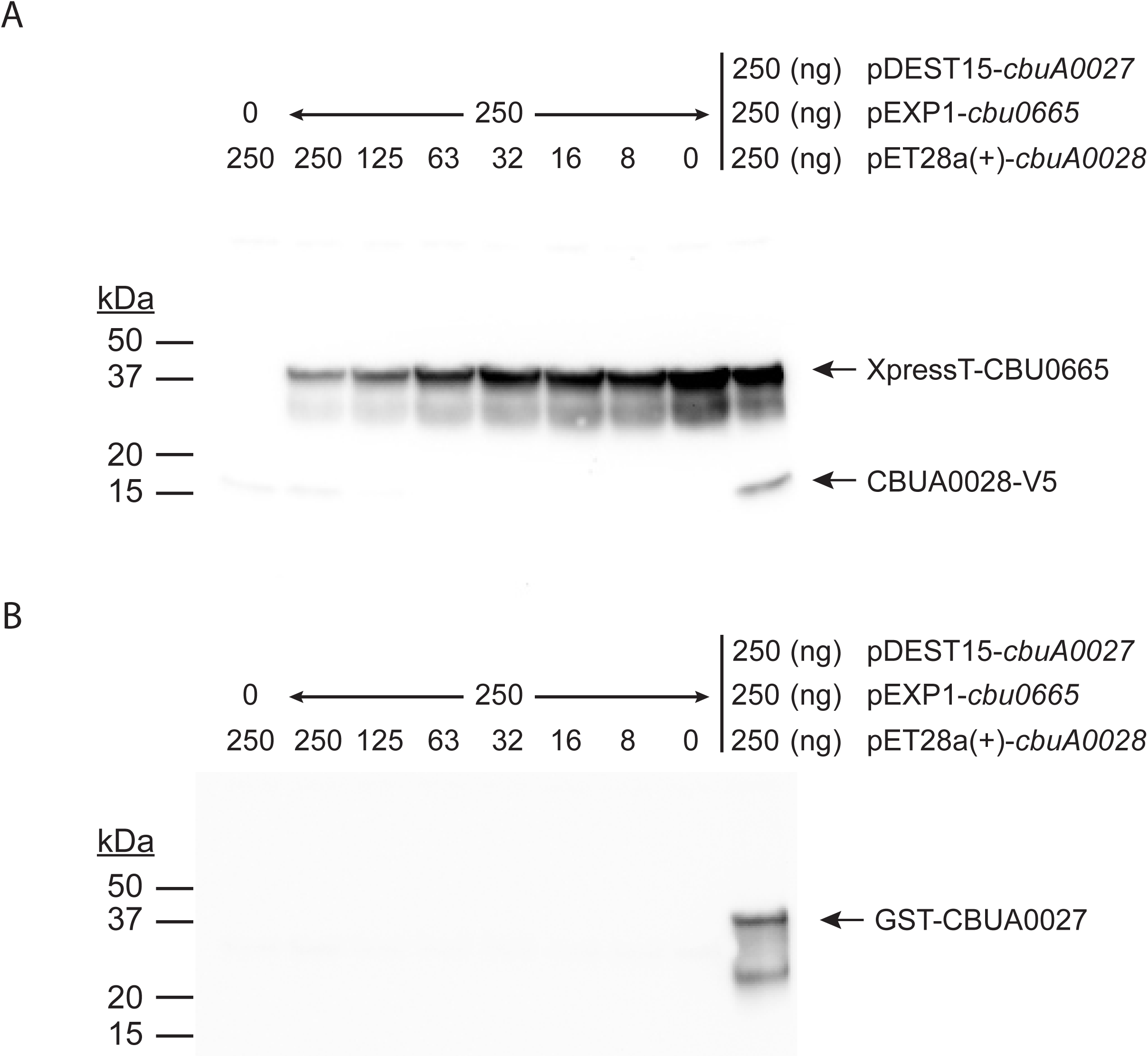
Production of CBUA0028 inhibits protein translation. Cell-free *in vitro* transcription and translation (IVTT) reactions incubated with pEXP1-*cbu0665* (250 ng) and decreasing amounts of pET28a(+)-*cbuA0028* (250 ng – 8 ng) as DNA template. Control IVTT reactions were conducted with either only pEXP1-*cbu0665* (250 ng) or pET28a(+)-*cbuA0028* (250 ng) plasmids or with all three plasmids together (250 ng of pEXP1-*cbu0665*, pET28a(+)-*cbuA0028* and pDEST15-*cbuA0027*) as template DNA. (a) Production of CBU0665 (42.5 kDa) and CBUA0028 (18 kDa) was detected by immunoblot of cell-free IVTT lysates probed simultaneously with anti-XpressT and anti-V5 antibodies, respectively. (b) Production of CBUA0027 (40.1 kDa) was detected by immunoblot of cell-free lysates probed with anti-GST antibody. A dose-response affect was seen on CBU0665 production when decreasing amounts of CBUA0028 were present indicating CBUA0028 inhibited translation of CBU0665. Translation inhibition of CBU0665 production by CBUA0028 was alleviated when CBUA0027 was present.

### 2.5 The CBUA0027 anti-toxin directly binds the CBUA0028 toxin

The HigB2 and HigA2 proteins are part of the RelE superfamily of type II TA systems (Christensen-Dalsgaard and Gerdes, 2006). In type II TA systems, the anti-toxin binds to the toxin to prevent degradation of ribosome-associated mRNA (Fraikin et al., 2020). To confirm that CBUA0028 and CBUA0027 are part of a type II TA system, pulldowns with N-terminally tagged CBUA0027 were investigated. Cell-free expression was used to produce GST-CBUA0027 or simultaneously produce GST-CBUA0027 and CBUA0028-V5 or XpressT-CBU0665 (Fig S9). Cell-free extracts were then incubated with GST-binding beads to extract proteins bound to GST-CBUA0027 and these complexes were eluted from the beads with a reduced glutathione buffer. Immunoblots of the pulldowns indicated specific binding between CBUA0027 and CBUA0028 (Fig 10), while the control pulldown with CBU0665 displayed no binding with CBUA0027. No CBUA0028 protein was seen in the GST pulldown elutions containing CBUA0028 alone. Together, these data indicate that CBUA0027 directly binds CBUA0028 confirming its function as an antitoxin.

**Figure 10.**
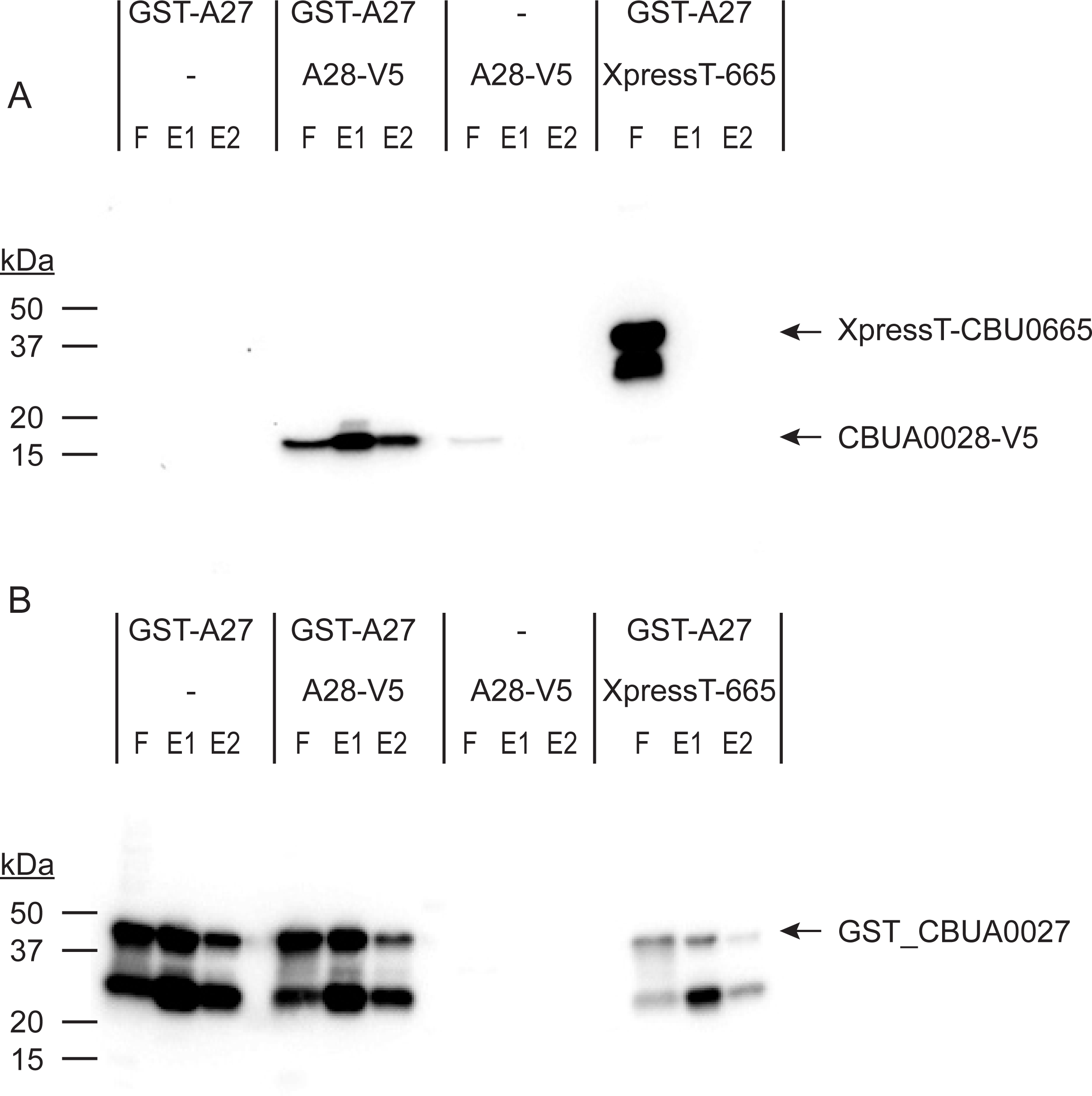
CBUA0027 binds directly to CBUA0028. Cell-free IVTT reactions containing GST-tagged CBUA0027 (GST-A27), GST-A27 and V5-tagged CBUA0028 (A28-V5), A28-V5 or GST-A27 and XpressT-tagged CBU00665 (XpressT-665) (Fig S9) were incubated with GST-affinity beads. After incubation beads were washed and bead-bound proteins were eluted using a reduced glutathione buffer. Flow-through (F), elution 1 (E1) and elution 2 (E2) fractions were analyzed by immunoblot probed simultaneously with (a) anti-XpressT and anti-V5 or (b) anti-GST. GST-CBUA0027 pulldown of only CBUA0028-V5 was observed.

### 2.6 CBUA0027 and the CBUA0028/CBUA0027 complex bind the *cbuA0028* promoter

RelE-like antitoxins form a complex with their cognate toxin and this complex, while preventing toxin binding to the ribosome and subsequent mRNA cleavage, also binds to the TA promoter to repress expression of the TA system (Jurenas et al., 2022, Overgaard et al., 2009). RelB alone has also been found to bind the *relBE* promoter (Overgaard et al., 2008). To investigate binding of CBUA0027 and CBUA0027/CBUA0028 complex with the *cbuA0028* promoter (P*cbuA0028*) electrophoretic mobility shift assays (EMSA) were performed. Elutions obtained from the CBUA0027 and CBUA0027/CBUA0028 pulldowns were incubated with labeled P*cbuA0028* or control *groES* promoter and these protein/DNA complexes analyzed. CBUA0027 and the CBUA0027/CBUA0028 complex both resulted in a mobility shift when in the presence of P*cbuA0028* but not the control *groES* promoter (Fig 11 and Fig S10) indicating specific DNA binding with *PcbuA0028*.

**Figure 11.**
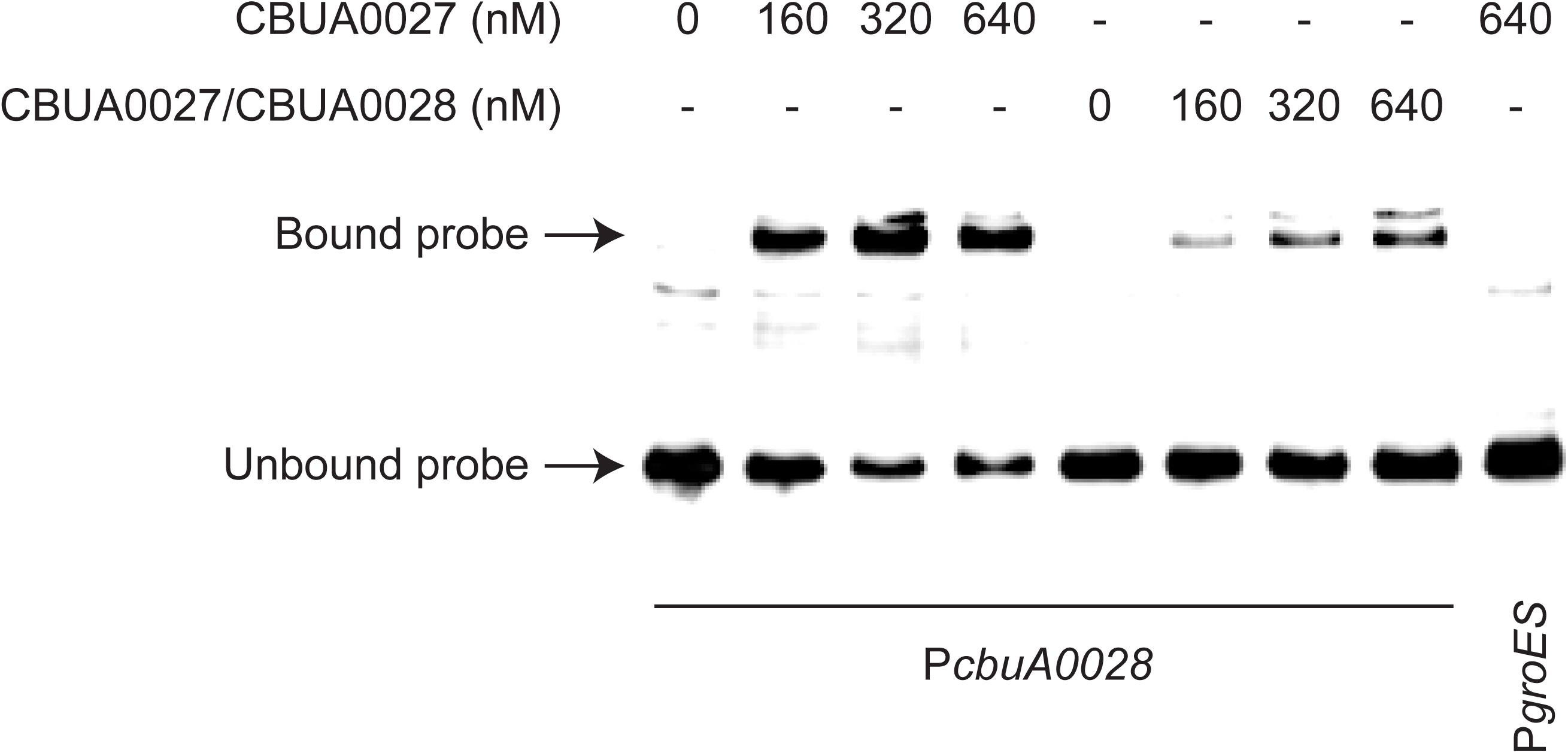
CBUA0027 and CBUA0027/CBUA0028 complex bind to the *cbuA0028* promoter. EMSAs show interactions between biotin-labeled *cbuA0028* promoter (P*cbuA0028*) and increasing concentrations of purified CBUA0027 or CBUA0027/CBUA0028 complex. Biotin-labeled P*groES* with CBUA0027 (640 nM) was included as negative control. The location of bound and unbound probe is depicted by arrows.

### 2.7 *cbuA0028* expression is repressed in the presence of CBUA0028/CBUA0027

Canonical type II TA systems have a promoter-toxin-antitoxin arrangement, that allows for a molar excess of antitoxin via translational coupling. However, non-canonical type II systems, such as *mqsRA*, display a promoter-antitoxin-toxin organization and in this configuration (Fraikin et al., 2019), one would expect that there would be a molar excess of toxin. TA systems like *mqsRA* alleviate this problem using promoters present within the toxin gene (Fraikin et al., 2019). The *cbuA0028*/*cbuA0027* TA system displays a non-canonical promoter-toxin-antitoxin arrangement (Fig 7a). We predicted that an additional promoter(s) would be present within *cbuA0028* that would allow P*cbuA0028*-independent expression of *cbuA0027*. The motif prediction tool in Geneious Prime was used to predict potential sigma 70 binding sites. Two potential *cbuA0027* promoters (P2 and P3) were identified and compared to P*cbuA0028* (P1) (Fig 12a and Fig S11), with the P3 promoter also still present in the G Q212 *cbuA0028* sequence (Fig S11). To test if these promoters are transcriptionally active in *C. burnetii* fragments containing either P1 (PA28-F1), both P2 and P3 (PA27-F1) or just the P3 region (PA27-F2) were fused to the red fluorescent protein mScarlet-i (Fig 12b) and a single copy inserted into wild type NMII and ΔQpH1 using the miniT7nT transposon system (Beare et al., 2011). Promoter activity was analyzed by measuring fluorescence at days 3, 5, 7 and 14 to compare promoter strength in LCVs and SCVs and between NMII and ΔQpH1. Expression from P*cbuA0028* was only detected in ΔQpH1 and was highest at Day 14, indicating in the absence of CBUA0028 and CBUA0027 the P*cbuA0028* promoter becomes dysregulated. No promoter activity was seen with the putative *cbuA*0027 promoters suggesting that some other mechanism regulates the levels of CBUA0028 and CBUA0027 in *C. burnetii* to maintain a balance between toxin and antitoxin.

**Figure 12.**
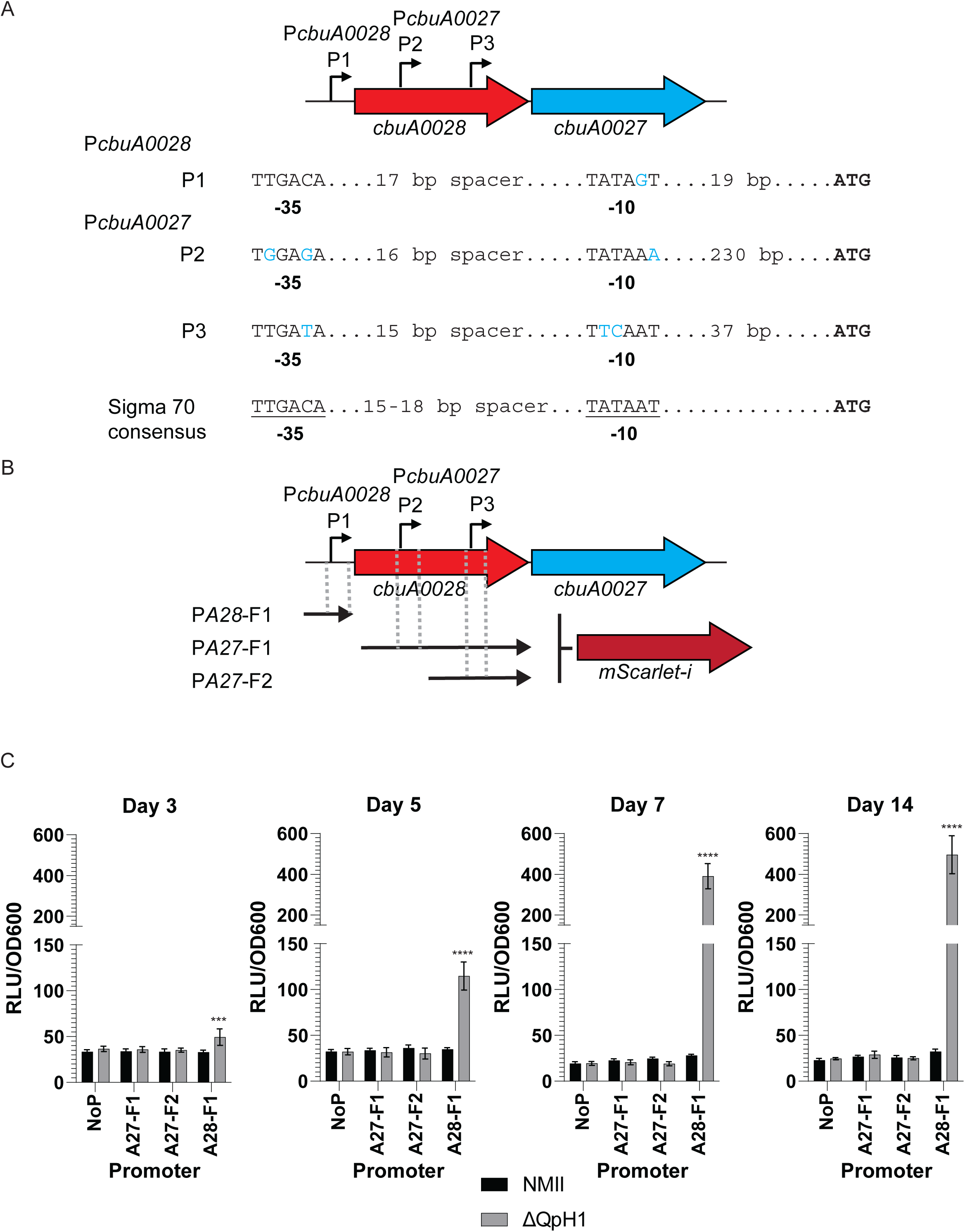
Identification of potential *cbuA0027* promoters within the *cbuA0028* coding sequence. (a) Schematic of the *cbuA0028*-*cbuA0027* operon and location of *cbuA0028* and *cbuA0027* promoters. The sequence of each predicted promoter (-35 and -10 regions) was compared to the consensus sigma 70 promoter, and deviations to this sequence are depicted in blue. (b) The DNA fragments used to create transcriptional fusions to *mScarlet-i* and their location in relation to *cbuA0027* and *cbuA0028* are depicted. (c) Activity of the transcriptional fusions in NMII and ΔQpH1 strains. Axenic medium was inoculated with *C. burnetii* and incubated for 3, 5, 7, or 14 days then fluorescence was measured. Absorbance at OD600 was used as a proxy for *C. burnetii* growth and used to normalize bacteria fluorescence. Results are expressed as the means of results from two biological replicates from four independent experiments. Error bars indicate the standard deviations from the means, and asterisks indicate a statistically significant difference (Two way ANOVA, *** - P < 0.001, **** - P < 0.0001).

## 3 Discussion

All *C. burnetii* strains contain an autonomously replicating plasmid (QpH1, QpDG, QpDV or QpRS) or IPS (Beare et al., 2006) suggesting that ORFs encoded by these DNA sequences are necessary for *C. burnetii* pathogenesis. In this study, we investigated the role of the ORFs found on QpH1. To initiate this research, we first cured the native QpH1 plasmid from NMII using a shuttle vector containing the QRM. Incompatibility between the shuttle vector QRM and the QRM on QpH1 and the use of a new nutritional-based selection for the shuttle vector in *C. burnetii* would cure NMII of QpH1 to generate ΔQpH1. Based on the known QpH1 ORF content the resulting strain would grow normally in axenic media. Plasmid incompatibility was recently used as a method to cure NMII of the QpH1 plasmid (Luo et al., 2021). Growth analysis of the ΔQpH1 strain found no growth defect in axenic media but the presence of QpH1 was essential for growth in Vero cells. Consistent with this finding, transposon mutants located in the *parB* and *repA* ORFs of the *C. burnetii* QpH1 plasmid have strong replication defects in Vero cells (Martinez et al., 2014). A recent study also reported the curing of QpH1 from NMII and showed that QpH1-deficient bacterium had a severe growth defect in murine bone marrow-derived macrophages, although no growth defect was observed in Buffalo green kidney cells and growth defects in THP-1 macrophages were inocula dependent (Luo et al., 2021). Due to the sequence conservation of the QRM ORFs within the 4 known *C. burnetii* plasmids, plasmid curing of QpDG, QpDV and QpRS (Beare et al., 2009) is possible using our shuttle vector and may give insight into the role of these plasmids in pathogenesis of different isolates, such as the K (Q154) and Dugway isolates that have unique sets of Dot/Icm substrates (Maturana et al., 2013).

Site-specific gene deletion in *C. burnetii* remains a challenging and lengthy process and to date has only resulted in 24 mutants (Beare et al., 2014, Clemente et al., 2018, Colonne et al., 2016, Cunha et al., 2015, Friedrich et al., 2021, Larson et al., 2019, Larson et al., 2013, Pechstein et al., 2020, Sandoz et al., 2021, Schäfer et al., 2020, Vallejo Esquerra et al., 2017, Larson et al., 2015) (Long et al., 2021, Stead et al., 2018, Moormeier et al., 2019), two of which are located on QpH1 (Colonne et al., 2016). To overcome the difficulty in creating large numbers of individual gene deletions, we developed a miniTn7T-based IPTG-inducible CRISPRi system to knockdown ORFs in *C. burnetii*. This system integrates as a single copy in the *C. burnetii* chromosome and selection for transformants is modulated using a new nutritional-based marker for proline. The *dcas9* we employed originated from *Streptococcus pyogenes* and recognizes the nGG protospacer adjacent motif (PAM) site (Peters et al., 2019). Previous attempts to use the dCas9 from *Staphylococcus aureus* and *Streptococcus thermophilus* were unsuccessful (data not shown). The number of *S. pyogenes* dCas9 PAM sites in the NMII genome is substantial, with 197,777 and 3,073 nGG sites present in the chromosome and QpH1 plasmid, respectively. The CRISPRi system was first tested by knocking down production of the host-cell essential Dot/Icm protein IcmD (Fig S1) and then by simultaneously knocking down both IcmD and the small SCV-specific protein ScvA production (39 amino acids) (Fig S2). This result indicated that the inducible CRISPRi system could target genes essential for intracellular infection and small SCV- specific genes. Constitutive CRISPRi knockdown of the Dot/Icm substrate CirB in *C. burnetii* was recently described (Fu et al., 2022). This shuttle vector-based system adopts an always on approach using the *cbu1169* promoter to drive both sgRNA and *dcas9* expression. While this system is functional, it is limited for use on non-essential genes for growth in axenic media. During testing of our CRISPRi system we found that expression of large amounts of dCas9 affected growth of *C. burnetii*. Also, bacteria containing knocked down essential genes would eventually start to replicate due to mutation of *dcas9*, the inducible promoter or sgRNA sequences or because of secondary compensatory mutations elsewhere in the genome (data not shown), thus becoming inert to further CRISPRi induction. The benefit of our inducible CRISPRi system is the ability to knockdown genes at any time during the bacterial growth cycle, target essential genes, and multiple genes at once and to function in both axenic media and infected host cells.

Complementation of gene deletions is a marker of Koch’s molecular postulates (Falkow, 1988). We sought to fulfill these postulates for CRISPRi repressed genes. Complementation of CRISPRi knockdowns in *C. burnetii* was achieved using aTc-inducible expression of a codon optimized version of the targeted gene *in trans* (Fig 6c and Fig S1), whereby silent mutations were made in the 20 bp target region and/or PAM sequence. Complementation of CRISPRi knockdown has also been achieved in *Chlamydia trachomatis*, where the complementing *incA* gene was transcriptionally fused 3′ to the aTc-inducible *dcas9* (Ouellette et al., 2021). In this case the CRISPRi target against *incA* was in the promoter region and so dCas9 would not bind to the plasmid-based complement. A recent study by Brockett etal (Brockett et al., 2021) reported that transcriptional fusion of a modified *Chlamydial bacA* gene, containing silent mutations to the CRISPRi target region, could complement CRISPRi knockdown of the chromosomal *bacA* gene. Together these data indicate that CRISPRi repression can be complemented by simultaneously expressing a codon optimized version of the knocked down gene.

The Dot/Icm type IVb system is essential for *C. burnetii* virulence (Beare et al., 2011, Carey et al., 2011) and many of its substrates are required for bacteria growth and CCV biogenesis (Friedrich et al., 2021, Larson et al., 2013, Larson et al., 2015, Martinez et al., 2014, Newton et al., 2014, Weber et al., 2013). CRISPRi targeting of each individual plasmid ORFs found no effect of knockdown of the Dot/Icm substrates located on QpH1. The absolute conservation of *cbuA0013* and *cbuA0023* in *C. burnetii* suggests that these genes are important for pathogenesis. It is possible that the products of these genes are redundant and their loss in function is alleviated by other Dot/Icm effectors (Isaac and Isberg, 2014, Luo and Isberg, 2004). Incomplete knockdown of the genes may also result in partial production of the encoded product. This has been observed in our lab for other CRISPRi targeted genes and can be alleviated by co-expressing multiple targets to the same gene (data not shown). CRISPRi silencing can also have unexpected results due to off-targeting of the dCas9-sgRNA complex to other regions of the genome (Zhang et al., 2021). In fact, knockdown of *cbuA0021* resulted in off-target effects on *cbu0334* (*thiDE*) expression, which may have affected growth in axenic media but not growth in THP-1 cells, and possibly *cbu0055* (*ubiA*) which may have affected growth in THP-1 cells (Fig S12). The *thiDE* gene products are required for thiamine biosynthesis (Liu et al., 2022). Mutants of *thiDE* have significant growth defects in media grown bacteria, which can be restored by adding exogenous thiamine (Liu et al., 2022). Hence, reduction in *cbu0334* expression may prevent thiamine production in axenic media but in THP-1 macrophages the bacteria can transport thiamine from the host. UbiA is part of a superfamily of intramembrane prenyltransferases that have roles in cellular respiration and antioxidation (Li, 2016). Therefore, it is possible that off-targeting effects on *cbu0055* may have a greater effect within the toxic environment of the THP-1 macrophage lysosome than in axenic media.

Using the newly developed inducible CRISPRi system we identified a TA system on QpH1 encoded by *cbuA0028* and *cbuA0027*. Historically TA systems were identified as a two-gene module that was associated with the maintenance of conjugative plasmids (Gerdes et al., 1986, Ogura and Hiraga, 1983). TA systems are ubiquitous throughout free-living prokaryotes and have been found on both chromosomal and extrachromosomal genetic elements (Schuster and Bertram, 2013). TA systems can be classified into 8 different types and in most instances the antitoxin is 5′ in the toxin module (Jurenas et al., 2022, Song and Wood, 2020). Toxin modules have a range of cellular effects including impairing degrading RNAs and disrupting translation, disrupting cell wall structure, inducing metabolic stress and DNA replication (Jurenas et al., 2022). Antitoxins are generally either RNA or protein molecules that abrogate the effects of the toxin modules (Jurenas et al., 2022). While TA systems are widespread in free-living prokaryotes, they are generally not found in many host-associated bacterium including *Chlamydia* species and *Borrelia burgdorferi* (Pandey and Gerdes, 2005). TA two-gene modules have been identified in several *Rickettsia* species, while in other intracellular bacteria such as *Orientia tsutsugamushi*, *Wolbachia* species, *Ehrlichia chaffeensis* and *Anaplasma* species only orphan toxin and anti-toxin genes are present (Audoly et al., 2011, Botelho-Nevers et al., 2012, Socolovschi et al., 2013).

The CBUA0028/CBUA0027 TA system is the first toxin-antitoxin identified and characterized in *C. burnetii*. This TA module is both sequence and structurally homologous to *V. cholerae* HigBA2, a type II TA system and member of the RelE superfamily. Type II toxins are most often ribonucleases that either cleave free mRNA, such as HicA and MazF (Christensen et al., 2003, Jorgensen et al., 2009) or as with HigBA2 and the RelE superfamily, act by cleaving mRNAs in a ribosome-dependent manner (Hurley and Woychik, 2009, Pedersen et al., 2003). The data presented here is consistent with other type II TA systems whereby the antitoxin directly binds its cognate toxin (Overgaard et al., 2009) (Fig 10), the toxin prevents growth in the absence of its cognate antitoxin (Heaton et al., 2012) (Fig 8), and toxin production inhibits protein synthesis, which can be restored in the presence of its antitoxin (Schureck et al., 2016) (Fig 9). The *cbuA0028*/*cbuA0027* TA module displays a non-canonical “reverse” gene orientation that is seen in other TA systems including *higBA*, *mqsRA*, *hicAB*, and *rnlAB* where the toxin is upstream of the antitoxin (Deter et al., 2017). In the canonical gene orientation bicistronic transcription of the TA module allows a molar excess of the antitoxin through translational coupling (Fraikin et al., 2020). Non-canonical TA systems require different mechanisms to achieve a molar excess of antitoxin including additional promoters inside the toxin gene coding sequence (Fraikin et al., 2019, Otsuka et al., 2010). Consistent with *rnlB* and *mqsA* antitoxin expression, internal promoter(s) were identified within the CBUA0028 toxin coding sequence (Fig 12). Interestingly, transcriptional activity was not seen from these putative promoters in *C. burnetii* under the conditions tested (Fig 12). Previous data from both microarray (Sandoz et al., 2014) and RNAseq experiments (Beare et al., 2014) carried out in axenic media indicated that level of *cbuA0028* transcript is higher than *cbuA0027*. However, mass spectrometry data from *C. burnetii* grown in axenic media, showed that at day 4 more CBUA0027 (Normalized spectral count = 28.27) was detected relative to CBUA0028 (Normalized spectral count = 11.86) (Beare et al., 2014) and that the amount of CBUA0027 increased as the bacteria entered stationary phase (Sandoz et al., 2014). Together these data indicate a disconnect between *cbuA0027* expression and CBUA0027 protein production but show that an excess in CBUA0027 is present within *C. burnetii*. Expression from P*cbuA0028* was detected but only in ΔQpH1 (Fig 12). This data indicated that deletion of the QpH1 plasmid alleviates the transcriptional control of the CBUA0028/CBUA0027 complex on the P*cbuA0028* promoter, consistent with results observed with the *mqsR* promoter activity in a Δ*mqsRA* mutant (Fraikin et al., 2019). CRISPRi repression of *cbuA0027* expression is in line with the toxin-antitoxin molar excess model whereby repression of *cbuA0027* expression will result in a molar excess of CBUA0028 and toxic effects to the bacterial cell (Figs 5, 6, 8 and 9). Restoration of CBUA0027 production by complementation alleviated the effects of CRISPRi repression (Figs 6, 8 and 9). Using the data presented here we developed a model for the CBUA0028/CBUA0027 toxin-antitoxin system (Fig 13).

**Figure 13.**
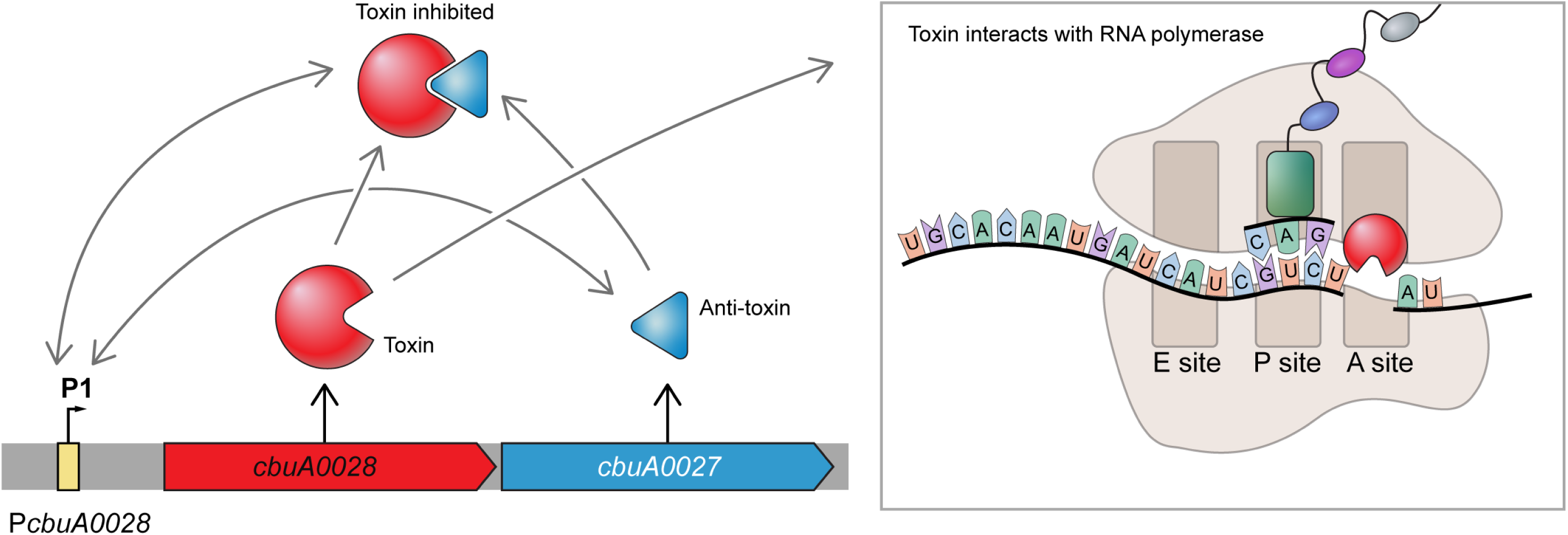
Putative model of CBUA0028/CBUA0027 TA system. The *cbuA0028*/*cbuA0027* operon has a non-conical “reverse” TA gene organization. The first gene encodes the putative mRNA interferase and co-repressor toxin CBUA0028 (red) and the second gene encodes the CBUA0027 antitoxin. The location of the *cbuA0028* promoter (P*cbuA0028*) is depicted upstream of *cbuA0028*. Consistent with the HigB2 toxin from *V. cholerae*, the CBUA0028 toxin is predicted to interact with the RNA polymerase A site where it would cleave mRNA and prevent translation from the corresponding mRNA. The CBUA0027 antitoxin binds CBUA0028, thereby preventing translation inhibition, and this complex can then bind to the P*cbuA0028* promoter resulting in repression of transcription from P*cbuA0028*. CBUA0027 also binds P*cbuA0028* in the absence of CBUA0028. Grey arrows represent interactions between components of the model. Black arrows represent production of protein from *cbuA0028* and *cbuA0027*.

CBU1490 is identified as an addiction molecule antidote protein and its crystal structure has been solved (Franklin et al., 2015). Recently CBU1490 was found to be structurally homologous to the *Mycobacterium tuberculosis* HigA3 antitoxin (Park et al., 2020). Based on the finding of another potential antitoxin in *C. burnetii* we mined the genome for other putative toxin and antitoxins. In addition to the plasmid based CBUA0028/CBUA0027 TA system, ten additional Type II TA modules were found on the *C. burnetii* chromosome (Fig S13, Table 1). This analysis identified CBU1491 as the toxin module for CBU1490. Interestingly, over half of these TA systems adopted the non-canonical gene orientation, each containing potential sigma 70 promoters in their cognate toxin coding sequence. Six TA modules were conserved in all *C. burnetii* suggesting they are important for *C. burnetii* pathogenesis. Both CBU0285 and CBUA0027 antitoxins are conserved throughout all *C. burnetii*, indicating that while their cognate toxin modules are missing in some strains, these antitoxin modules may be necessary for other functions outside of their role as an antitoxin. Consistent with this hypothesis was the ability of CBUA0027 to bind DNA in the absence of its cognate toxin (Fig 11) and reports showing antitoxins acting as transcription factors that control other regulons (Kim et al., 2010, Lin et al., 2013). Transposon mutants within toxin genes *cbu0007a* and *cbu1992* or *cbu0644* result in strong or mild replication defects in Vero cells suggesting that their loss or disruption of their cognate antitoxin expression is important for intracellular survival, while *himar1* mutant of *cbu1294* had no growth defects (Table 1) (Martinez et al., 2014, Newton et al., 2014). Furthermore, mutation of the *cbu0007*, *cbu0285*, or *cbu2084* antitoxins also display a strong replication defect in Vero cells presumably because of their cognate toxins (Martinez et al., 2014). The large number of TA systems in *C. burnetii* is unusual for a host-associated bacterium (Pandey and Gerdes, 2005) but fits the hypothesis that pathogenic bacteria have a larger number of TA systems (Georgiades and Raoult, 2011).

**Table 1:**
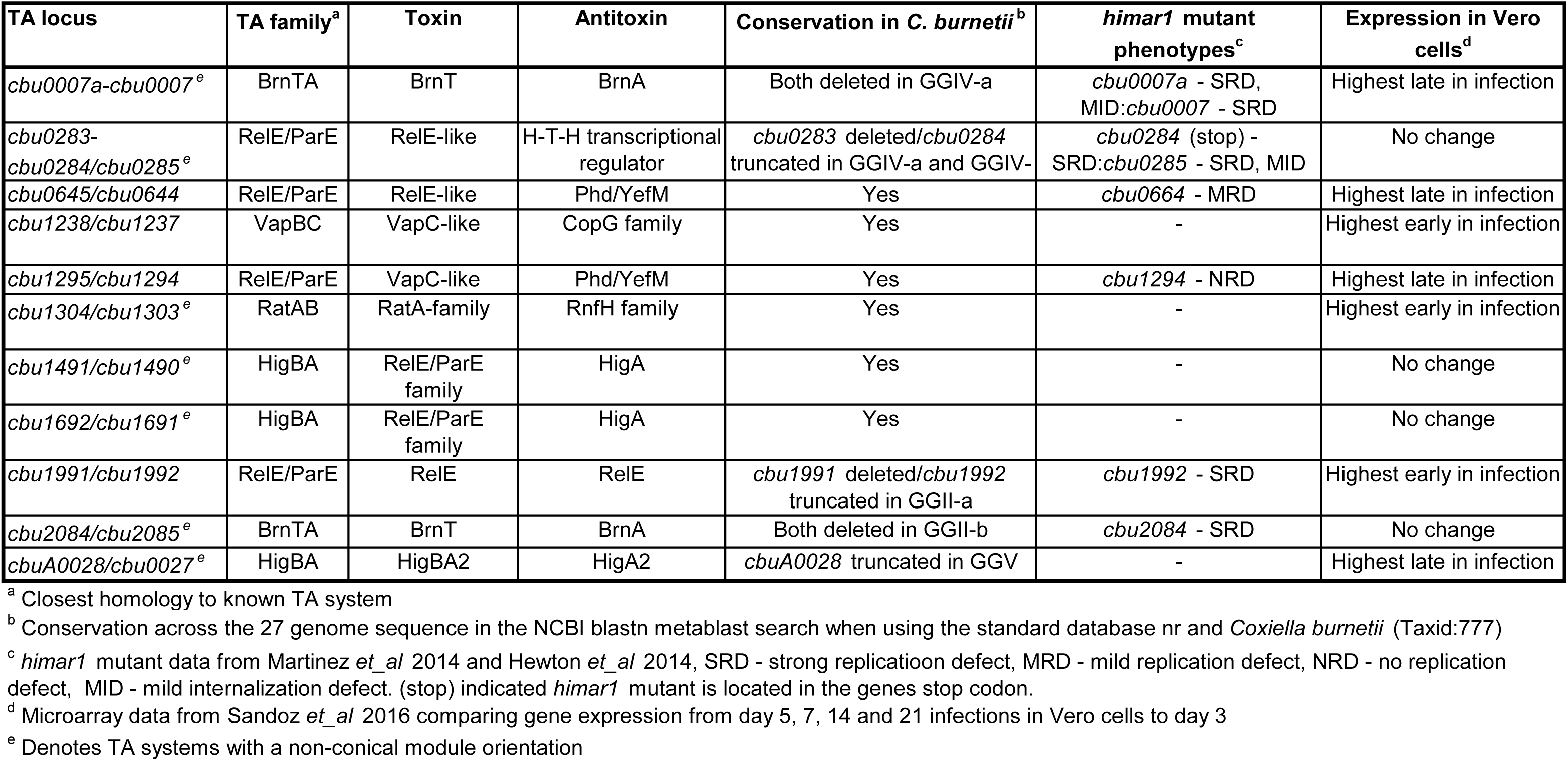
TA systems in *C. burnetii*.

Type II TA systems have a range of functions including maintenance of plasmids, virulence and pathogenicity and bacterial stress responses (Kamruzzaman et al., 2021). It is intriguing to speculate on their role in *C. burnetii* pathogenesis. CBUA0028/CBUA0027 have an obvious role in maintenance of the endogenous plasmid found in most *C. burnetii* strains. However, the absolute conservation of *cbuA0027*, including the conservation of its promoter internal to *cbuA0028*, and its ability to bind DNA suggests there are other roles for this TA system in *C. burnetii* survival. The *cbuA0028*/*cbuA0027* TA system is expressed highest in late stages of Vero cell infection (Table 1) (Sandoz et al., 2016) suggesting that it is important for late-stage infection and possibly for transition from LCV to SCV, which requires a large redefinition of protein content, metabolism, and cellular structure of the bacterium (Coleman et al., 2007, Coleman et al., 2004, Omsland et al., 2009, Sanchez et al., 2021). Similar roles for TA systems have been associated with reduced metabolism and persister cell formation upon stress conditions in other bacteria (Wang and Wood, 2011). Three other *C. burnetii* TA systems also have higher expression levels late in Vero cell infection (Table 1) (Sandoz et al., 2016) and may also aid in LCV to SCV transition or SCV persistence, while three TA systems have higher expression levels early in infection and may play a role in the SCV to LCV transition, including *cbu1238*/*cbu1237* which appears to be regulated by the *cbu0712* (*gacA*) regulator (data not shown, Wachter etal 2022). Four additional *C. burnetii* TA systems were expressed throughout the infection cycle (Table 1) (Sandoz et al., 2016) suggesting they may be important for responding to the harsh environmental conditions present in the *C. burnetii* CCV (Gutierrez and Colombo, 2005, Heinzen et al., 1999). Consistent with this hypothesis are findings that show TA systems also mediate several stress responses, including oxidative and nutritional stresses (Chan et al., 2018, Christensen et al., 2001, Christensen et al., 2003).

In summary, this study utilized a newly developed inducible CRISPRi system and two new nutritional selection systems to examine the role of the QpH1 plasmid and its gene content in *C. burnetii* pathogenesis. The data presented here identified and characterized the first TA system in *C. burnetii*, encoded by the QpH1 plasmid, and explains why most *C. burnetii* strains still contain an autonomously replicating plasmid. The CBUA0028/CBUA0027 TA system is one of 11 TA systems discovered in the *C. burnetii* genome and suggests an important role for TA systems in *Coxiella* pathogenesis.

## 4 Experimental procedures

### 4.1 Bacterial strains and mammalian cell lines

Bacterial strains used in this study are listed in Table S1. The *C. burnetii* Nine Mile phase II (RSA439, clone 4, NMII) strain was used for this work. NMII *C. burnetii* and genetic transformants were axenically cultured in ACCM-D as previously described (Sandoz et al., 2014), with the addition of 4-hydroxyphenylpyruvic acid (4-HPP) for tyrosine-based nutritional selection. For storage, bacteria were pelleted following 7 days of growth, washed 3 times in phosphate-buffered saline (PBS; 1 mM KH2PO4, 155 mM NaCl, 3 mM Na2HPO4, pH 7.4), and then suspended in freezing medium (ACCM-D containing 10% dimethyl sulfoxide) and frozen at −80°C. *E. coli* Stellar (Takara) and *E. coli* PIR2 (Invitrogen) cells were used for recombinant DNA procedures and cultivated in Luria-Bertani (LB) broth or terrific broth. THP-1 macrophages (TIB-202; ATCC) and African green monkey kidney (Vero) cells (CCL-81; ATCC) were maintained in RPMI 1640 medium containing 10% FBS at 37°C and 5% CO2. *C. burnetii* replication in host cells or in ACCM-D was measured by quantitative PCR of genome equivalents (GE) as previously described (Howe et al., 2010, Omsland et al., 2009) using a probe specific to *groEL*.

### 4.2 DNA amplification and plasmid construction

Plasmids used in this study are listed in Table S1. Oligonucleotides were purchased from Integrated DNA Technologies and are listed in Table S2. PCRs were carried out using AccuPrime Pfx or Taq (Invitrogen). PCRs purified using the Nucleospin gel and PCR clean-up kit were cloned into plasmid backbones digested with restriction enzymes (NEB) using the In-Fusion HD cloning system (Takara). In-Fusion reactions were transformed into *E. coli* Stellar cells (Takara) for non-R6K origin of replication-based vectors or PIR2 cells (Invitrogen) for R6K origin of replication plasmids. The sgRNA dsDNA fragments were cloned into plasmid backbones using the NEBridge Golden Gate Assembly kit (BsaI-HF ®v2) Construction of all the plasmids used in this study is detailed in File S1.

### 4.3 Transformation and selection of *C. burnetii* transformants

*C. burnetii* grown in ACCM-D was transformed by electroporation as described previously (Omsland et al., 2011). For transformation with pMiniTn7T or pB-CRISPRi vectors, bacteria were also co-electroporated with pTnS2::*1169^P^-tnsABCD* as previously described (Beare et al., 2011). Selection of tyrosine-complemented strains was achieved using ACCM-D minus tyrosine and additional 4-HPP. Selection of arginine-, lysine- and proline-complemented strains was carried out using ACCM-D minus arginine, lysine or proline, respectively. For arginine selection citrulline was also added.

### 4.4 Immunofluorescence microscopy

Vero cells were fixed with 4% paraformaldehyde in phosphate-buffered saline (PBS) for 30 min at room temperature and simultaneously permeabilized and blocked for 30 min with 0.1% Triton X-100 plus 1% BSA. Antibodies (3–5 μg/ml) were diluted in Triton buffer and samples stained for 30–60 min. Cells were stained for indirect immunofluorescence as previously described (Howe et al., 2003). Rabbit anti-*C. burnetii* serum and a mouse monoclonal antibody directed against the lysosomal marker CD63 (LAMP3) (clone H5C6, BD Biosciences) were used as primary antibodies. Alexa Fluor 488- and 594- IgG (Invitrogen) were used as secondary antibodies. Nuclei were stained with Hoescht 33342 (ThermoFisher). Coverslips were mounted using ProLong Gold (Thermo Fisher Scientific). Microscopy was conducted using a Zeiss LSM-710 confocal fluorescence microscope (Carl Zeiss). The area of CCVs was measured using CD63 as a CCV membrane marker. Unless otherwise stated, a minimum of 50 cells for each condition from 3 independent experiments was used for analyses. Fiji (Image J, National Institutes of Health) was used for all image analysis.

### 4.5 Induction and complementation of the CRISPRi system

Induction of dCas9-3x-cMyc and targeted sgRNA in the CRISPRi system was carried out using IPTG. Induction in axenic media and in host-cells was achieved using 0.1 mM and 2 mM IPTG, respectively. Induction of the CRISPRi complementation constructs was carried out using aTc. In axenic media and host-cells 5 ng/ml and 25 ng/ml of aTc was used to induce expression of the codon optimized complement constructs.

### 4.6 qRT-PCR analysis of CRISPRi knockdown

Bacterial cell culture pellets were lysed in 1.0 mL of Trizol (ThermoFisher Scientific, Waltham, MA). RNA containing aqueous phase was collected according to manufacturer’s recommended protocol except 1-Bromo-3-chloropropane was used instead of chloroform (MilliporeSigma, St. Louis, MO). RNA containing aqueous phase was combined with equal volume of RLT lysis buffer (Qiagen, Valencia, CA) and extracted using Qiagen AllPrep DNA/RNA 96-well system (Valencia, CA). The RNA quality was assessed using the Agilent 2100 Bioanalyzer using RNA 6000 Pico kit (Agilent Technologies, Santa Clara, CA). qRT-PCR probe and primer sets (Table S2) were designed using Primer Express version 3.0 (Life Technologies, Carlsbad, CA). qRT-PCR assay was performed using the AgPath-ID One-Step RT-PCR Buffer and Enzyme Mix (Life Technologies, Carlsbad, CA). AgPath-ID One Step RT PCR reactions were carried out in 20µl reactions with 1X RT-PCR Buffer, 1X RT-PCR Enzyme mix, 400 nM forward and reverse primers, and 120 nM of the fluorescent TaqMan probes. The *groEL* forward, reverse, and fluorescent oligo was combined with the oligos for each gene listed in the table. Oligos were purchased from Biosearch Technologies located in Novato, CA. The QPCR reactions were carried out at 50 °C for 10 minutes, 95 °C for 10 minutes, 55 cycles of 95 °C for 15 seconds and 60 °C for 45 seconds. Data was analyzed using ABI 7900HT version 2.4 sequence detection system software (Life Technologies, Carlsbad, CA). Normalized gene expressions were determined by comparative CT method (Thermofisher Scientific, Waltham, MA).

### 4.7 Immunoblotting

*C. burnetii* production of IcmD, IcmK, dCas9-3x-cMyc, GST-CBUA0027, CBUA0028-V5-6xHis and XpressT-6xHis-CBU0665 was examined by sodium dodecyl sulphate-polyacrylamide gel electrophoresis (SDS-PAGE) and immunoblotting. PVDF membrane was incubated with the following antibodies: polyclonal rabbit anti-IcmD antibody (1:5000), polyclonal rabbit anti-IcmK antibody (1:2500; generously provided by Edward Shaw, PCOM South Georgia), mouse monoclonal anti-c-myc antibody (1:5000; clone 9E10 BD Biosciences), goat polyclonal anti-GST antibody (1:5000, Sigma-Aldrich), mouse monoclonal anti-V5 antibody (1:5000; Invitrogen) and mouse monoclonal anti-XpressT antibody (1:5000; Invitrogen). Following incubation of membranes with primary antibody, reacting proteins were detected using anti-rabbit (IcmD and IcmK), anti-goat (GST-CBUA0027) or anti-mouse (dCas9-3x-c-myc, CBUA0028-V5-6xHis and XpressT-6xHis-CBU0665) IgG secondary antibodies conjugated to horseradish peroxidase (Pierce, Rockford, IL) and chemiluminescence using ECL Pico or Femto reagent (Pierce). Chemiluminescence was detected using the UVP ChemStudio plus imager (Analytik Jena) and images were processed using the VisionWorks software (Version 9.1.21054.7804).

### 4.8 Cell-free protein expression

T7-polymerase-based expression plasmids (pET28a(+)-*cbuA0028*, pDEST15-*cbuA0027* and pEXP1-cbu0665) were used as DNA template in cell-free *in vitro* transcription and translation (IVTT) reactions (100 µl) using the Expressway mini cell-free expression kit (Invitrogen) to achieve cell-free production of protein. The reactions were incubated at 30 °C with shaking at 300 rpm for 30 minutes, followed by the addition of feeding buffer and further incubation at 30 °C with shaking at 300 rpm for 5 hours. Lysates where then used in pulldown assays (see next section) or processed for SDS-PAGE analysis by resuspending in SDS-PAGE loading buffer and boiling for 10□min. Proteins were separated by SDS-PAGE on a 4-20% gradient gel and then immunoblotted as described above using antibodies directed against the GST tag (Sigma-Aldrich) to detect CBUA0027, V5 tag to detect CBUA0028 or the Xpress tag (Invitrogen) to detect CBU0665.

### 4.9 Pulldown assays

IVTT reaction mixtures (100 µl) were incubated at 4 °C for 1 h with a 50% slurry of PBS/Glutathione-Sepharose 4B (GE Healthcare) with gentle mixing. Glutathione-Sepharose was pelleted in Pierce spin cups (Thermo Fisher) by centrifugation at 1,000□×□g for 1□min and washed 8 times in Tris-buffered saline (pH 7.2) containing 1% Triton X-100. Proteins bound to the Glutathione-Sepharose beads were eluted using a 10 mM glutathione buffer (50 mM Tris, 10 mM reduced glutathione, pH 8.0). The final eluted protein mixes were then used in EMSA’s (see next section) or processed for SDS-PAGE analysis by resuspending in SDS-PAGE loading buffer and boiling for 10□min. Proteins were separated by SDS-PAGE on a 4-20% gradient gel and then immunoblotted as described above using antibodies directed against the GST tag (EMD Millipore) to detect CBUA0027, V5 tag to detect CBUA0028 or the Xpress tag (Invitrogen) to detect CBU0665.

### 4.10 Electrophoretic mobility shift assays

The upstream regions of the *cbuA0028*, *cbuA0029*, and groEL genes were first selected for PCR amplification. PCR products (1□μg) of desired templates were 3’ end-labeled using a Pierce biotin 3’ End DNA Labeling Kit (Thermo Scientific). The resulting probe reaction mixtures were electrophoresed on a 0.8% agarose gel for 30 min at 100 V and then gel purified with a NucleoSpin Gel and PCR Clean-up kit (Takara Bio USA). The EMSA binding reaction, consisting of 2.5% glycerol, 5 mM MgCl2, 50 mM KCl, 1 nM biotin-labeled DNA and varying concentrations of either CBUA0028 or CBUA0027-CBUA0028 in 1X Binding Buffer (LightShift Chemiluminescence kit; Thermo Scientific), were assembled and incubated at room temperature for 30 min. A nondenaturing loading dye (0.25% bromophenol blue) was added, and the resulting RNA mixtures were resolved on a 10% polyacrylamide gel for 2 h at 100 V. DNA/protein complexes were transferred to a Hybond-N+ positively charged nylon membrane (Amersham Pharmacia Biotech) using an electroblot transfer system (Bio-Rad) and cross-linked with short-wave UV light in a GS gene linker UV chamber (Bio-Rad). A North2South chemiluminescence hybridization and detection kit (Thermo Scientific) was used to detect resulting bands. The blot was imaged on a UVP ChemStudio PLUS Imager (Analytik Jena).

### 4.11 Promoter-mScarlet-i fluorescence reporter

Individual wells of a 12-well tissue culture plate containing 2 ml of ACCM-D minus lysine were inoculated with *C. burnetii* harboring promoter-mScarlet-i constructs at a cell density of 2 × 10^6^ GE/ml. After 7 or 14 days of incubation, the growth medium was thoroughly mixed by pipetting and 200 μl of each culture was added to a black Cellstar 96-well microplate (Greiner Bio-One). Fluorescence was measured using a Cytation 5 plate reader (Agilent).

### 4.12 Computer analysis

Geneious Prime 2021.1.1 (https://www.geneious.com) was used to analyze *C. burnetii* genome sequences for TA systems and to conduct sigma 70 motif searches. DNA alignments were carried out using the Clustal Omega multiple sequence alignment software (https://www.ebi.ac.uk/Tools/msa/clustalo/). Structural-based homology searches were carried out using the I-Tasser program (https://zhanggroup.org/I-TASSER/).

### 4.13 Statistical analysis

Statistical analyses were performed using a one-way analysis of variance (ANOVA) or unpaired student *t* test and Prism software (Version 9.3.1; GraphPad Software, Inc., La Jolla, CA).

---

Figure S1 – Inducible knockdown of IcmD production by CRISPR interference and complementation using codon optimized *icmD*. (a) Sequences of wild-type (*icmD*) and codon optimized (*icmD-CO*) icmD. Sequences targeting *icmD* target A and target B are denoted in blue and red, respectively. Silent mutations made to the *icmD-CO* nucleotide sequence are denoted in bold nucleotides. PAM sequences are underlined. (b) Schematic of the aTc-inducible expression of *icmD*-target A-CO. The orientation of the *tetR* repressor, P*tetA* promoters and *icmD*-target A-CO codon optimized gene are depicted. (c) Immunoblots of *icmD* CRISPRi strains containing the pJB-TetRA-icmD-CO complementation plasmid in the absence of CRISPRi inducer (IPTG) or complement inducer (aTc). Specific protein detection was achieved using anti-IcmK, anti-cmyc (dCas9) and anti-IcmD antibodies. Expression of codon optimized Target A *icmD* restored production of IcmD in the *icmD* CRISPRi Target A strain but was unable to restore IcmD production in the *icmD* CRISPRi Target B strain.

Figure S2 – Dual CRISPRi repression of IcmD and ScvA translation. (a) Schematic of the IPTG-inducible double CRISPRi expression construct (from pB-CRISPRi-*icmD*-sgRNA-1-*scvA*-sgRNA-1). (b) Immunoblots of the *icmD*/*scvA* double CRISPRi knockdown strain in the absence or presence of IPTG. Specific protein detection was achieved using anti-c-Myc (dCas9), anti-IcmD, anti-ScvA, and anti-IcmK, antibodies. Induction of CRISPRi resulted in dual knockdown of both IcmD and ScvA production.

Figure S3 – CRISPRi knockdown of QpH1 ORFs in axenic media. Individual *C. burnetii* CRISPRi strains targeting one of two separate regions of each ORF of QpH1 were grown ACCM-D in the absence (CRISPRi off) or presence of 0.1 mM IPTG (CRISPRi on) for 6 days. Replication is shown as fold increase in *C. burnetii* genome equivalents (GE) of CRISPRi strains. Results are expressed as the means of results from two biological replicates from three independent experiments. Error bars indicate the standard deviations from the means, and asterisks indicate a statistically significant difference (Student’s *t* test, * - P < 0.05, ** - P < 0.01 *** - P < 0.001, and **** - P < 0.0001) compared to values for the CRISPRi off samples.

Figure S4 – CRISPRi knockdown of QpH1 ORFs in THP-1 macrophages. Individual *C. burnetii* CRISPRi strains targeting one of three separate regions of each ORF of QpH1 were grown in the absence (CRISPRi off) or presence of 2 mM IPTG (CRISPRi on) for 5 days. Replication is shown as fold increase in *C. burnetii* genome equivalents (GE) of CRISPRi strains. Results are expressed as the means of results from two biological replicates from three independent experiments. Error bars indicate the standard deviations from the means, and asterisks indicate a statistically significant difference (Student’s *t* test, * - P < 0.05, ** - P < 0.01 *** - P < 0.001, and **** - P < 0.0001) compared to values for the CRISPRi off samples.

Figure S5 – Detection of dCas9 induction in *C. burnetii* QpH1 CRISPRi strains. Lysates of *C. burnetii* QpH1 CRISPRi strains grown for 7 days in ACCM-D with or without 0.1 mM IPTG were analyzed by immunoblot. Production of dCas9 was detected using an anti-c-myc antibody. The dCas9 protein was detected in all induced strains except for the *cbuA0027* CRISPRi strains. CRISPRi knockdown of *cbuA0027* causes severe growth defects in axenic media, hence no dCas9 protein was detected.

Figure S6 – QPCR analysis of CRISPRi knockdown of QpH1 Dot/Icm substrate genes. RNA was extracted from QpH1 Dot/Icm substrate strains grown in ACCM-D for 6 days in the absence (CRISPRi off) or presence of IPTG (CRISPRi on). Samples for QPCR analysis were normalized using the comparative CT method. For each RNA sample a gene specific or control *groEL* probe and primer set was used to examine the level of expression.

Figure S7 – QPCR analysis confirmed CRISPRi knockdown *cbuA0027*. RNA was extracted from *cbuA0027*-84 CRISPRi strain grown in ACCM-D for 6 days in the absence (CRISPRi off) or presence of IPTG (CRISPRi on). Samples for QPCR analysis were normalized using the comparative CT method. For each RNA sample a *cbuA0027* specific or control *groEL* probe and rimer set was used to examine the level of expression.

Figure S8 – Alignment of *cbuA0028* and *cbuA0027* from NMII and G Q212. The *cbuA0028* (a) and *cbuA0027* (b) genes from NMII and G Q212 strains were aligned using clustal omega. The G Q212 genes are labeled *cbuA0028*G and *cbuA0027*G. The putative promoter (-35 and -10 regions) for *cbuA0028* are denoted and the sequence underlined. The start codon for *cbuA0028* and *cbuA0028*G are colored red and blue, respectively. Asterisks demark conserved nucleotides.

Figure S9 – Production of CBUA0027, CBUA0028 and CBU0665 by cell-free IVTT. Cell-free IVTT was carried out using the following combinations of template DNA (250 ng each): pDEST15-*cbuA0027* (GST-A27), pET28a(+)-*cbuA0028* (A28-V5), pEXP1-*cbu0665* (XpressT-665), GST-A27/A28-V5 or GST-A27/XpressT-665. Lysates from cell-free IVTT were analyzed by immunoblots probed (a) simultaneously with anti-V5 and anti-XpressT to detect CBUA0028 and CBU0665, respectively, or with (b) anti-GST to detect CBUA0027.

Figure S10 – EMSA shows the CBUA0027/CBUA0028 complex does not bind the *cbuA0029* or *groEL* promoters. EMSAs show interactions between biotin-labeled *cbuA0028* (P*cbuA0028*) or *cbuA0029* (P*cbuA0029*) promoters and increasing concentrations of purified CBUA0027/CBUA0028 complex. Biotin-labeled P*groEL* with CBUA0028/CBUA0027 (640 nM) was included as negative control. The location of bound and unbound probe is depicted by arrows.

Figure S11 - Alignment of the *cbuA0028*-*cbuA0027* operon from NMII and G Q212. The *cbuA0028*-*cbuA0027* operon from NMII and G Q212 strains were aligned using clustal omega. The G Q212 operon is labeled *cbuA0028*G/*cbuA0027*G. The putative sigma 70 promoters (-35 and -10 regions) are denoted, and the sequence underlined. The start codon for *cbuA0028* and *cbuA0028G* are colored red and blue, respectively. Bold italicized sequence denotes the *cbuA0027* and *cbuA0027G* start codons. The *cbuA0027* P2 and P3 promoter sequences are colored green and grey, respectively. Asterisks demark conserved nucleotides. Only the *cbuA0027* P3 promoter is conserved in both strains.

Figure S12 – Off-targeting of *cbuA0021*-sgRNAs. (a) Alignments of the CRISPRi targeted region of *cbuA0021* with *cbu0334* (*thiDE*) and *cbu0055* (*ubiA*). The location of the three CRISPRi target regions in demarked with arrows. Sequences colored red denote conserved regions within the 20 bp CRISPRi targets. (b) CRISRPi repression by *cbuA0021*-sgRNA-61 (CBUA0021-61) and *cbuA0021*-sgRNA-63 (CBUA0021-63) causes a growth defect in axenic media, whereas only *cbuA0021*-sgRNA-62 (CBUA0021-62) causes a growth defect in THP-1 macrophages. Replication is shown as fold increase in *C. burnetii* genome equivalents (GE) of CRISPRi strains targeting *cbuA0021* after 6 days in ACCM-D or after a 5-day infection of THP-1 macrophages in the absence (CRISPRi off) or presence (CRISPRi on) of CRISPRi system inducer molecule IPTG. Replication of a no sgRNA target was examined as a negative control. CRISPRi repression of the essential *icmD* Dot/Icm apparatus gene was used as a positive control in THP-1 macrophages. Results are expressed as the means of results from two biological replicates from three independent experiments. Error bars indicate the standard deviations from the means, and asterisks indicate a statistically significant difference (Student’s t test, ** - P < 0.01, and *** - P < 0.001) compared to values for the CRISPRi off samples. (c) QPCR analysis confirmed CRISPRi knockdown *cbuA0021* and shows off-targeting knockdown on *cbuA0334*. RNA was extracted from *cbuA0021*-61 CRISPRi strain grown in ACCM-D for 6 days in the absence (CRISPRi off) or presence of IPTG (CRISPRi on). Samples for QPCR analysis were normalized using the comparative CT method. For each RNA sample a *cbuA0021*- and *cbu0334*-specific or control *groEL* probe and primer set was used to examine the level of expression.

Figure S13 – Eleven TA systems were identified in *C. burnetii*. Schematic of the 11 TA modules in *C. burnetii* indicating their gene orientation. Black arrows indicate putative promoter regions in non-canonical TA modules for driving increased expression of the antitoxin module. Red arrows depict the location of 5′ truncations in other *C. burnetii* strains. Dark blue arrows represent *himar1* insertions that result in strong replication defects in Vero cells. The light blue arrow depicts a *himar1* mutant with a mild replication defect. The green arrow represents the location of a *himar1* mutant that has normal replication. Key: antitoxin and toxin genes are colored light and dark grey, respectively.

## Acknowledgements

We thank Rose Perry-Gottschalk of the Research Technologies Branch, National Institute of Allergy and Infectious Diseases for graphical support. This work was supported by funding from the Intramural Research Program of the NIH, NIAID.

## Author contributions

Conceptualization: PAB; Methodology and investigation: SW, DCC, HEM, KV, BD, KK, PAB; Data analysis: PAB, SW; Supervision: RAH; Writing original draft: PAB; Review and editing: all authors.

