## Supplemental Figures S1-S13 for "A toxin-antitoxin system ensures plasmid stability in *Coxiella burnetii*"

Supplemental figure 1

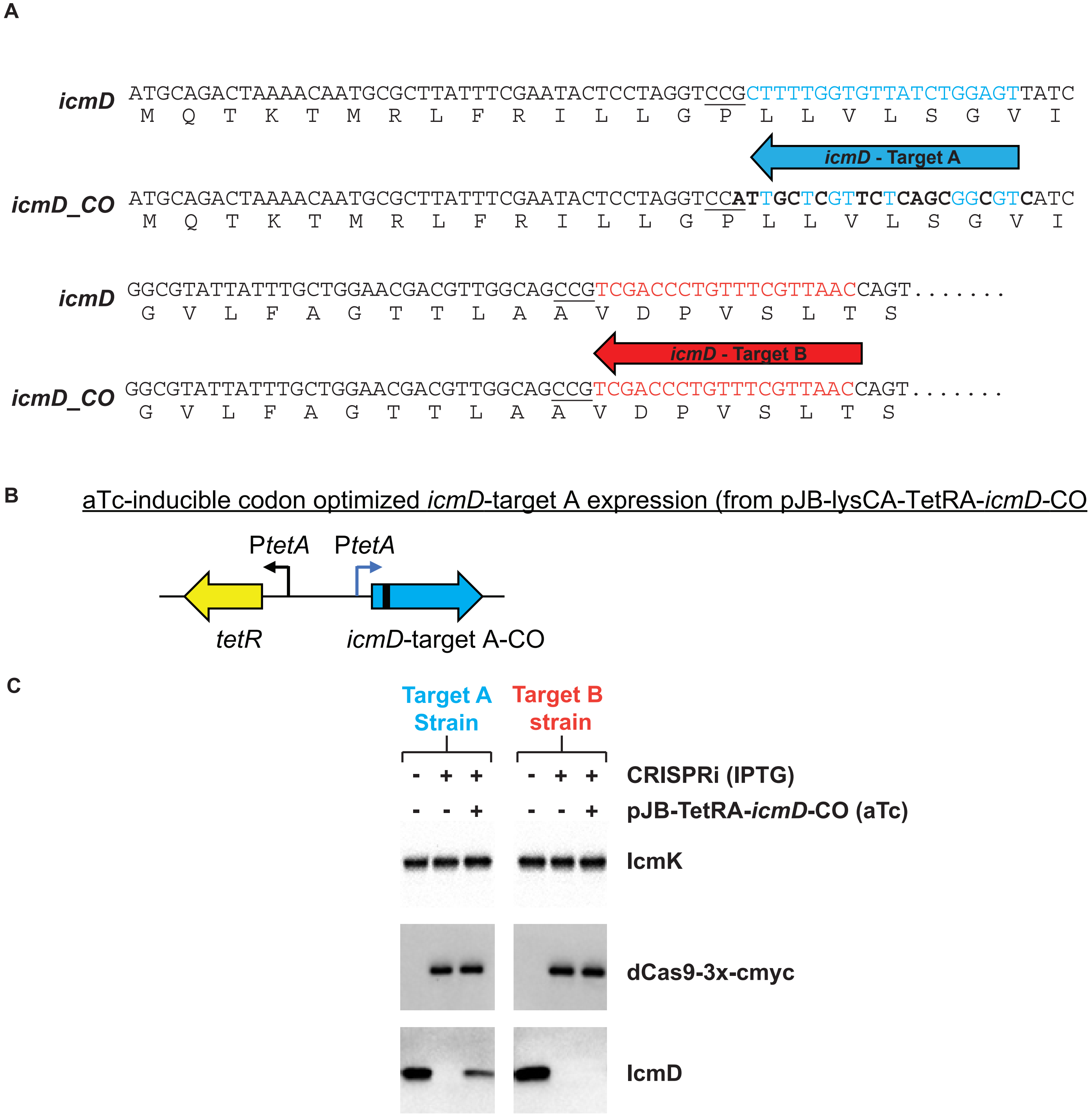

Supplemental figure 2

**A**

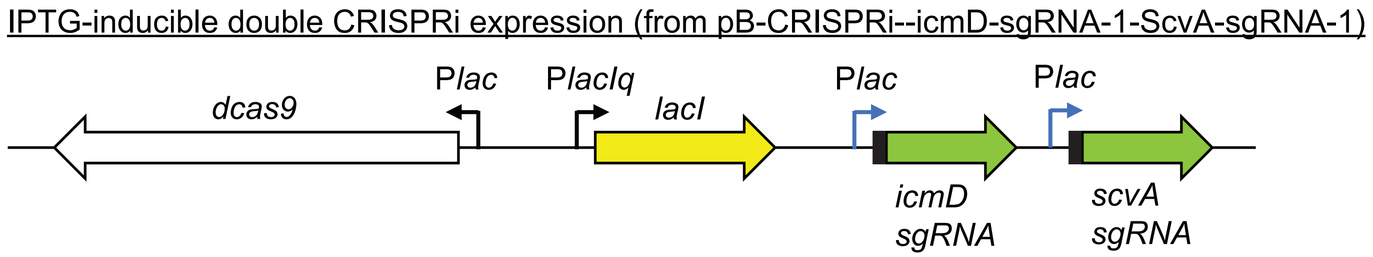


**B**

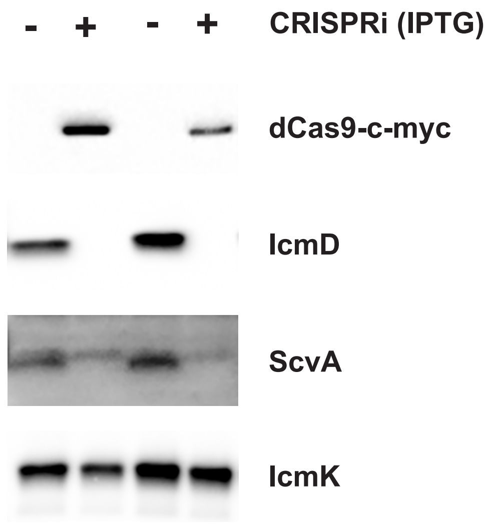

Supplemental figure 2

**A**

IPTG-inducible double CRISPRi expression (from pB-CRISPRi--icmD-sgRNA-1-ScvA-sgRNA-1)

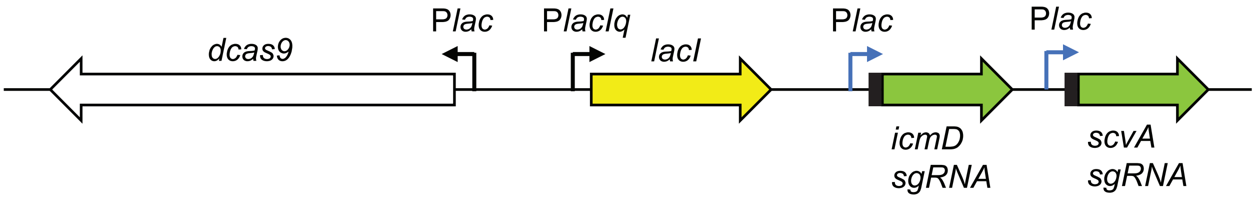


**B**

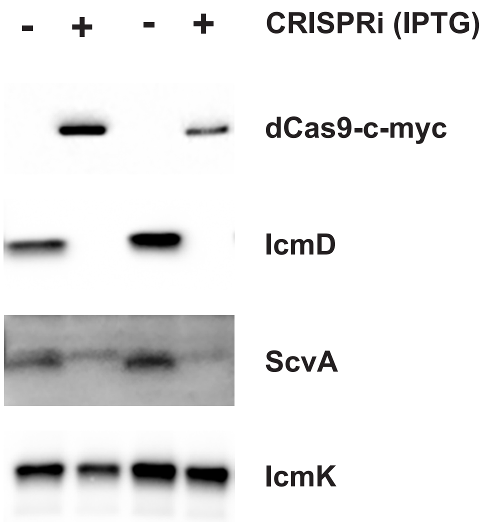

Supplemental figure 3

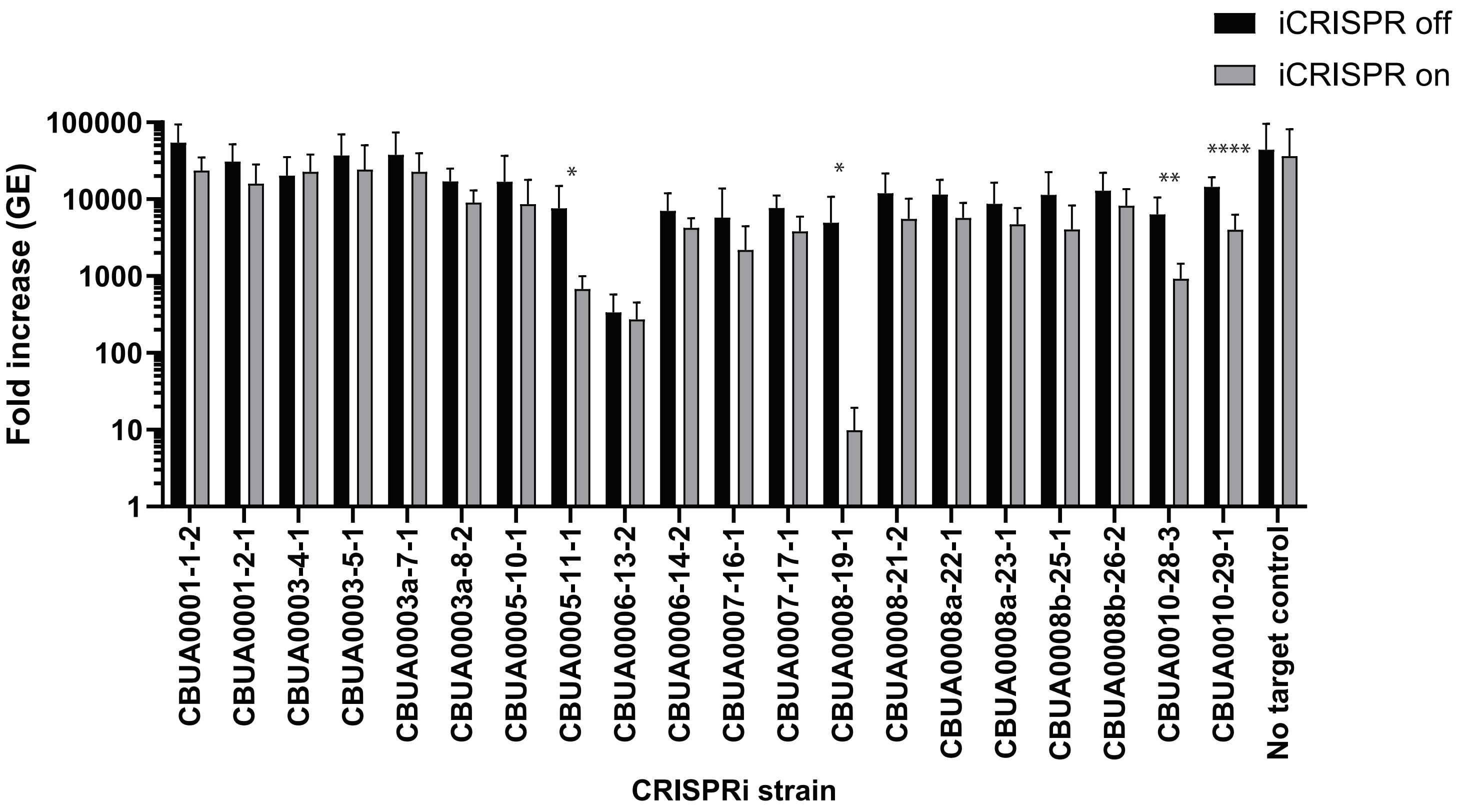


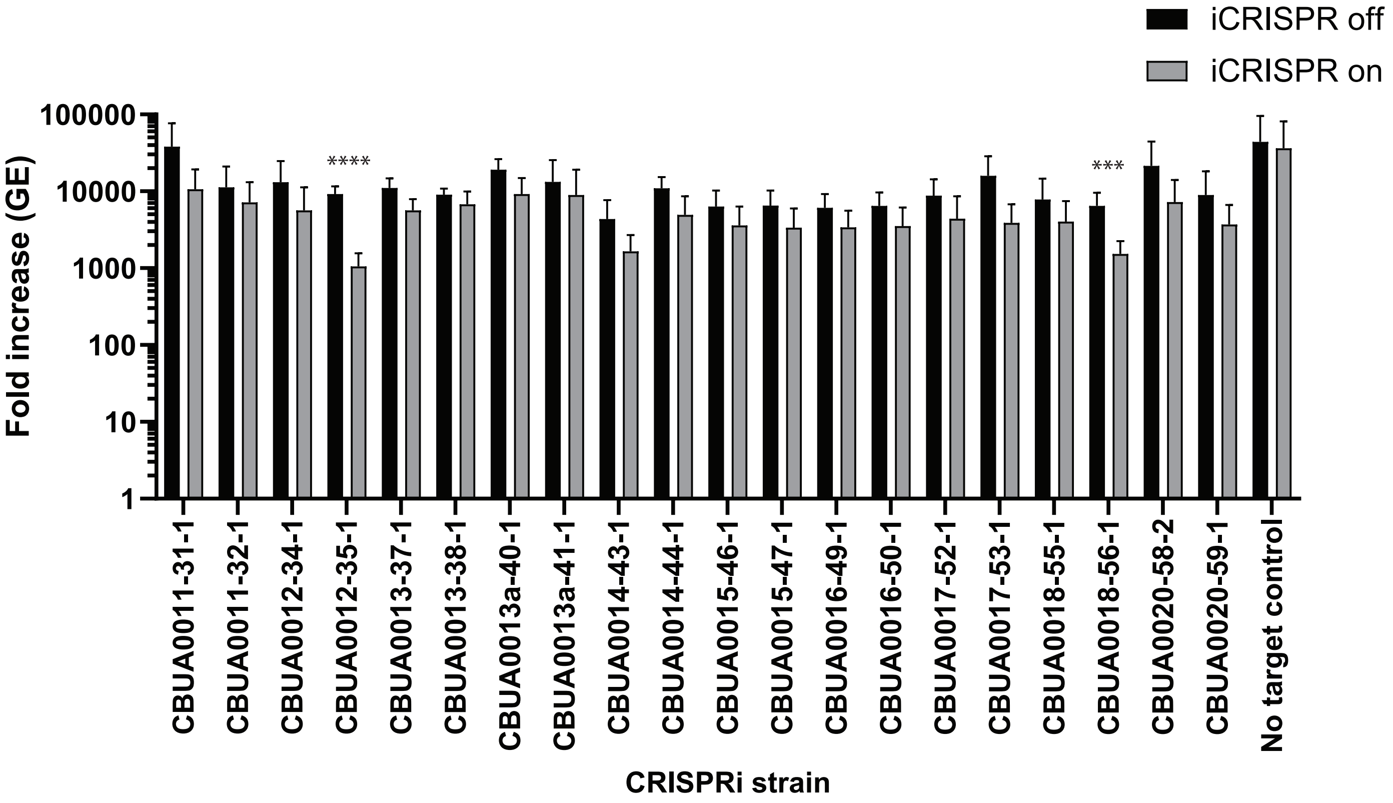

Supplemental figure 3

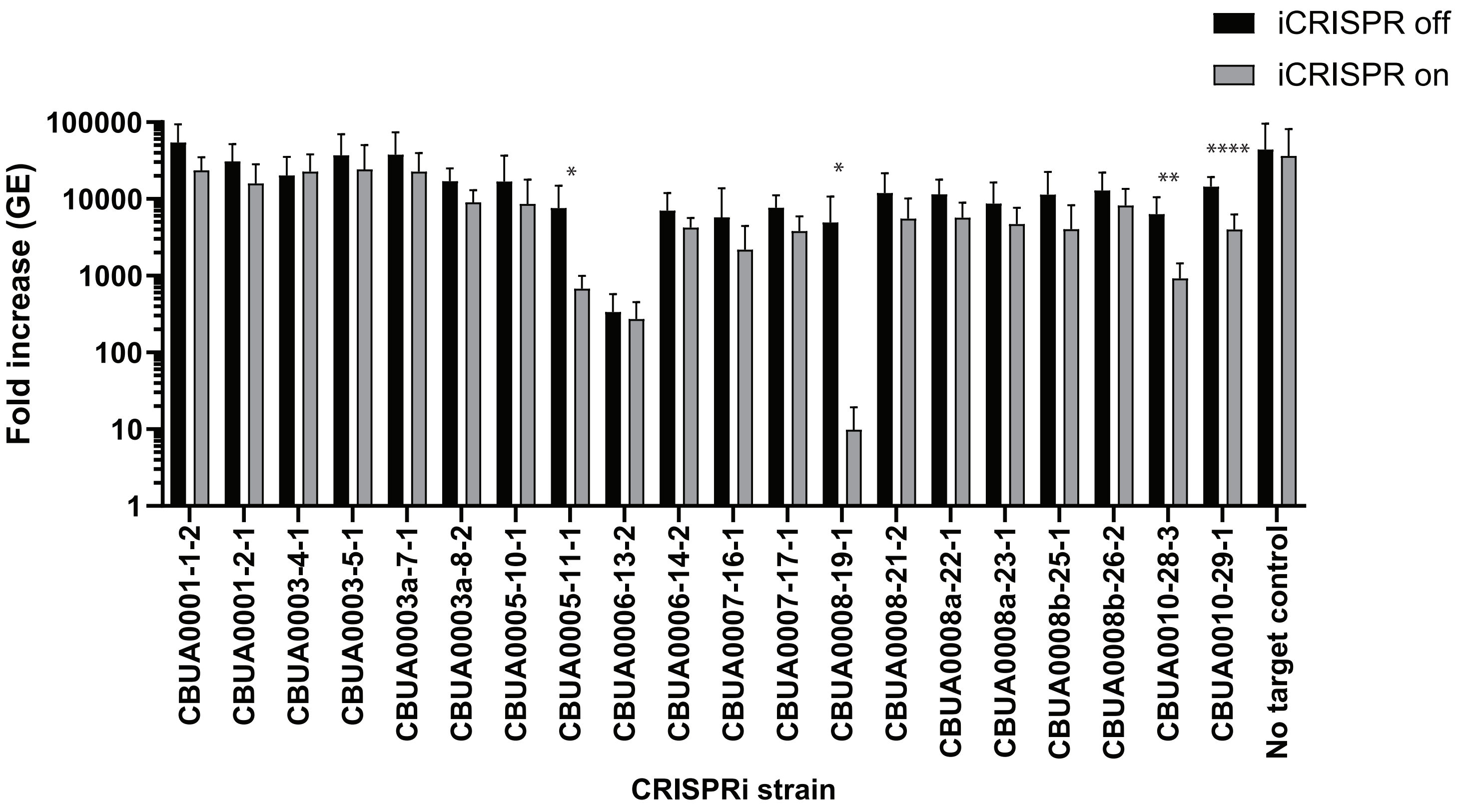


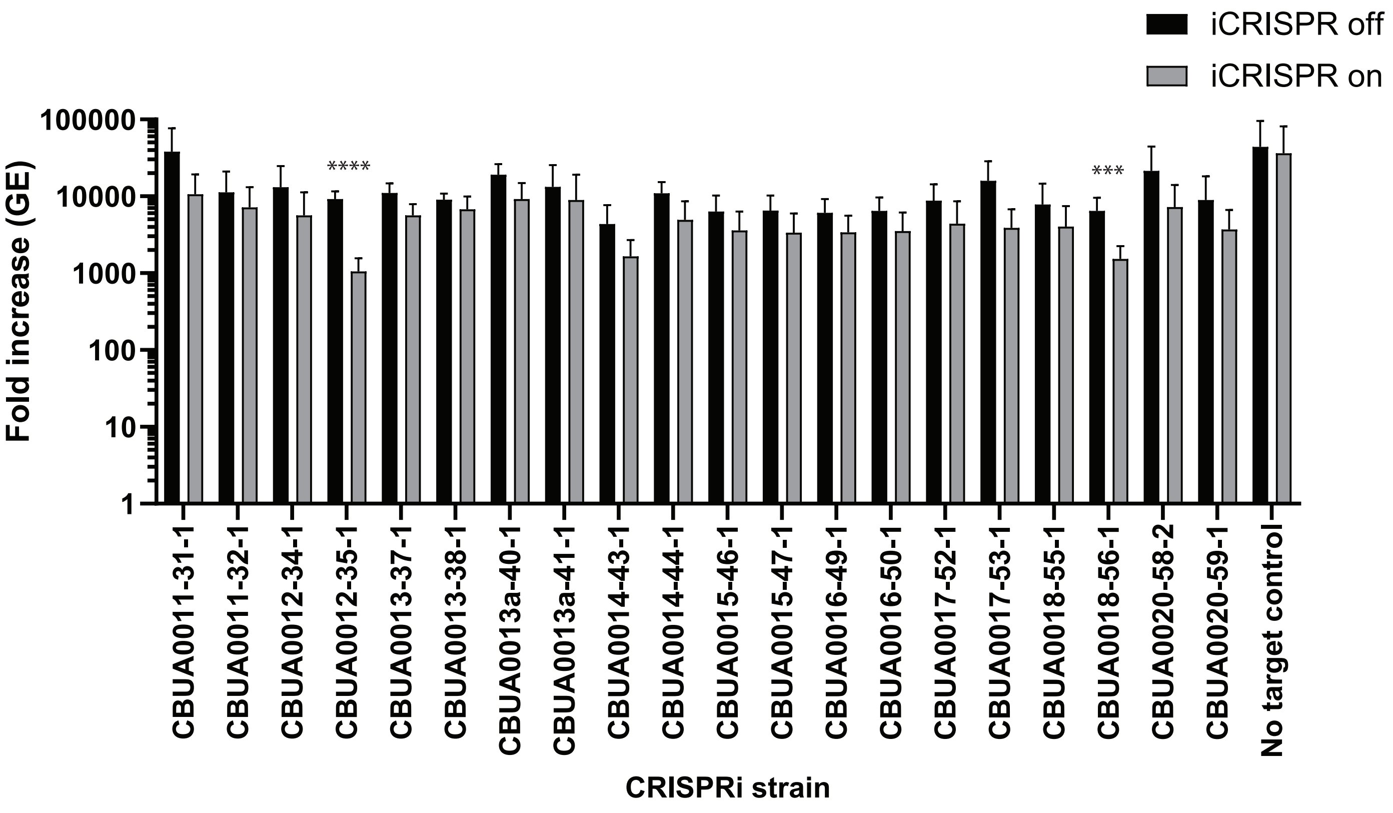

Supplemental figure 3

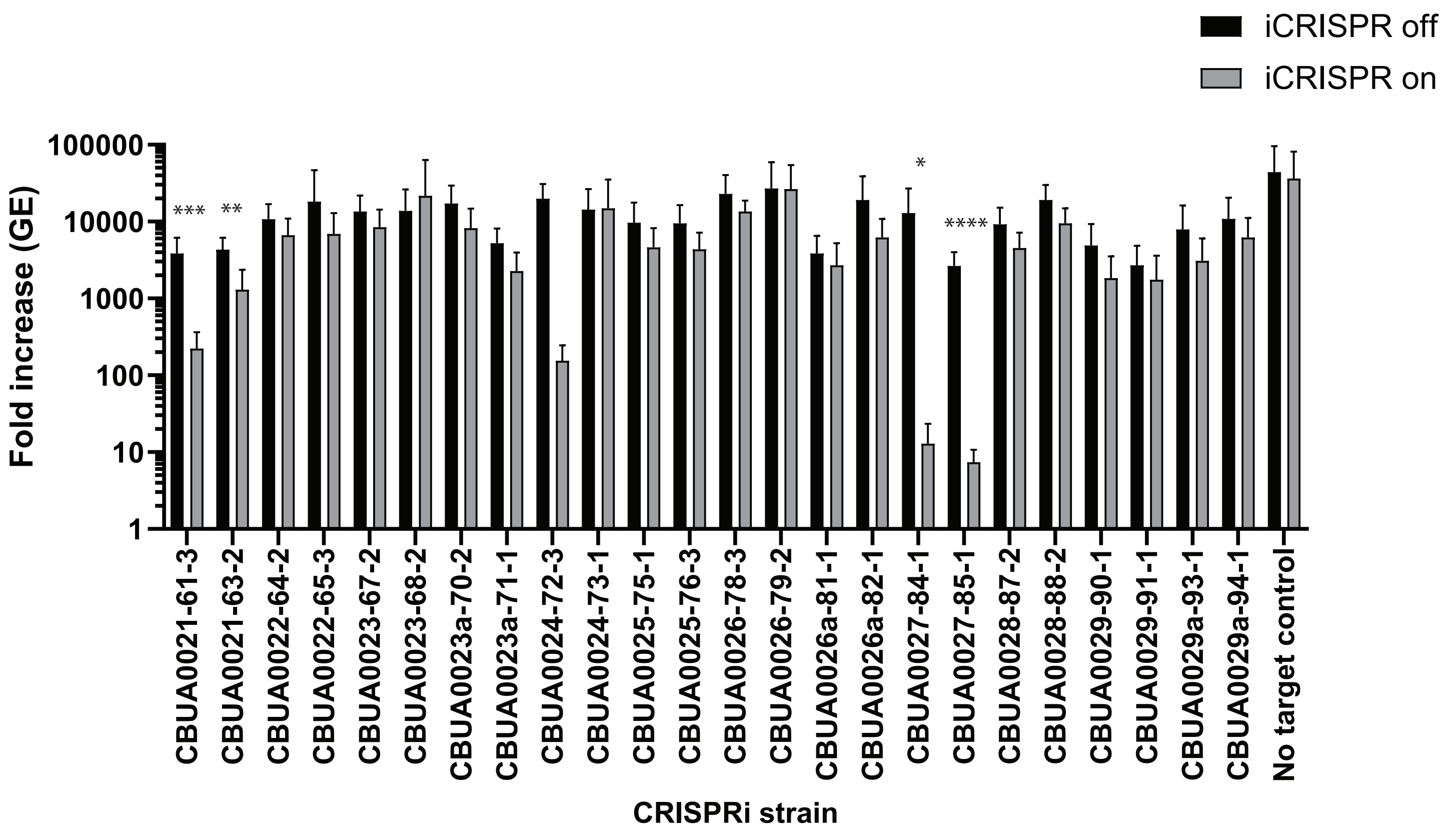


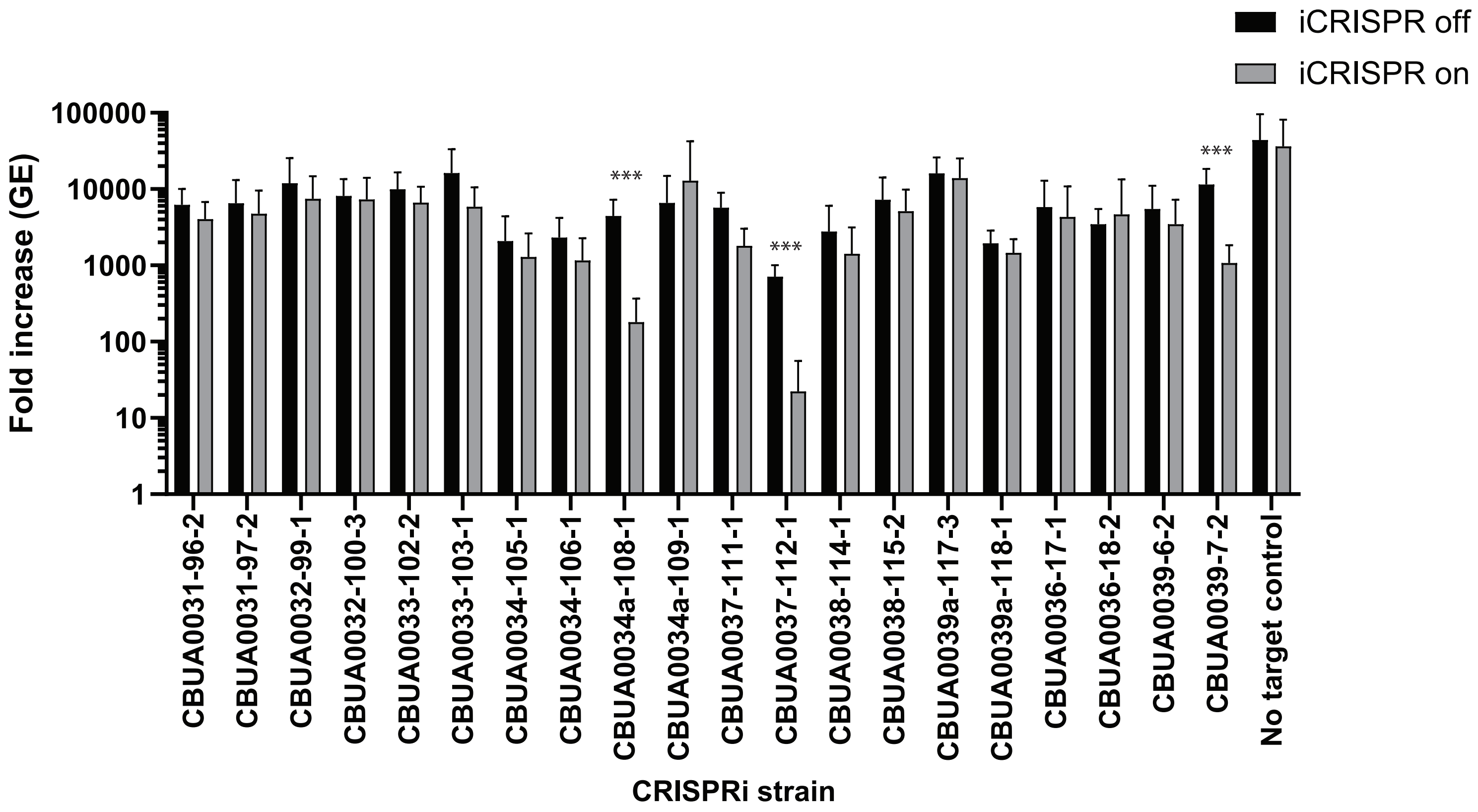

Supplemental figure 3

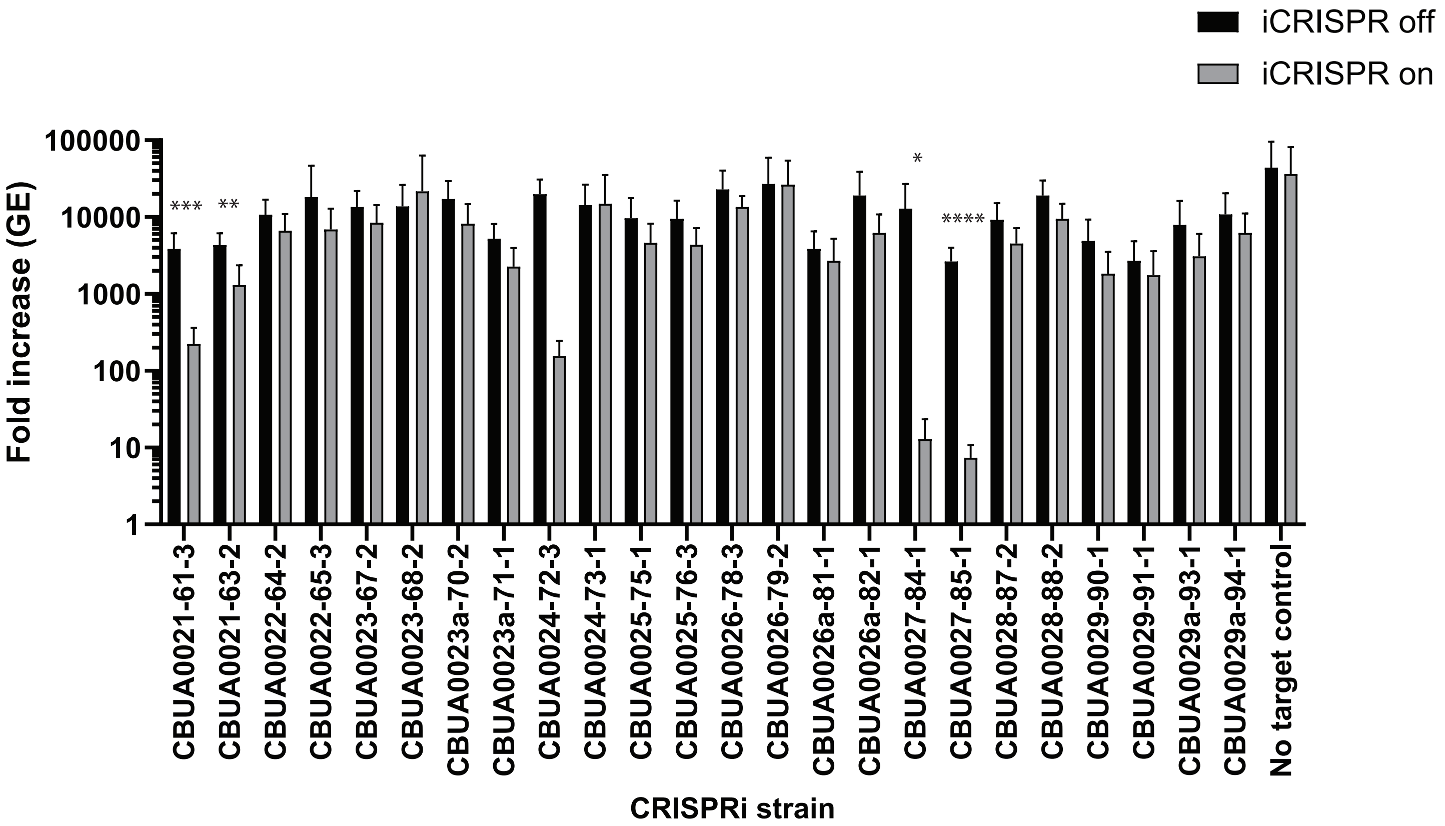


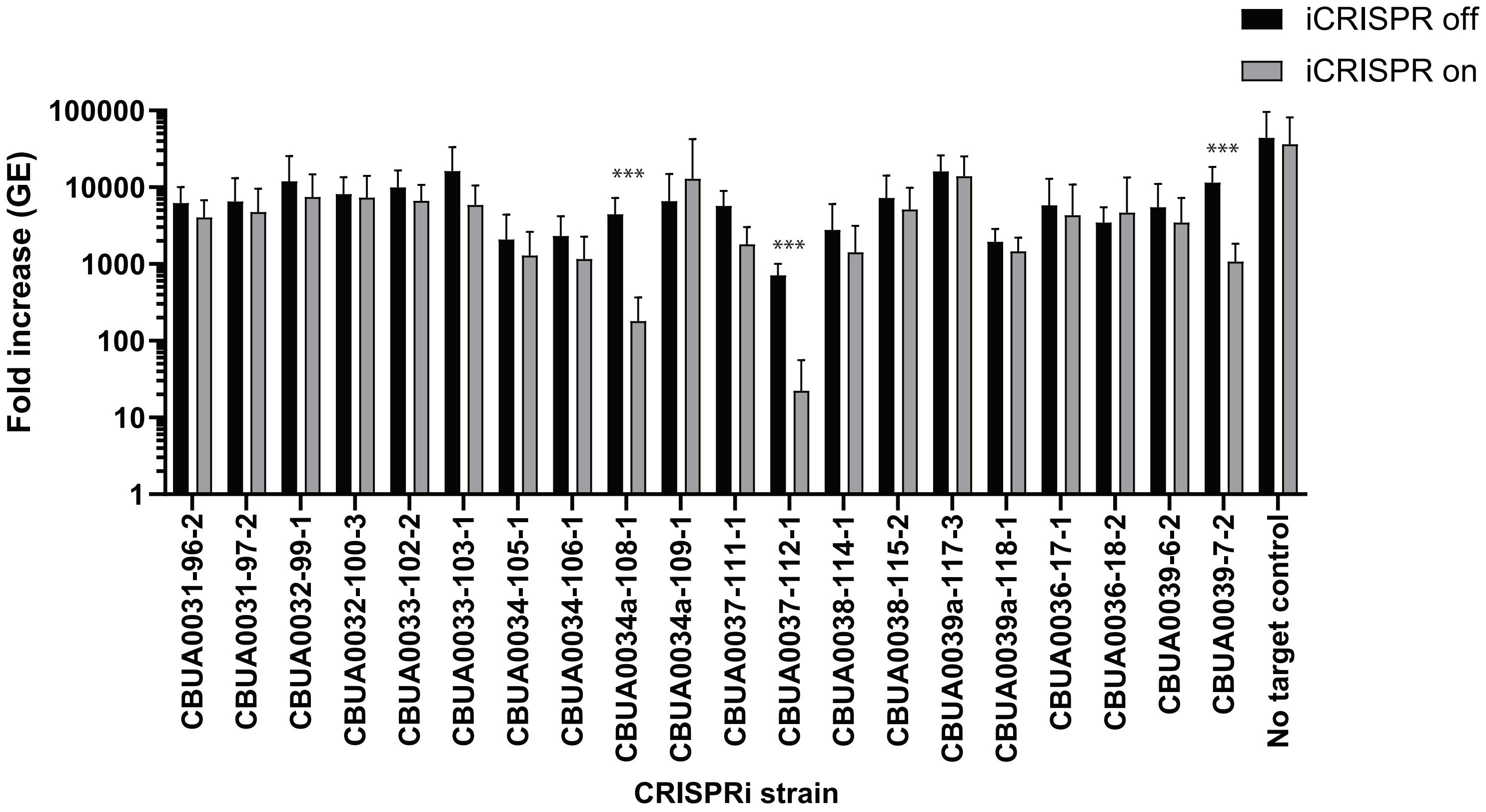

Supplemental figure 4

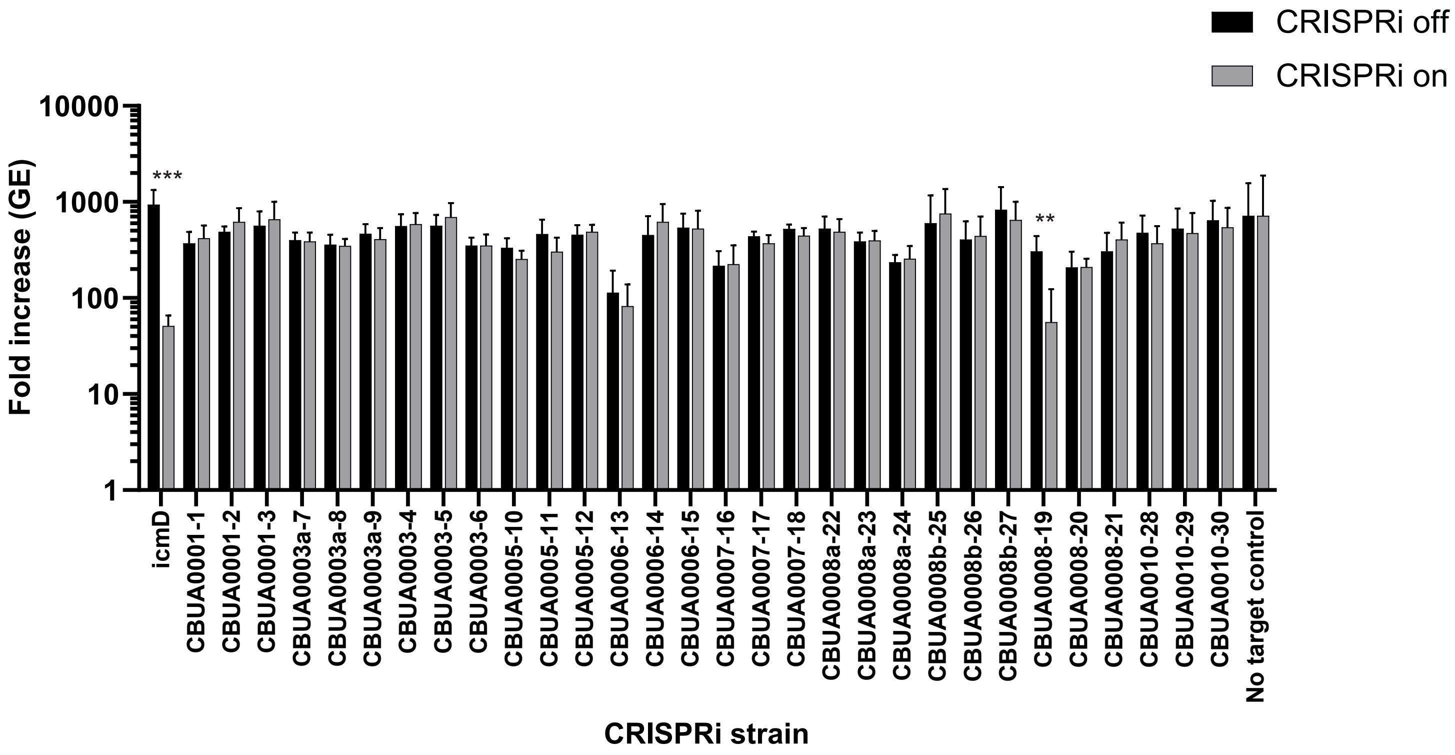


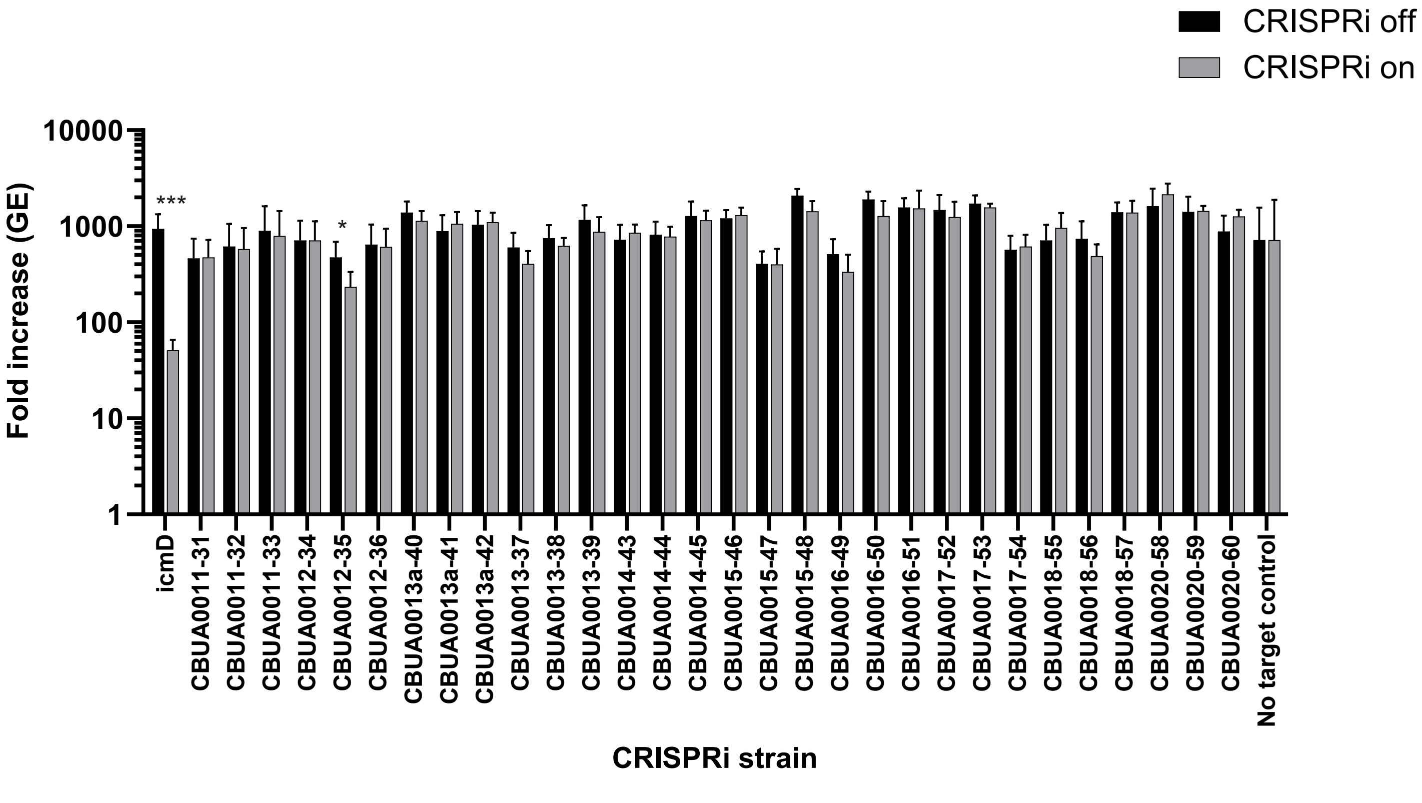

Supplemental figure 4

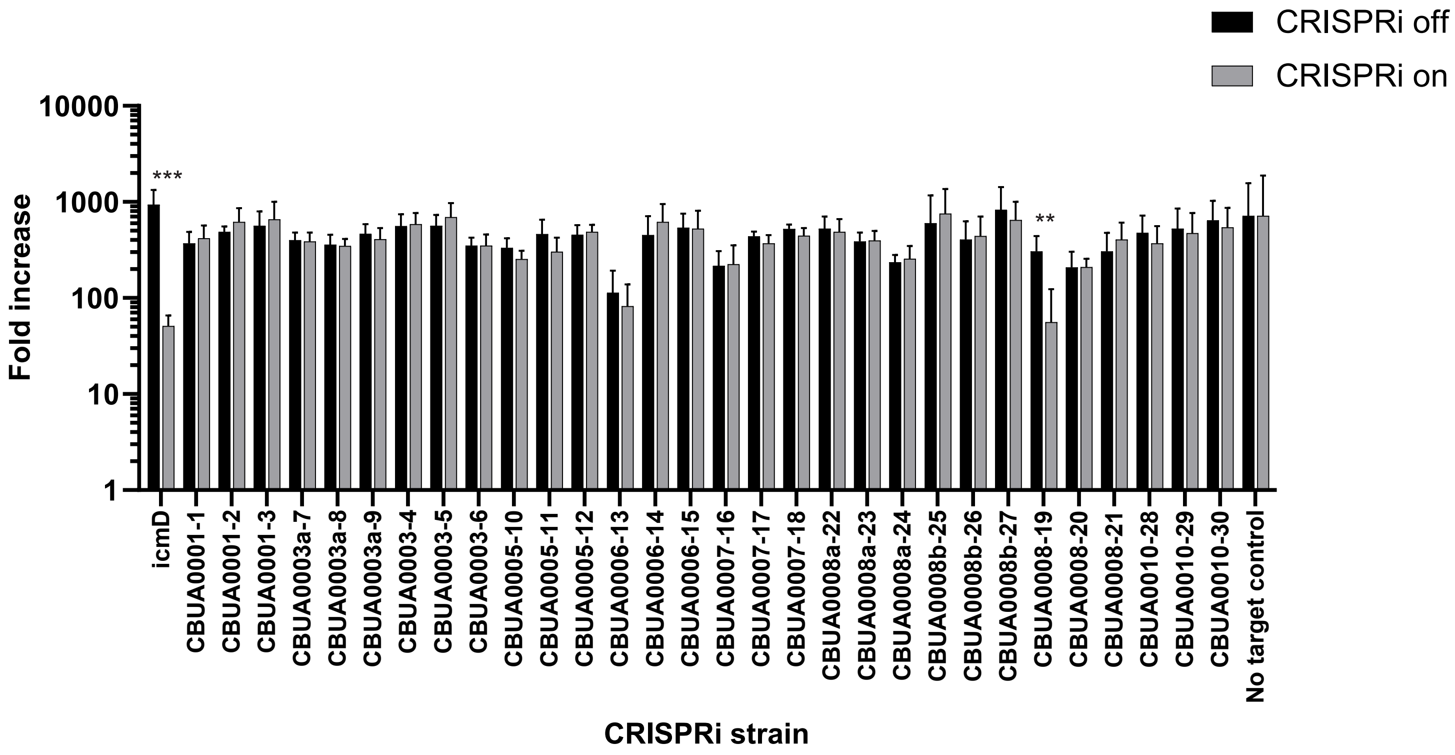


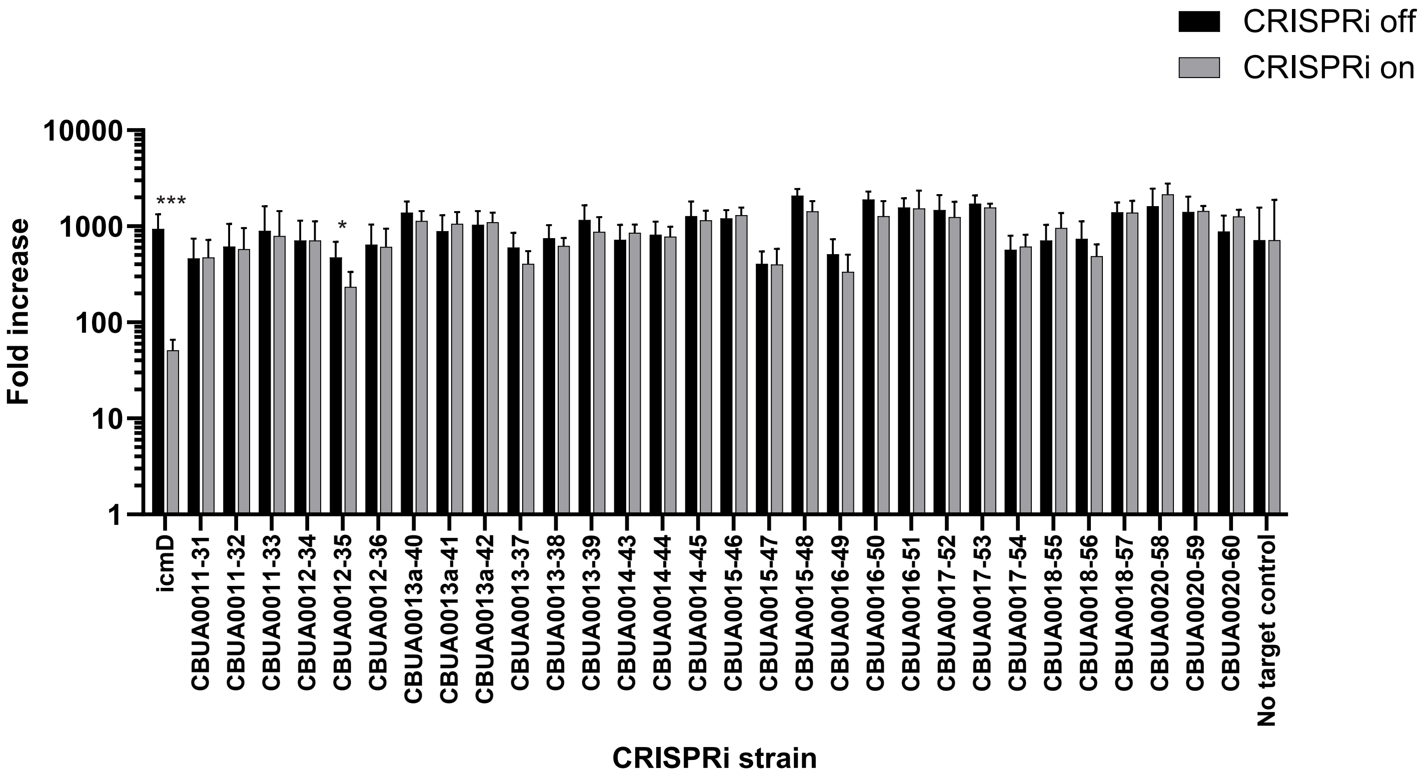

Supplemental figure 4

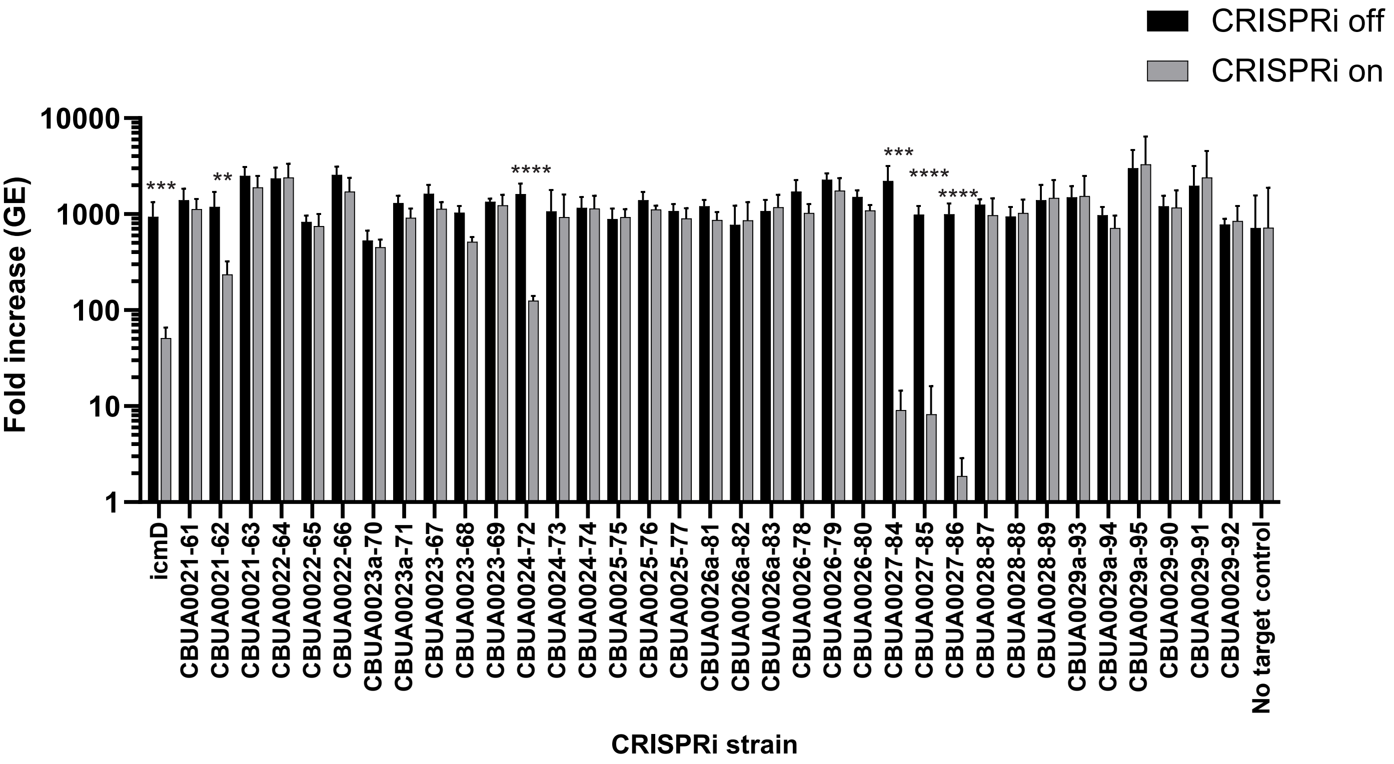


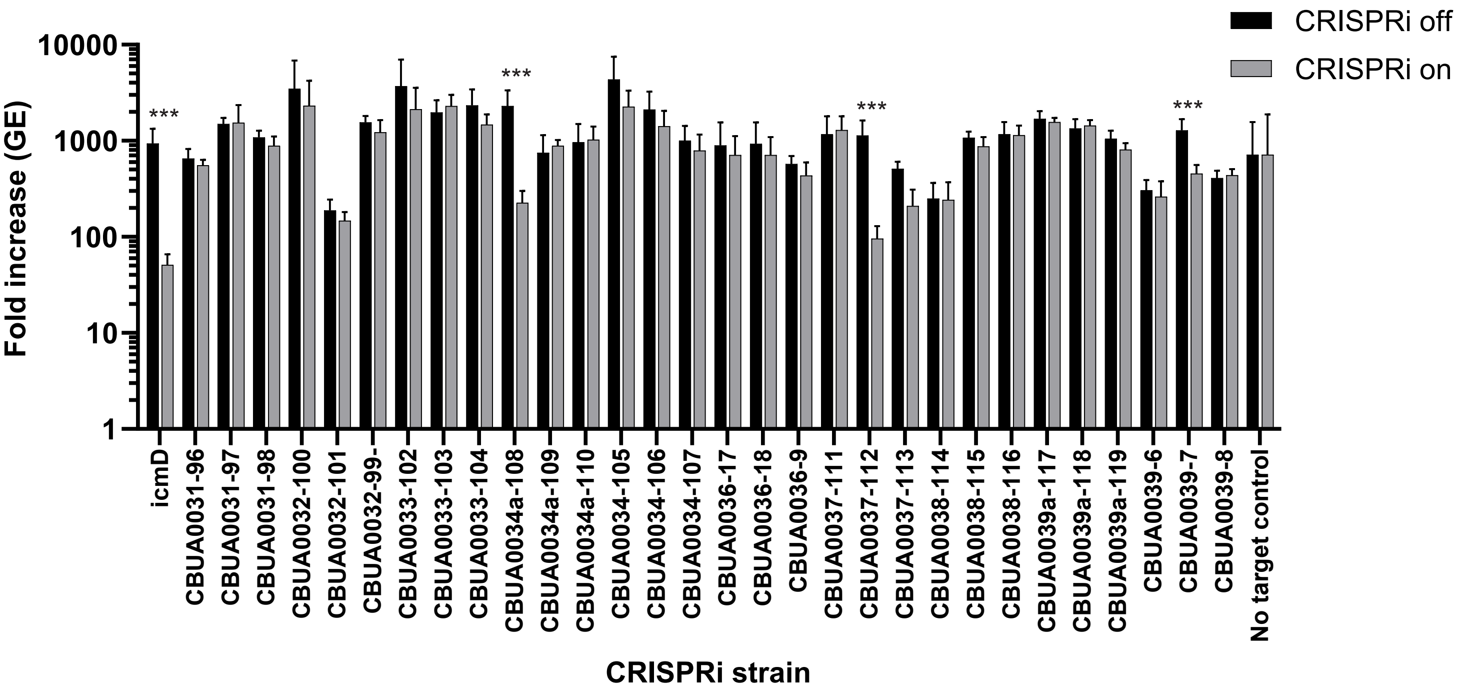

Supplemental figure 4

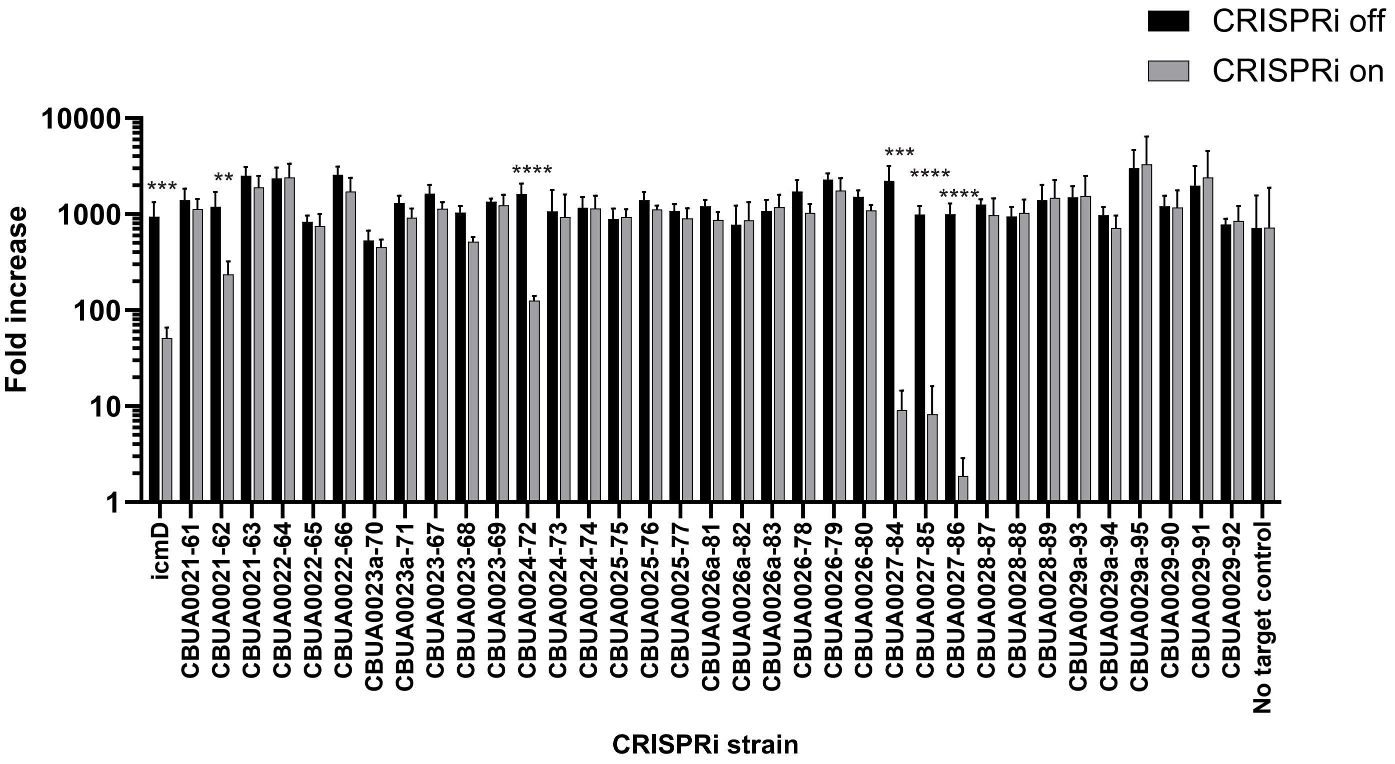


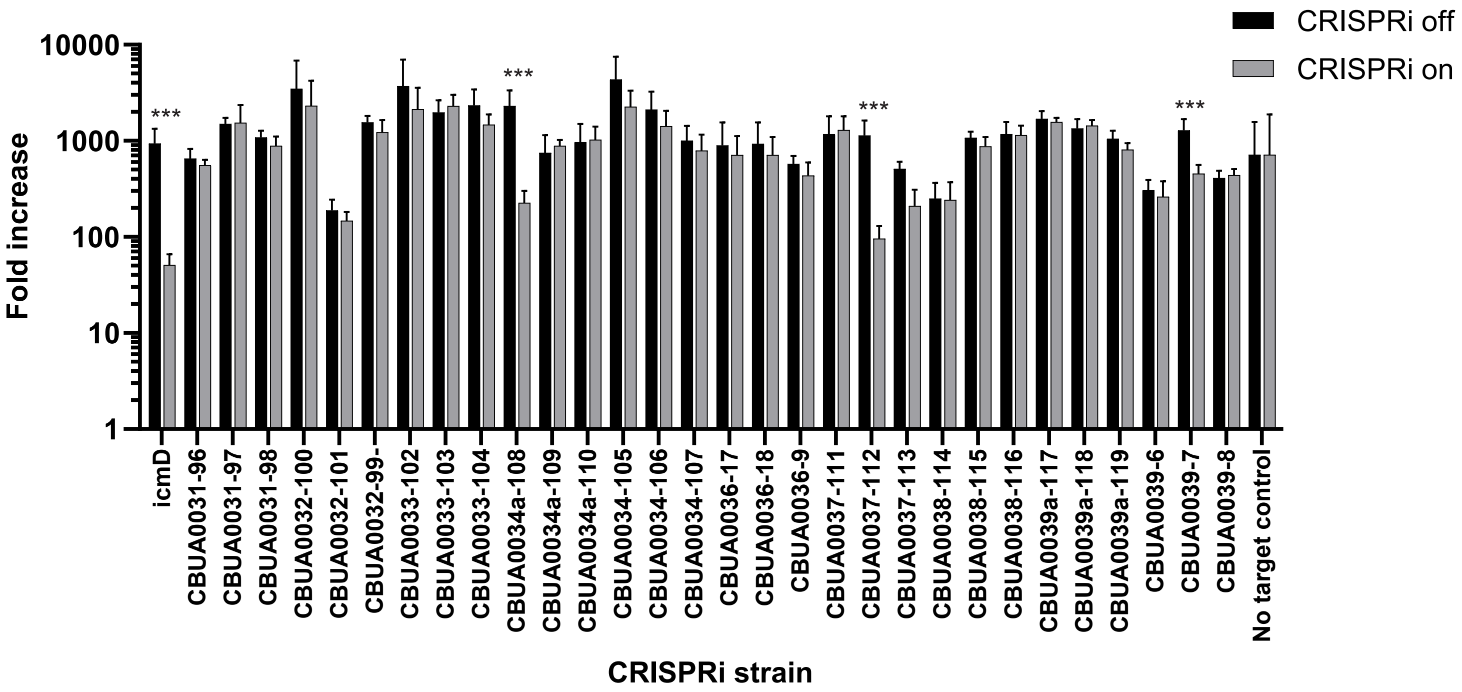

Supplemental figure 5

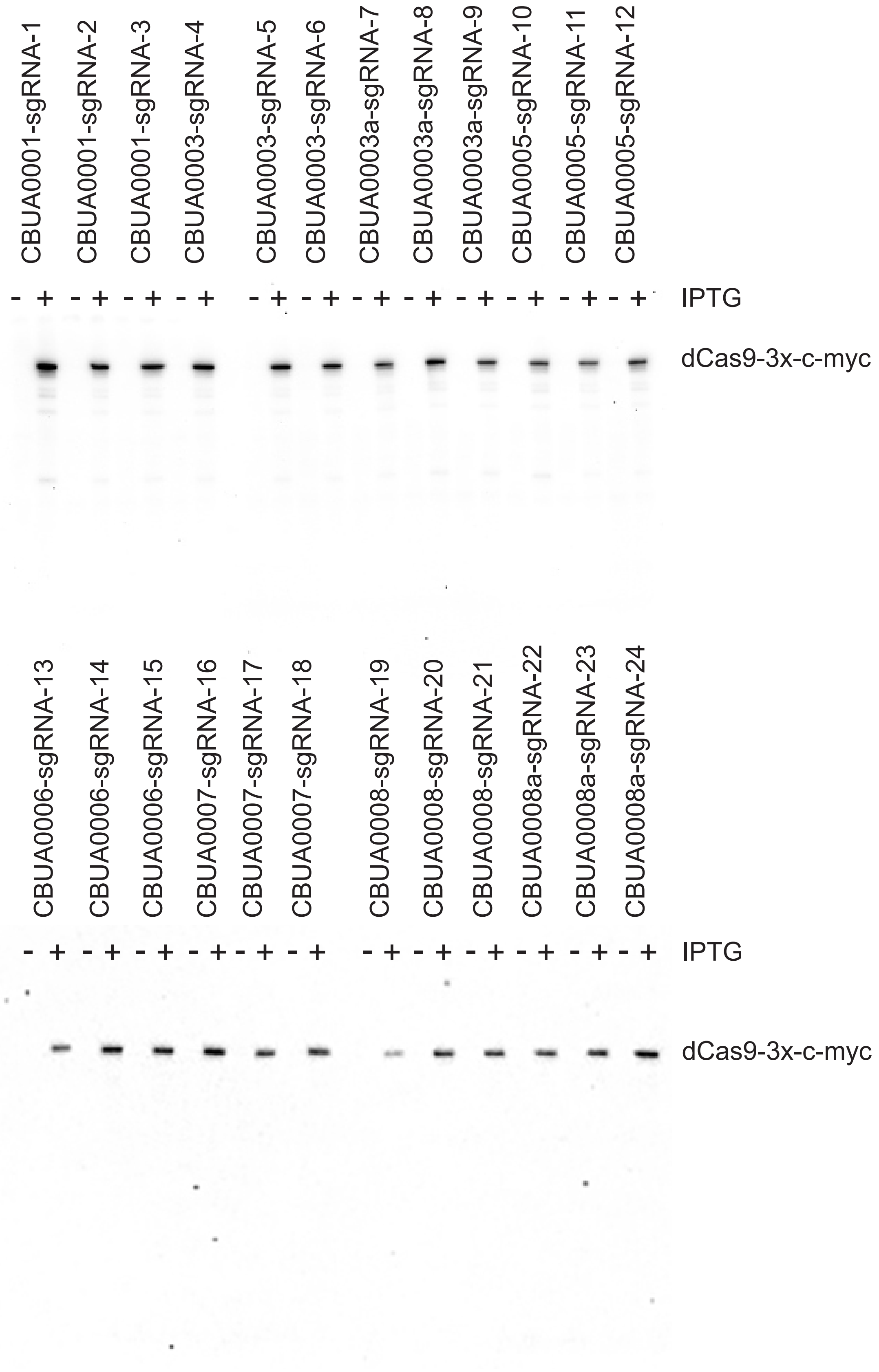

Supplemental figure 5

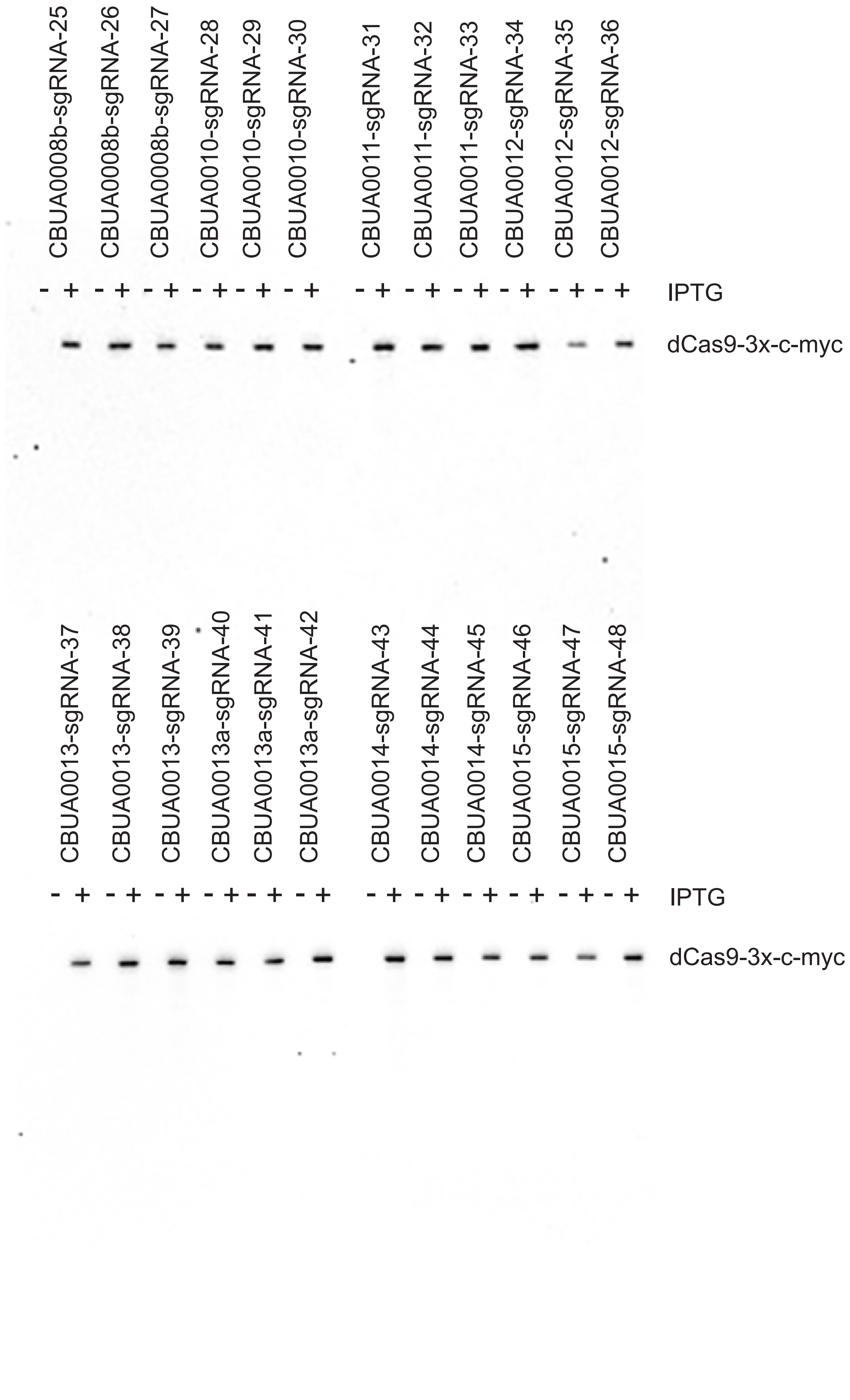

Supplemental figure 5

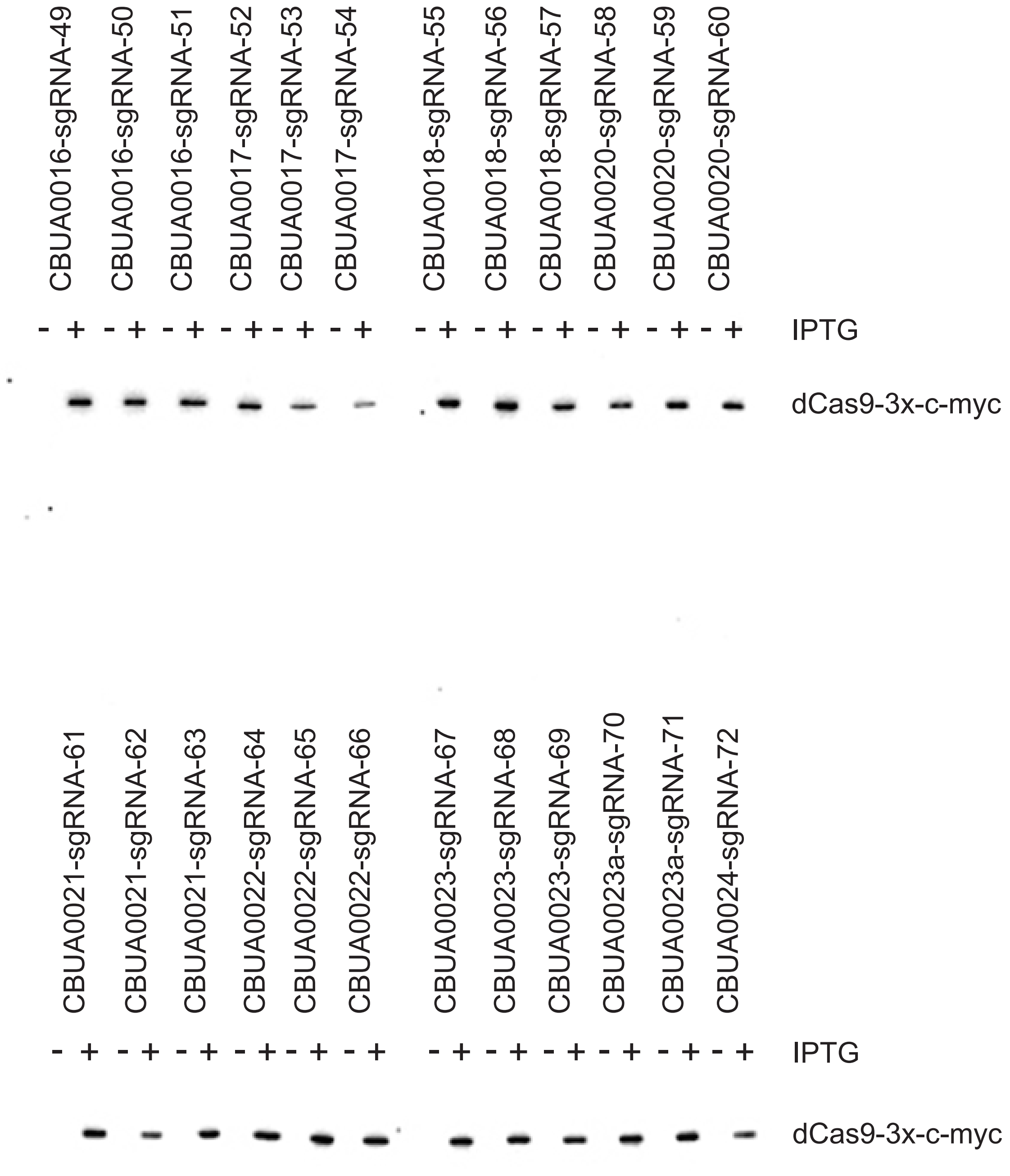

Supplemental figure 5

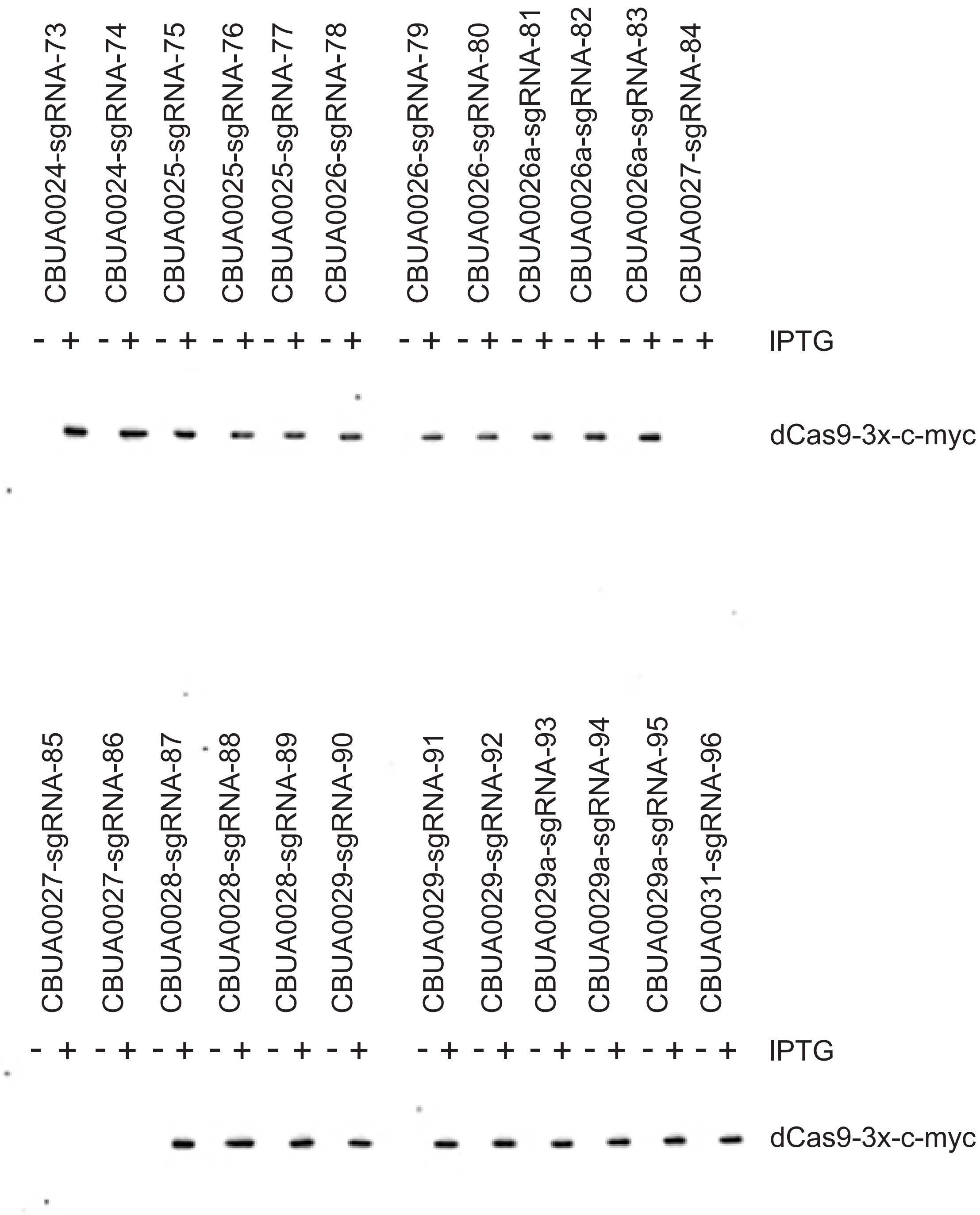

Supplemental figure 5

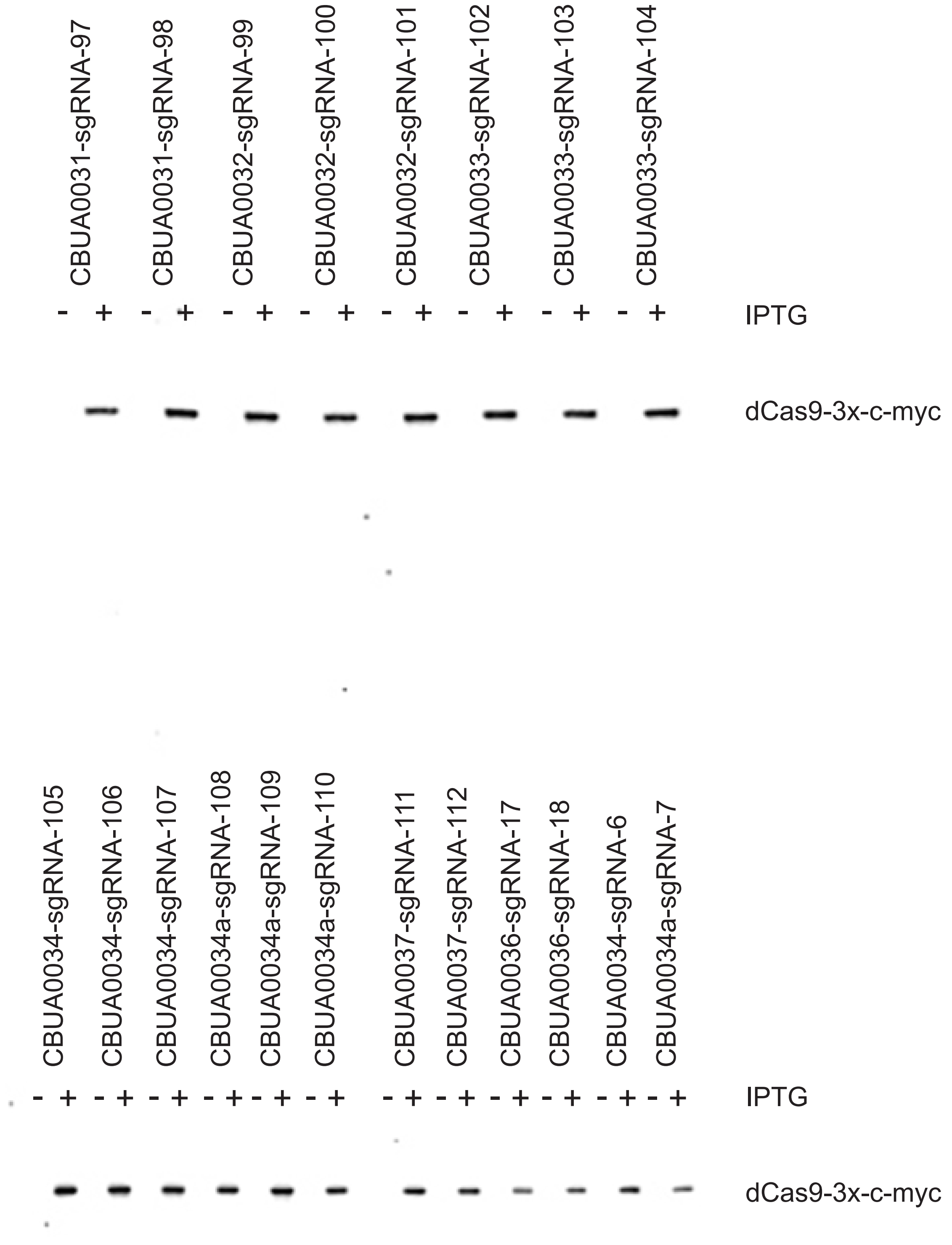

Supplemental figure 5

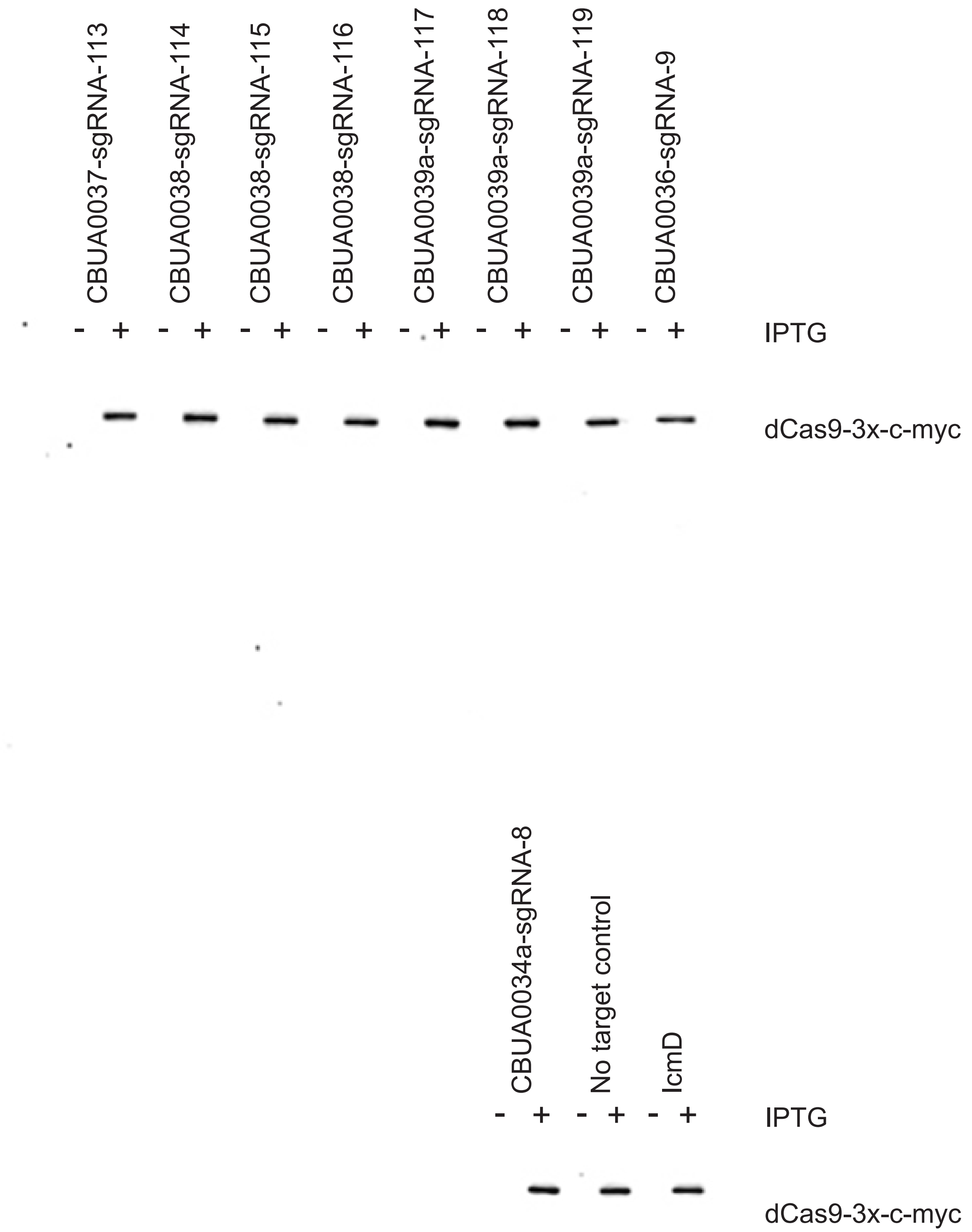

Supplemental figure 6

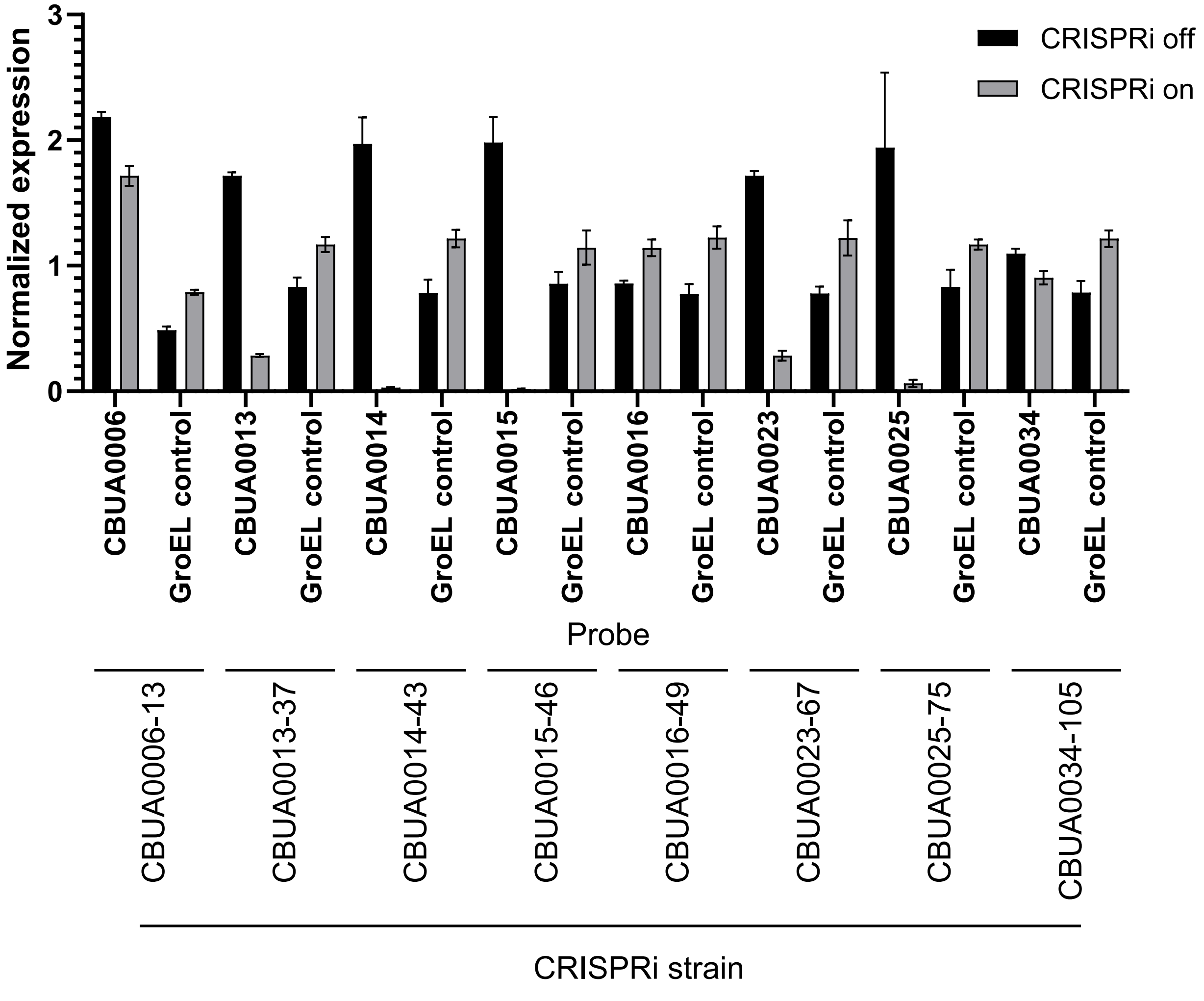

Supplemental figure 7

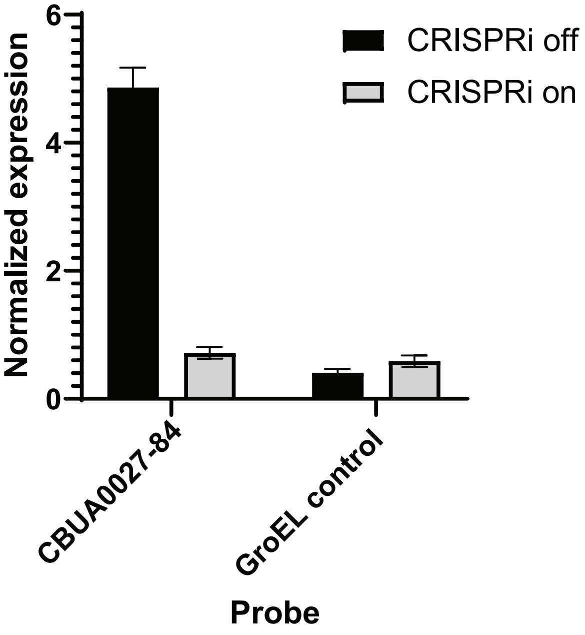

### Supplemental figure 8

A

```
                                                              -35
cbuA0028    CTGATGGAAATCTGCTCGAGATAGGCAATGAGTACAGATTGACAAATAGTTCACTTAGTG 60
cbuA0028G   ------------------------------------------------------------ 0

              -10
cbuA0028    ATATAGTCTTAACAATAAGGGAAGTTATGAGGATATTTAAAACACGCTATTTCCATCGAT 120
cbuA0028G   --AGTTTTTTGGAGAATAAATAACTAGGGCGCGTCGTCAATTAACCCTGCCACCGTCGTC 58
              *   * **    *  *   ** *   * *  *  * **   ** **    ** ***

cbuA0028    GGGCAAAATCAGAAAATCTGTCCGATAGCCAACTGAAAACGGCCAT--TAGTGAACTGAT 178
cbuA0028G   CCG----------CGGCTTGTCCGCGGATCCAGAGAAACCACGTTAAGTTCTAGAAATT 108
                  *               ******   *        ****  *  *       *     *

cbuA0028    GAAGGGATTACACGATGGAGATTTAGGTTCCTATATCTATAAAAAACGGATTCCTATACT 238
cbuA0028G   GGCTAGTTAATTAAGTATTCTGGGATATTCCTTTATCTATAAAAAACGGATTCCTATACT 168
            *    * * *     *        *  ***** ***************************

cbuA0028    AGGTAAAGGAAAACGAGGTGGGCTTCGAACAATAATTGCTTACCGAGCAGAAGATAAAGC 298
cbuA0028G   AGGTAAAGGAAAACGAGGTGGGCTTCGAACAATAATTGCTTACCGAGCAGAAGATAAAGC 228
            ************************************************************

cbuA0028    TTTTTTTGTATATGGTTATGCGAAAAATGTTCAAGCCAATATAACTCCAAAAGAAAAGGA 358
cbuA0028G   TTTTTTTGTATATGGTTATGCGAAAAATGTTCAAGCCAATATAACTCCAAAAGAAAAGGA 288
            ************************************************************

cbuA0028    GGCGTACAAAAAGTTGTCAAAAACTTATTTTGATATGAAAGAATTAGAGCTTCAATCATT 418
cbuA0028G   GGCGTACAAAAAGTTGTCAAAAACTTATTTTGATATGAAAGAATTAGAGCTTCAATCATT 348
            ************************************************************

cbuA0028    ATTAAAAATAGGCGAACTGATAGAGGTGCTGTGA   452
cbuA0028G   ATTAAAAATAGGCGAACTGATAGAGGTGCTGTGA   382
            **********************************
```

B

```
cbuA0027    ATGAAAGATAAAAAAAAGAAAAGTAAAACTCGTTTTAAACCTTTATCGCCCATTGAGGCC 60
cbuA0027G   ATGAAAGATAAAAAAAAGAAAAGTAAAACTCGTTTTAAACCTTTATCGCCCATTGAGGCC 60
            ************************************************************

cbuA0027    GTTCATGAATCTGCAAAAGGATTATATGATGCCGGTGTTATTGACGCGACAACAATGCAT 120
cbuA0027G   GTTCATGAATCTGCAAAAGGATTATATGATGCCGGTGTTATTGACGCGACAACAATGCAT 120
            ************************************************************

cbuA0027    GAATTTGATGCATTGTGTTTGACTCCAGTCCGTGAATTATCGCCTCGTGAAATTAAGCGC 180
cbuA0027G   GAATTTGATGCATTGTGTTTGACTCCAGTCCGTGAATTATCGCCTCGTGAAATTAAGCGC 180
            ************************************************************

cbuA0027    ATTCGAATTCATGAAAAAGTGAGCCAGGCCGTTTTTGCAAAATATTTGAATACCAGTGTT 240
cbuA0027G   ATTCGAATTCATGAAAAAGTGAGCCAGGCCGTTTTTGCAAAATATTTGAATACCAGTGTT 240
            ************************************************************

cbuA0027    TCTACGGTAAAGCAATGGGAGTTGGGTGAGAAACACCCTCGAGGCACTTCATTAAAATTA 300
cbuA0027G   TCTACGGTAAAGCAATGGGAGTTGGGTGAGAAACATCCTCGAGGCACTTCATTAAAATTA 300
            *********************************** ************************

cbuA0027    CTGAACCTCGTTGATAGAAAAGGCCTTCAAGCCATCGCTTAG 342
cbuA0027G   CTGAACCTCGTTGATAGAAAAGGCCTTCAAGCCATCGCTTAG 342
            ******************************************
```

Supplemental figure 9

A

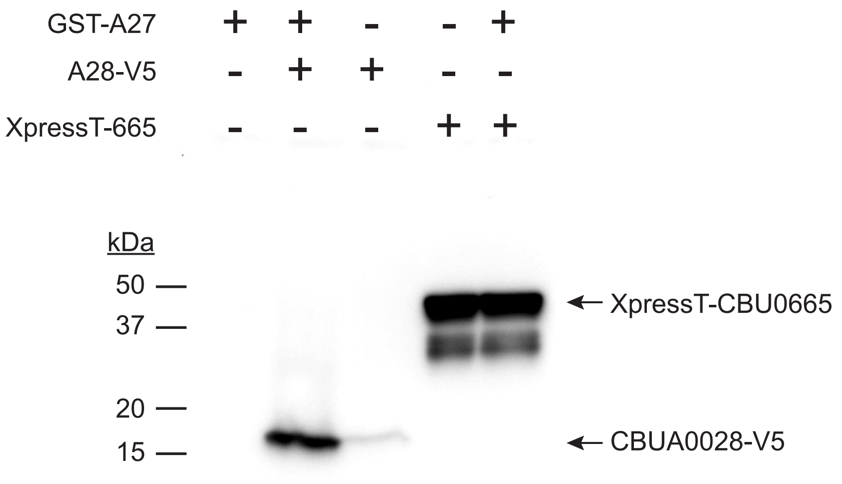


B

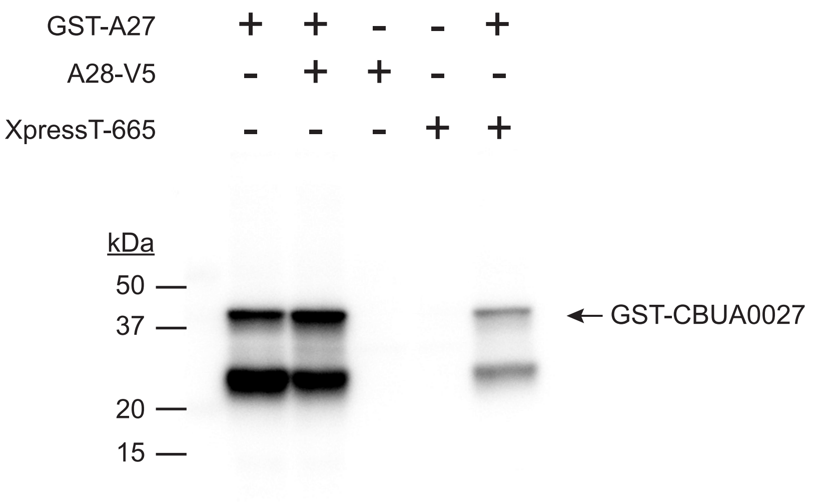

### Supplemental figure 10

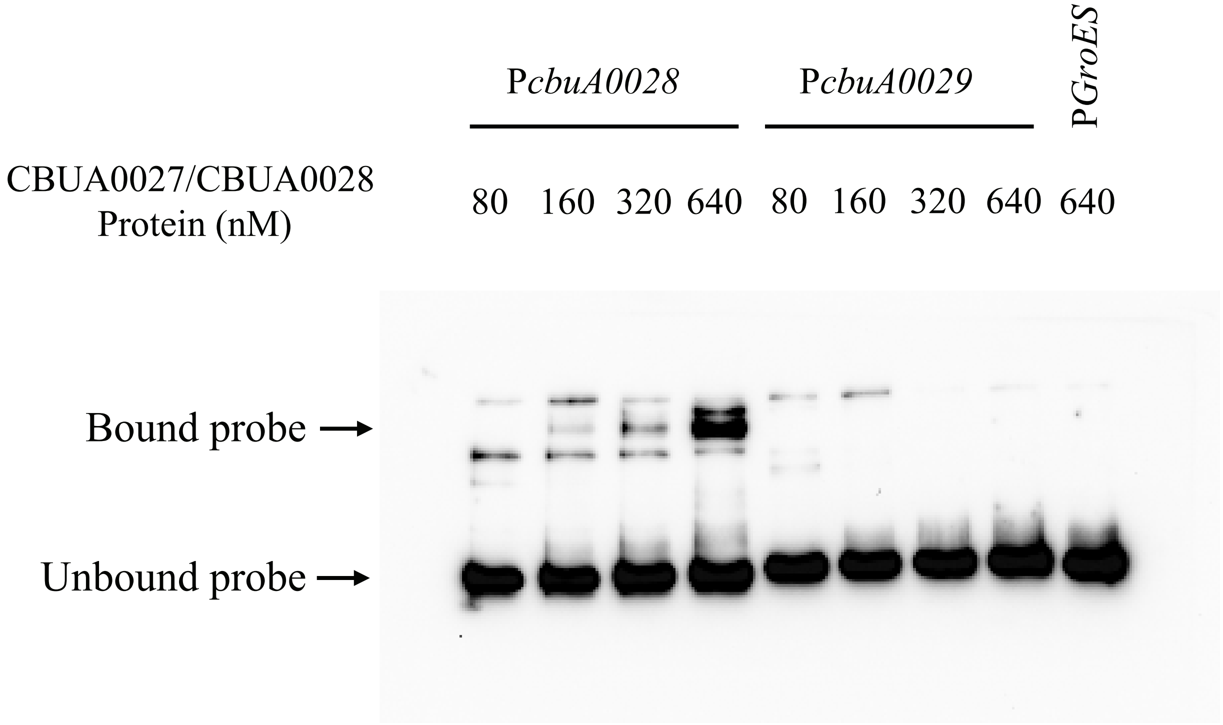

### Supplemental figure 11

```
                                                            -35
cbuA0027/cbuA0028     CTGATGGAAATCTGCTCGAGATAGGCAATGAGTACAGATTGACAAATAGTTCACTTAGTG   60
cbuA0028G/cbuA0027G   ------------------------------------------------------------   0

                        -10
cbuA0027/cbuA0028     ATATAGTCTTAACAATAAGGGAAGTTATGAGGATATTT-AAA------AC----------   103
cbuA0028G/cbuA0027G   ----------------------AGTTTTTTGGAGAATAAATAACTAGGGCGCGTCGTCAA   38
                                            ****:* :*** *:*: *:*      .*

cbuA0027/cbuA0028     ---ACGCTATTTCCATCGATGGGCAAAATCAGAAAATCTGTCCGATA--GCCAACTGAAA   158
cbuA0028G/cbuA0027G   TTAACCCTGCCACCGTCGTCCCGCG------G----CTTGTCCGCGGATCCAGA-G-AA   86
                         ** **.  :**.***:   **.      *      ******. .   ***.. * **
                                                          -35
cbuA0027/cbuA0028     ACGGCCATTAGTGAACTGATGAAGGGATTACAC-GATGGAGA---TTTAGGTTCCTATAT   214
cbuA0028G/cbuA0027G   ACCACGTT--AAGTTCTAGAAATTGGCTAGTTAATTAAGTATTCTGGGATATTCCTTTAT   144
                      ** .* :*  .:*::**..:.*: **.*:. :.  ::.*:.:      * .*****:***
                        -10
cbuA0027/cbuA0028     CTATAAAAAACGGATTCCTATACTAGGTAAAGGAAAACGAGGTGGGCTTCGAACAATAAT   274
cbuA0028G/cbuA0027G   CTATAAAAAACGGATTCCTATACTAGGTAAAGGAAAACGAGGTGGGCTTCGAACAATAAT   204
                      ************************************************************

cbuA0027/cbuA0028     TGCTTACCGAGCAGAAGATAAAGCTTTTTTTGTATATGGTTATGCGAAAAATGTTCAAGC   334
cbuA0028G/cbuA0027G   TGCTTACCGAGCAGAAGATAAAGCTTTTTTTGTATATGGTTATGCGAAAAATGTTCAAGC   264
                      ************************************************************
                                                                             -35
cbuA0027/cbuA0028     CAATATAACTCCAAAAGAAAAGGAGGCGTACAAAAAGTTGTCAAAAACTTATTTTGATAT   394
cbuA0028G/cbuA0027G   CAATATAACTCCAAAAGAAAAGGAGGCGTACAAAAAGTTGTCAAAAACTTATTTTGATAT   324
                      ************************************************************
                                        -10
cbuA0027/cbuA0028     GAAAGAATTAGAGCTTCAATCATTATTAAAAATAGGCGAACTGATAGAGGTGCTGTGATG   454
cbuA0028G/cbuA0027G   GAAAGAATTAGAGCTTCAATCATTATTAAAAATAGGCGAACTGATAGAGGTGCTGTGATG   384
                      ************************************************************

cbuA0027/cbuA0028     AAAGATAAAAAAAAGAAAAGTAAAACTCGTTTTAAACCTTTATCGCCCATTGAGGCCGTT   514
cbuA0028G/cbuA0027G   AAAGATAAAAAAAAGAAAAGTAAAACTCGTTTTAAACCTTTATCGCCCATTGAGGCCGTT   444
                      ************************************************************

cbuA0027/cbuA0028     CATGAATCTGCAAAAGGATTATATGATGCCGGTGTTATTGACGCGACAACAATGCATGAA   574
cbuA0028G/cbuA0027G   CATGAATCTGCAAAAGGATTATATGATGCCGGTGTTATTGACGCGACAACAATGCATGAA   504
                      ************************************************************

cbuA0027/cbuA0028     TTTGATGCATTGTGTTTGACTCCAGTCCGTGAATTATCGCCTCGTGAAATTAAGCGCATT   634
cbuA0028G/cbuA0027G   TTTGATGCATTGTGTTTGACTCCAGTCCGTGAATTATCGCCTCGTGAAATTAAGCGCATT   564
                      ************************************************************

cbuA0027/cbuA0028     CGAATTCATGAAAAAGTGAGCCAGGCCGTTTTTGCAAAATATTTGAATACCAGTGTTTCT   694
cbuA0028G/cbuA0027G   CGAATTCATGAAAAAGTGAGCCAGGCCGTTTTTGCAAAATATTTGAATACCAGTGTTTCT   624
                      ************************************************************

cbuA0027/cbuA0028     ACGGTAAAGCAATGGGAGTTGGGTGAGAAACACCCTCGAGGCACTTCATTAAAATTACTG   754
cbuA0028G/cbuA0027G   ACGGTAAAGCAATGGGAGTTGGGTGAGAAACATCCTCGAGGCACTTCATTAAAATTACTG   684
                      ******************************** ***************************

cbuA0027/cbuA0028     AACCTCGTTGATAGAAAAGGCCTTCAAGCCATCGCTTAG                        793
cbuA0028G/cbuA0027G   AACCTCGTTGATAGAAAAGGCCTTCAAGCCATCGCTTAG                        723
                      ***************************************
```

Supplemental figure 12

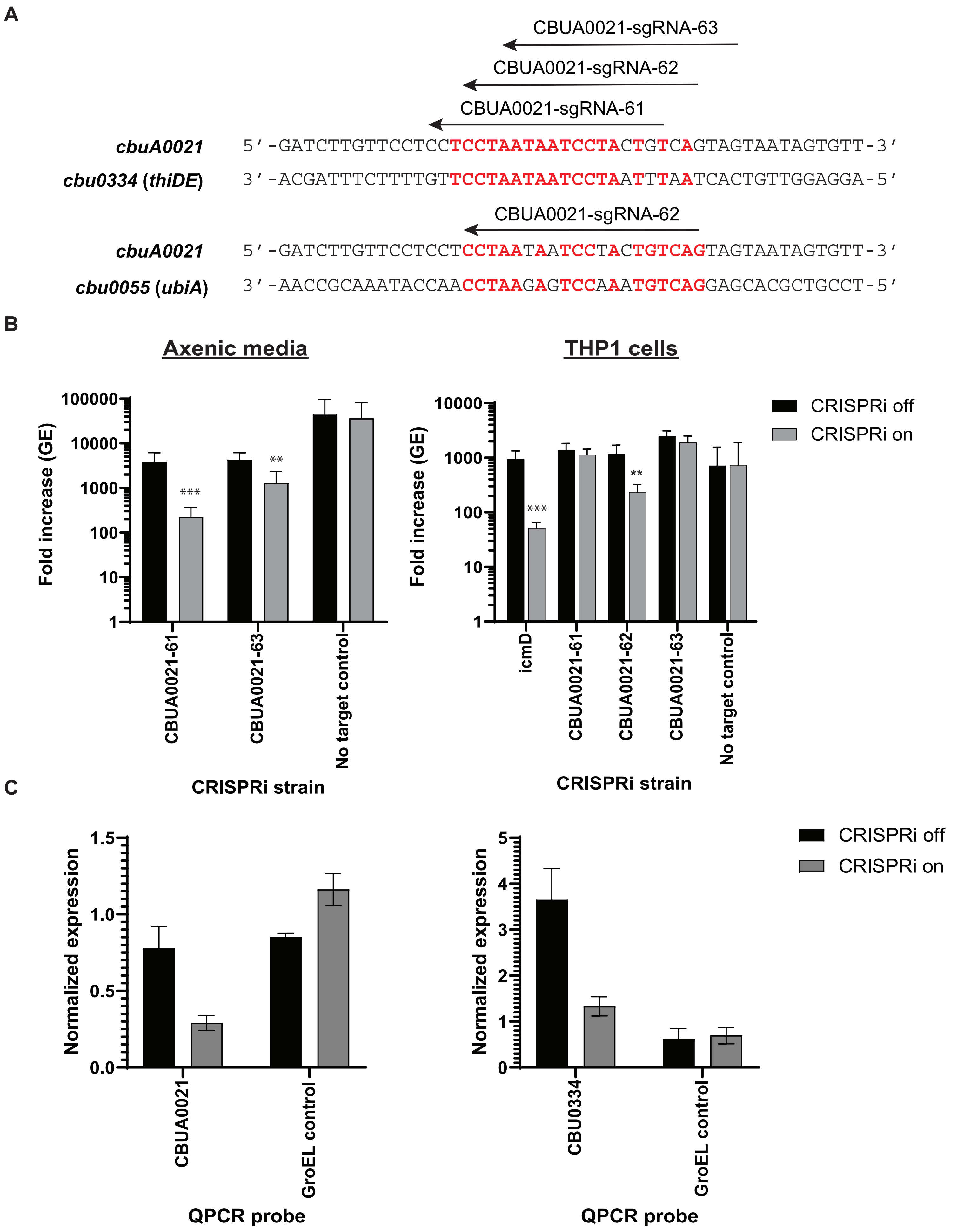

Supplemental figure 13

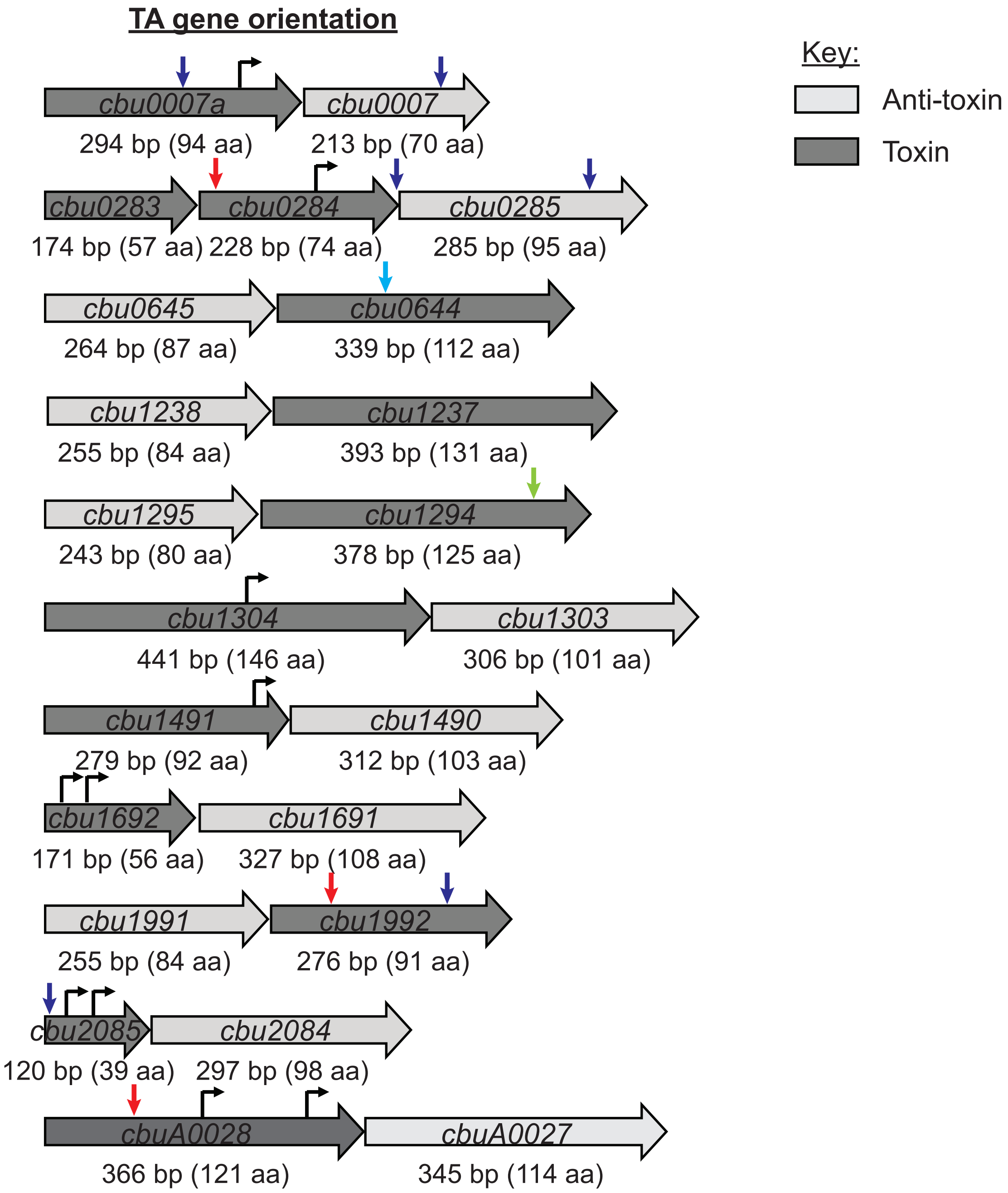
