## Supplemental Tables S1 and S2 for "A toxin-antitoxin system ensures plasmid stability in *Coxiella burnetii*"

| Strain or plasmid | Genotype and/or phenotype | Source/reference |
|---|---|---|
| Strain | | |
| *E. coli* Stellar cells | *F–, endA1, supE44, thi-1, recA1, relA1, gyrA96, phoA, Φ80D lacZΔ M15, Δ (lacZYA - argF) U169, Δ(mrr - hsdRMS - mcrBC), ΔmcrA, λ–* | Takara |
| *E. coli* PIR1 cells | *F- Δlac169, rpoS(Am), robA1, creC510, hsdR514, endA, recA1, uidA(ΔMluI)::pir-116* | Invitrogen |
| *C. burnetii* Nine Mile RSA439 (NMII) | Phase II, clone 4 | Beare *et_al* 2012 |
| NMII ΔQpH1 | NMII QpH1-less strain containing pB-TyrB-QpH1ori | This study |
| Plasmids | | |
| QpH1 | Endogenous *C. burnetii* plasmid | This study |
| pTnS2::*1169$^{P}$ -tnsABCD* | *cbu1169* promoter cloned into pTnS2, R6K *ori*, Amp$^{r}$ | Beare *et_al* 2011 |
| pJB-CAT-LacIO-MC | *lacIO* cloned into pJB-CAT, replacing *lacIq* -P*tac*, Amp$^{r}$, Cm$^{r}$ | This study |
| *Coxiella shuttle vector* | | |
| pJB-CAT | pJB2581-*mCherry* containing cat driven by *1169$^{P}$*; Amp$^{r}$, Cm$^{r}$ | Omsland *et_al* 2011 |
| pJB-CAT-*tyrB* | pJB-CAT containing *tyrB* driven by *1169$^{P}$*, Cmr, Amp$^{r}$ | This study |
| pJC-CAT | pJC84 containing *cat* driven by *1169$^{P}$*; Cm$^{r}$ | Beare *et_al* 2011 |
| pB-TyrB | *cbu1169* promoter driving expression of *cat* and *sacB*, pMB1 *ori*, Cm$^{r}$ | This study |
| pB-TyrB-QpH1ori | *cbuA0036-cbuA0037-cbuA0038-cbuA0039* cloned into pB-TyrB, Cm$^{r}$ | This study |
| *CRISPRi constructs* | | |
| pJMP1356 | pMiniTn7 vector containing elements of a CRISPRi system, dCas9, lacI, and targeting sgRNA, R6k *ori*, Amp$^{r}$, Cm$^{r}$ | Peters *et_al* 2019 |
| pB-iCRISPR | *proBA* cloned in place of cat in pJMP1356, R6K *ori*, Amp$^{r}$ | This study |
| pB-iCRISPR-*icmD* -sgRNA-1 | pB-iCRISPR containing *cbu1624* -sgRNA-1 target sequence, R6K *ori*, Amp$^{r}$ | This study |
| pB-iCRISPR-*icmD* -sgRNA-2 | pB-iCRISPR containing *cbu1624* -sgRNA-2 target sequence, R6K *ori*, Amp$^{r}$ | This study |
| pB-iCRISPR-*icmD* -sgRNA-1-*scvA* -sgRNA-1 | pB-iCRISPR containing *cbu1624* -sgRNA-1 (*icmD*) and *cbu1267a* (*scvA*) target sequences, R6K *ori*, Amp$^{r}$ | This study |
| pB-iCRISPR-*icmD* -sgRNA-3 | pB-iCRISPR containing *cbu1624* -sgRNA-3 target sequence, R6K *ori*, Amp$^{r}$ | This study |
| pB-iCRISPR-*cbuA0001* -sgRNA-1 | pB-iCRISPR containing *cbuA0001* -sgRNA-1 target sequence, R6K *ori*, Amp$^{r}$ | This study |
| pB-iCRISPR-*cbuA0001* -sgRNA-2 | pB-iCRISPR containing *cbuA0001* -sgRNA-2 target sequence, R6K *ori*, Amp$^{r}$ | This study |
| pB-iCRISPR-*cbuA0001* -sgRNA-3 | pB-iCRISPR containing *cbuA0001* -sgRNA-3 target sequence, R6K *ori*, Amp$^{r}$ | This study |
| pB-iCRISPR-*cbuA0003* -sgRNA-4 | pB-iCRISPR containing *cbuA0003* -sgRNA-4 target sequence, R6K *ori*, Amp$^{r}$ | This study |
| pB-iCRISPR-*cbuA0003* -sgRNA-5 | pB-iCRISPR containing *cbuA0003* -sgRNA-5 target sequence, R6K *ori*, Amp$^{r}$ | This study |
| pB-iCRISPR-*cbuA0003* -sgRNA-6 | pB-iCRISPR containing *cbuA0003* -sgRNA-6 target sequence, R6K *ori*, Amp$^{r}$ | This study |
| pB-iCRISPR-*cbuA0003a* -sgRNA-7 | pB-iCRISPR containing *cbuA0003a* -sgRNA-7 target sequence, R6K *ori*, Amp$^{r}$ | This study |
| pB-iCRISPR-*cbuA0003a* -sgRNA-8 | pB-iCRISPR containing *cbuA0003a* -sgRNA-8 target sequence, R6K *ori*, Amp$^{r}$ | This study |
| pB-iCRISPR-*cbuA0003a* -sgRNA-9 | pB-iCRISPR containing *cbuA0003a* -sgRNA-9 target sequence, R6K *ori*, Amp$^{r}$ | This study |
| pB-iCRISPR-*cbuA0005* -sgRNA-10 | pB-iCRISPR containing *cbuA0005* -sgRNA-10 target sequence, R6K *ori*, Amp$^{r}$ | This study |
| pB-iCRISPR-*cbuA0005* -sgRNA-11 | pB-iCRISPR containing *cbuA0005* -sgRNA-11 target sequence, R6K *ori*, Amp$^{r}$ | This study |
| pB-iCRISPR-*cbuA0005* -sgRNA-12 | pB-iCRISPR containing *cbuA0005* -sgRNA-12 target sequence, R6K *ori*, Amp$^{r}$ | This study |
| pB-iCRISPR-*cbuA0006* -sgRNA-13 | pB-iCRISPR containing *cbuA0006* -sgRNA-13 target sequence, R6K *ori*, Amp$^{r}$ | This study |
| pB-iCRISPR-*cbuA0006* -sgRNA-14 | pB-iCRISPR containing *cbuA0006* -sgRNA-14 target sequence, R6K *ori*, Amp$^{r}$ | This study |
| pB-iCRISPR-*cbuA0006* -sgRNA-15 | pB-iCRISPR containing *cbuA0006* -sgRNA-15 target sequence, R6K *ori*, Amp$^{r}$ | This study |
| pB-iCRISPR-*cbuA0007* -sgRNA-16 | pB-iCRISPR containing *cbuA0007* -sgRNA-16 target sequence, R6K *ori*, Amp$^{r}$ | This study |
| pB-iCRISPR-*cbuA0007* -sgRNA-17 | pB-iCRISPR containing *cbuA0007* -sgRNA-17 target sequence, R6K *ori*, Amp$^{r}$ | This study |
| pB-iCRISPR-*cbuA0007* -sgRNA-18 | pB-iCRISPR containing *cbuA0007* -sgRNA-18 target sequence, R6K *ori*, Amp$^{r}$ | This study |
| pB-iCRISPR-*cbuA0008* -sgRNA-19 | pB-iCRISPR containing *cbuA0008* -sgRNA-19 target sequence, R6K *ori*, Amp$^{r}$ | This study |
| pB-iCRISPR-*cbuA0008* -sgRNA-20 | pB-iCRISPR containing *cbuA0008* -sgRNA-20 target sequence, R6K *ori*, Amp$^{r}$ | This study |
| pB-iCRISPR-*cbuA0008* -sgRNA-21 | pB-iCRISPR containing *cbuA0008* -sgRNA-21 target sequence, R6K *ori*, Amp$^{r}$ | This study |
| pB-iCRISPR-*cbuA0008a* -sgRNA-22 | pB-iCRISPR containing *cbuA0008a* -sgRNA-22 target sequence, R6K *ori*, Amp$^{r}$ | This study |
| pB-iCRISPR-*cbuA000* 8a-sgRNA-23 | pB-iCRISPR containing *cbuA0008a* -sgRNA-23 target sequence, R6K *ori*, Amp$^{r}$ | This study |
| pB-iCRISPR-*cbuA0008a* -sgRNA-24 | pB-iCRISPR containing *cbuA0008a* -sgRNA-24 target sequence, R6K *ori*, Amp$^{r}$ | This study |

| | | |
|---|---|---|
| pB-iCRISPR-*cbuA0008b* -sgRNA-25 | pB-iCRISPR containing *cbuA0008b* -sgRNA-25 target sequence, R6K *ori* , Amp$^r$ | This study |
| pB-iCRISPR-*cbuA0008b* -sgRNA-26 | pB-iCRISPR containing *cbuA0008b* -sgRNA-26 target sequence, R6K *ori* , Amp$^r$ | This study |
| pB-iCRISPR-*cbuA0008b* -sgRNA-27 | pB-iCRISPR containing *cbuA0008b* -sgRNA-27 target sequence, R6K *ori* , Amp$^r$ | This study |
| pB-iCRISPR-*cbuA0010* -sgRNA-28 | pB-iCRISPR containing *cbuA0010* -sgRNA-28 target sequence, R6K *ori* , Amp$^r$ | This study |
| pB-iCRISPR-*cbuA0010* -sgRNA-29 | pB-iCRISPR containing *cbuA0010* -sgRNA-29 target sequence, R6K *ori* , Amp$^r$ | This study |
| pB-iCRISPR-*cbuA0010* -sgRNA-30 | pB-iCRISPR containing *cbuA0010* -sgRNA-30 target sequence, R6K *ori* , Amp$^r$ | This study |
| pB-iCRISPR-*cbuA0011* -sgRNA-31 | pB-iCRISPR containing *cbuA0011* -sgRNA-31 target sequence, R6K *ori* , Amp$^r$ | This study |
| pB-iCRISPR-*cbuA0011* -sgRNA-32 | pB-iCRISPR containing *cbuA0011* -sgRNA-32 target sequence, R6K *ori* , Amp$^r$ | This study |
| pB-iCRISPR-*cbuA0011* -sgRNA-33 | pB-iCRISPR containing *cbuA0011-* sgRNA-33 target sequence, R6K *ori* , Amp$^r$ | This study |
| pB-iCRISPR-*cbuA0012* -sgRNA-34 | pB-iCRISPR containing *cbuA0012* -sgRNA-34 target sequence, R6K *ori* , Amp$^r$ | This study |
| pB-iCRISPR-*cbuA0012* -sgRNA-35 | pB-iCRISPR containing *cbuA0012* -sgRNA-35 target sequence, R6K *ori* , Amp$^r$ | This study |
| pB-iCRISPR-*cbuA0012* -sgRNA-36 | pB-iCRISPR containing *cbuA0012* -sgRNA-36 target sequence, R6K *ori* , Amp$^r$ | This study |
| pB-iCRISPR-*cbuA0013* -sgRNA-37 | pB-iCRISPR containing *cbuA0013* -sgRNA-37 target sequence, R6K *ori* , Amp$^r$ | This study |
| pB-iCRISPR-*cbuA0013* -sgRNA-38 | pB-iCRISPR containing *cbuA0013* -sgRNA-38 target sequence, R6K *ori* , Amp$^r$ | This study |
| pB-iCRISPR-*cbuA0013* -sgRNA-39 | pB-iCRISPR containing *cbuA0013* -sgRNA-39 target sequence, R6K *ori* , Amp$^r$ | This study |
| pB-iCRISPR-*cbuA0013* a-sgRNA-40 | pB-iCRISPR containing *cbuA0013a* -sgRNA-40 target sequence, R6K *ori* , Amp$^r$ | This study |
| pB-iCRISPR-*cbuA0013a* -sgRNA-41 | pB-iCRISPR containing *cbuA0013a* -sgRNA-41 target sequence, R6K *ori* , Amp$^r$ | This study |
| pB-iCRISPR-*cbuA0013a* -sgRNA-42 | pB-iCRISPR containing *cbuA0013a* -sgRNA-42 target sequence, R6K *ori* , Amp$^r$ | This study |
| pB-iCRISPR-*cbuA0014* -sgRNA-43 | pB-iCRISPR containing *cbuA0014* -sgRNA-43 target sequence, R6K *ori* , Amp$^r$ | This study |
| pB-iCRISPR-*cbuA0014* -sgRNA-44 | pB-iCRISPR containing *cbuA0014* -sgRNA-44 target sequence, R6K *ori* , Amp$^r$ | This study |
| pB-iCRISPR-*cbuA0014* -sgRNA-45 | pB-iCRISPR containing *cbuA0014* -sgRNA-45 target sequence, R6K *ori* , Amp$^r$ | This study |
| pB-iCRISPR-*cbuA0015* -sgRNA-46 | pB-iCRISPR containing *cbuA0015* -sgRNA-46 target sequence, R6K *ori* , Amp$^r$ | This study |
| pB-iCRISPR-*cbuA0015* -sgRNA-47 | pB-iCRISPR containing *cbuA0015* -sgRNA-47 target sequence, R6K *ori* , Amp$^r$ | This study |
| pB-iCRISPR-*cbuA0015* -sgRNA-48 | pB-iCRISPR containing *cbuA0015* -sgRNA-48 target sequence, R6K *ori* , Amp$^r$ | This study |
| pB-iCRISPR-*cbuA0016* -sgRNA-49 | pB-iCRISPR containing *cbuA0016* -sgRNA-49 target sequence, R6K *ori* , Amp$^r$ | This study |
| pB-iCRISPR-*cbuA0016* -sgRNA-50 | pB-iCRISPR containing *cbuA0016* -sgRNA-50 target sequence, R6K *ori* , Amp$^r$ | This study |
| pB-iCRISPR-*cbuA0016* -sgRNA-51 | pB-iCRISPR containing *cbuA0016* -sgRNA-51 target sequence, R6K *ori* , Amp$^r$ | This study |
| pB-iCRISPR-*cbuA0017* -sgRNA-52 | pB-iCRISPR containing *cbuA0017* -sgRNA-52 target sequence, R6K *ori* , Amp$^r$ | This study |
| pB-iCRISPR-*cbuA0017* -sgRNA-53 | pB-iCRISPR containing *cbuA0017* -sgRNA-53 target sequence, R6K *ori* , Amp$^r$ | This study |
| pB-iCRISPR-*cbuA0017* -sgRNA-54 | pB-iCRISPR containing *cbuA0017* -sgRNA-54 target sequence, R6K *ori* , Amp$^r$ | This study |
| pB-iCRISPR-*cbuA0018* -sgRNA-55 | pB-iCRISPR containing *cbuA0018* -sgRNA-55 target sequence, R6K *ori* , Amp$^r$ | This study |
| pB-iCRISPR-*cbuA0018* -sgRNA-56 | pB-iCRISPR containing *cbuA0018* -sgRNA-56 target sequence, R6K *ori* , Amp$^r$ | This study |
| pB-iCRISPR-*cbuA0018* -sgRNA-57 | pB-iCRISPR containing *cbuA0018* -sgRNA-57 target sequence, R6K *ori* , Amp$^r$ | This study |
| pB-iCRISPR-*cbuA0020* -sgRNA-58 | pB-iCRISPR containing *cbuA0020* -sgRNA-58 target sequence, R6K *ori* , Amp$^r$ | This study |
| pB-iCRISPR-*cbuA0020* -sgRNA-59 | pB-iCRISPR containing *cbuA0020* -sgRNA-59 target sequence, R6K *ori,* Amp$^r$ | This study |
| pB-iCRISPR-*cbuA0020* -sgRNA-60 | pB-iCRISPR containing *cbuA0020* -sgRNA-60 target sequence, R6K *ori* , Amp$^r$ | This study |
| pB-iCRISPR-*cbuA0021* -sgRNA-61 | pB-iCRISPR containing *cbuA0021* -sgRNA-61 target sequence, R6K *ori* , Amp$^r$ | This study |
| pB-iCRISPR-*cbuA0021* -sgRNA-62 | pB-iCRISPR containing *cbuA0021* -sgRNA-62 target sequence, R6K *ori* , Amp$^r$ | This study |
| pB-iCRISPR-*cbuA0021* -sgRNA-63 | pB-iCRISPR containing *cbuA0021* -sgRNA-63 target sequence, R6K *ori* , Amp$^r$ | This study |
| pB-iCRISPR-*cbuA0022* -sgRNA-64 | pB-iCRISPR containing *cbuA0022* -sgRNA-64 target sequence, R6K *ori* , Amp$^r$ | This study |
| pB-iCRISPR-*cbuA0022* -sgRNA-65 | pB-iCRISPR containing *cbuA0022* -sgRNA-65 target sequence, R6K *ori* , Amp$^r$ | This study |
| pB-iCRISPR-*cbuA0022* -sgRNA-66 | pB-iCRISPR containing *cbuA0022* -sgRNA-66 target sequence, R6K *ori,* Amp$^r$ | This study |
| pB-iCRISPR-*cbuA0023* -sgRNA-67 | pB-iCRISPR containing *cbuA0023* -sgRNA-67 target sequence, R6K *ori* , Amp$^r$ | This study |
| pB-iCRISPR-*cbuA0023* -sgRNA-68 | pB-iCRISPR containing *cbuA0023* -sgRNA-68 target sequence, R6K *ori* , Amp$^r$ | This study |
| pB-iCRISPR-*cbuA0023* -sgRNA-69 | pB-iCRISPR containing *cbuA0023* -sgRNA-69 target sequence, R6K *ori* , Amp$^r$ | This study |
| pB-iCRISPR-*cbuA0023a* -sgRNA-70 | pB-iCRISPR containing *cbuA0023a* -sgRNA-70 target sequence, R6K *ori* , Amp$^r$ | This study |
| pB-iCRISPR-*cbuA0023a* -sgRNA-71 | pB-iCRISPR containing *cbuA0023a* -sgRNA-71 target sequence, R6K *ori* , Amp$^r$ | This study |
| pB-iCRISPR-*cbuA0024* -sgRNA-72 | pB-iCRISPR containing *cbuA0024* -sgRNA-72 target sequence, R6K *ori* , Amp$^r$ | This study |
| pB-iCRISPR-*cbuA0024* -sgRNA-73 | pB-iCRISPR containing *cbuA0024* -sgRNA-73 target sequence, R6K *ori* , Amp$^r$ | This study |
| pB-iCRISPR-*cbuA0024* -sgRNA-74 | pB-iCRISPR containing *cbuA0024* -sgRNA-74 target sequence, R6K *ori* , Amp$^r$ | This study |
| pB-iCRISPR-*cbuA0025* -sgRNA-75 | pB-iCRISPR containing *cbuA0025* -sgRNA-75 target sequence, R6K *ori* , Amp$^r$ | This study |

| | | |
|---|---|---|
| pB-iCRISPR-*cbuA0025* -sgRNA-76 | pB-iCRISPR containing *cbuA0025*- sgRNA-76 target sequence, R6K *ori* , $Amp^r$ | This study |
| pB-iCRISPR-*cbuA0025* -sgRNA-77 | pB-iCRISPR containing *cbuA0025* -sgRNA-77 target sequence, R6K *ori* , $Amp^r$ | This study |
| pB-iCRISPR-*cbuA0026* -sgRNA-78 | pB-iCRISPR containing *cbuA0026* -sgRNA-78 target sequence, R6K *ori* , $Amp^r$ | This study |
| pB-iCRISPR-*cbuA0026* -sgRNA-79 | pB-iCRISPR containing *cbuA0026* -sgRNA-79 target sequence, R6K *ori* , $Amp^r$ | This study |
| pB-iCRISPR-*cbuA0026* -sgRNA-80 | pB-iCRISPR containing *cbuA0026* -sgRNA-80 target sequence, R6K *ori* , $Amp^r$ | This study |
| pB-iCRISPR-*cbuA0026a* -sgRNA-81 | pB-iCRISPR containing *cbuA0026a* -sgRNA-81 target sequence, R6K *ori* , $Amp^r$ | This study |
| pB-iCRISPR-*cbuA0026a* -sgRNA-82 | pB-iCRISPR containing *cbuA0026a* -sgRNA-82 target sequence, R6K *ori* , $Amp^r$ | This study |
| pB-iCRISPR-*cbuA0026a* -sgRNA-83 | pB-iCRISPR containing *cbuA0026a* -sgRNA-83 target sequence, R6K *ori* , $Amp^r$ | This study |
| pB-iCRISPR-*cbuA0027* -sgRNA-84 | pB-iCRISPR containing *cbuA0027* -sgRNA-84 target sequence, R6K *ori* , $Amp^r$ | This study |
| pB-iCRISPR-*cbuA0027* -sgRNA-85 | pB-iCRISPR containing *cbuA0027* -sgRNA-85 target sequence, R6K *ori* , $Amp^r$ | This study |
| pB-iCRISPR-*cbuA0027* -sgRNA-86 | pB-iCRISPR containing *cbuA0027* -sgRNA-86 target sequence, R6K *ori* , $Amp^r$ | This study |
| pB-iCRISPR-*cbuA0028* -sgRNA-87 | pB-iCRISPR containing *cbuA0028* -sgRNA-87 target sequence, R6K *ori* , $Amp^r$ | This study |
| pB-iCRISPR-*cbuA0028* -sgRNA-88 | pB-iCRISPR containing *cbuA0028* -sgRNA-88 target sequence, R6K *ori* , $Amp^r$ | This study |
| pB-iCRISPR-*cbuA0028* -sgRNA-89 | pB-iCRISPR containing *cbuA0028* -sgRNA-89 target sequence, R6K *ori* , $Amp^r$ | This study |
| pB-iCRISPR-*cbuA0029* -sgRNA-90 | pB-iCRISPR containing *cbuA0029* -sgRNA-90 target sequence, R6K *ori* , $Amp^r$ | This study |
| pB-iCRISPR-*cbuA0029* -sgRNA-91 | pB-iCRISPR containing *cbuA0029* -sgRNA-91 target sequence, R6K *ori* , $Amp^r$ | This study |
| pB-iCRISPR-*cbuA0029* -sgRNA-92 | pB-iCRISPR containing *cbuA0029* -sgRNA-92 target sequence, R6K *ori* , $Amp^r$ | This study |
| pB-iCRISPR-*cbuA0029a* -sgRNA-93 | pB-iCRISPR containing *cbuA0029a* -sgRNA-93 target sequence, R6K *ori* , $Amp^r$ | This study |
| pB-iCRISPR-*cbuA0029a* -sgRNA-94 | pB-iCRISPR containing *cbuA0029a* -sgRNA-94 target sequence, R6K *ori* , $Amp^r$ | This study |
| pB-iCRISPR-*cbuA0029a* -sgRNA-95 | pB-iCRISPR containing *cbuA0029a* -sgRNA-95 target sequence, R6K *ori* , $Amp^r$ | This study |
| pB-iCRISPR-*cbuA0031* -sgRNA-96 | pB-iCRISPR containing *cbuA0031* -sgRNA-96 target sequence, R6K *ori,* $Amp^r$ | This study |
| pB-iCRISPR-*cbuA0031* -sgRNA-97 | pB-iCRISPR containing *cbuA0031* -sgRNA-97 target sequence, R6K *ori* , $Amp^r$ | This study |
| pB-iCRISPR-*cbuA0031* -sgRNA-98 | pB-iCRISPR containing *cbuA0031* -sgRNA-98 target sequence, R6K *ori* , $Amp^r$ | This study |
| pB-iCRISPR-*cbuA0032* -sgRNA-99 | pB-iCRISPR containing *cbuA0032* -sgRNA-99 target sequence, R6K *ori* , $Amp^r$ | This study |
| pB-iCRISPR-*cbuA0032* -sgRNA-100 | pB-iCRISPR containing *cbuA0032* -sgRNA-100 target sequence, R6K *ori* , $Amp^r$ | This study |
| pB-iCRISPR-*cbuA0032* -sgRNA-101 | pB-iCRISPR containing *cbuA0032* -sgRNA-101 target sequence, R6K *ori* , $Amp^r$ | This study |
| pB-iCRISPR-*cbuA0033* -sgRNA-102 | pB-iCRISPR containing *cbuA0033* -sgRNA-102 target sequence, R6K *ori* , $Amp^r$ | This study |
| pB-iCRISPR-*cbuA0033* -sgRNA-103 | pB-iCRISPR containing *cbuA0033* -sgRNA-103 target sequence, R6K *ori* , $Amp^r$ | This study |
| pB-iCRISPR-*cbuA0033* -sgRNA-104 | pB-iCRISPR containing *cbuA0033* -sgRNA-104 target sequence, R6K *ori* , $Amp^r$ | This study |
| pB-iCRISPR-*cbuA0034* -sgRNA-105 | pB-iCRISPR containing *cbuA0034* -sgRNA-105 target sequence, R6K *ori* , $Amp^r$ | This study |
| pB-iCRISPR-*cbuA0034* -sgRNA-106 | pB-iCRISPR containing *cbuA0034* -sgRNA-106 target sequence, R6K *ori* , $Amp^r$ | This study |
| pB-iCRISPR-*cbuA0034* -sgRNA-107 | pB-iCRISPR containing *cbuA0034* -sgRNA-107 target sequence, R6K *ori* , $Amp^r$ | This study |
| pB-iCRISPR-*cbuA0034a* -sgRNA-108 | pB-iCRISPR containing *cbuA0034a* -sgRNA-108 target sequence, R6K *ori* , $Amp^r$ | This study |
| pB-iCRISPR-*cbuA0034a* -sgRNA-109 | pB-iCRISPR containing *cbuA0034a* -sgRNA-109 target sequence, R6K *ori* , $Amp^r$ | This study |
| pB-iCRISPR-*cbuA0034a* -sgRNA-110 | pB-iCRISPR containing *cbuA0034a* -sgRNA-110 target sequence, R6K *ori* , $Amp^r$ | This study |
| pB-iCRISPR-*cbuA0036* -sgRNA-111 | pB-iCRISPR containing *cbuA0036* -sgRNA-111 target sequence, R6K *ori* , $Amp^r$ | This study |
| pB-iCRISPR-*cbuA0036* -sgRNA-112 | pB-iCRISPR containing *cbuA0036* -sgRNA-112 target sequence, R6K *ori* , $Amp^r$ | This study |
| pB-iCRISPR-*cbuA0036* -sgRNA-113 | pB-iCRISPR containing *cbuA0036* -sgRNA-113 target sequence, R6K *ori* , $Amp^r$ | This study |
| pB-iCRISPR-*cbuA0036* -sgRNA-114 | pB-iCRISPR containing *cbuA0036* -sgRNA-114 target sequence, R6K *ori* , $Amp^r$ | This study |
| pB-iCRISPR-*cbuA0036* -sgRNA-115 | pB-iCRISPR containing *cbuA0036* -sgRNA-115 target sequence, R6K *ori* , $Amp^r$ | This study |
| pB-iCRISPR-*cbuA0037* -sgRNA-116 | pB-iCRISPR containing *cbuA0037* -sgRNA-116 target sequence, R6K *ori* , $Amp^r$ | This study |
| pB-iCRISPR-*cbuA0037* -sgRNA-117 | pB-iCRISPR containing *cbuA0037* -sgRNA-117 target sequence, R6K *ori* , $Amp^r$ | This study |
| pB-iCRISPR-*cbuA0037* -sgRNA-118 | pB-iCRISPR containing *cbuA0037* -sgRNA-118 target sequence, R6K *ori* , $Amp^r$ | This study |
| pB-iCRISPR-*cbuA0038* -sgRNA-119 | pB-iCRISPR containing *cbuA0038* -sgRNA-119 target sequence, R6K ori, $Amp^r$ | This study |
| pB-iCRISPR-*cbuA0038* -sgRNA-120 | pB-iCRISPR containing *cbuA0038* -sgRNA-120 target sequence, R6K *ori* , $Amp^r$ | This study |
| pB-iCRISPR-*cbuA0038* -sgRNA-121 | pB-iCRISPR containing *cbuA0038* -sgRNA-121 target sequence, R6K *ori* , $Amp^r$ | This study |
| pB-iCRISPR-*cbuA0039* -sgRNA-122 | pB-iCRISPR containing *cbuA0039* -sgRNA-122 target sequence, R6K *ori,* $Amp^r$ | This study |
| pB-iCRISPR-*cbuA0039* -sgRNA-123 | pB-iCRISPR containing *cbuA0039* -sgRNA-123 target sequence, R6K *ori,* $Amp^r$ | This study |
| pB-iCRISPR-*cbuA0039* -sgRNA-124 | pB-iCRISPR containing *cbuA0039* -sgRNA-124 target sequence, R6K *ori* , $Amp^r$ | This study |
| pB-iCRISPR-*cbuA0039* -sgRNA-125 | pB-iCRISPR containing *cbuA0039* -sgRNA-125 target sequence, R6K *ori* , $Amp^r$ | This study |
| pB-iCRISPR-*cbuA0039* -sgRNA-126 | pB-iCRISPR containing *cbuA0039* -sgRNA-126 target sequence, R6K *ori* , $Amp^r$ | This study |

| | | |
|---|---|---|
| pB-iCRISPR-*cbuA0039a* -sgRNA127 | pB-iCRISPR containing *cbuA0039a* -sgRNA127 target sequence, R6K *ori* , $Amp^r$ | This study |
| pB-iCRISPR-*cbuA0039a* -sgRNA128 | pB-iCRISPR containing *cbuA0039a* -sgRNA128 target sequence, R6K *ori* , $Amp^r$ | This study |
| pB-iCRISPR-*cbuA0039a* -sgRNA129 | pB-iCRISPR containing *cbuA0039a* -sgRNA129 target sequence, R6K *ori* , $Amp^r$ | This study |
| *CRISPRi complementation vectors* | | |
| pJB-*lysCA* | *lysCA* driven by *cbu1169* promoter replacing *cat* in pJB-CAT, $Amp^r$ | Beare *et_al* 2018 |
| pJB-*lysCA-tetRA* | *tetR* repressor and *tetA* promoter cloned into pJB-*lysCA* , $Amp^r$ | This study |
| pJB-*lysCA-tetRA -icmD* -CO | pJB-*lysCA* containing codon optimized *cbu1624* , $Amp^r$ | This study |
| pJB-*lysCA-tetRA-cbuA0027* -CO | pJB-*lysCA* containing codon optimized *cbuA0027* , $Amp^r$ | This study |
| *cbuA0027, cbuA0028 and cbuA0028G C. burnetii expression vectors* | | |
| pMiniTn7T-*argGH* | Arginine-based nutritional selection vector, R6K *ori* , $Amp^r$ | This study |
| pMiniTn7T-*argGH-lacIO* | pMiniTn7T-*argGH* containing lacI repressor and *lacO* operator from pJMP1356, R6K *ori* , $Amp^r$ | This study |
| pMiniTn7T-*argGH-lacIO-cbuA0027* | pMiniTn7T-*argGH-lacIO* containing *cbuA0027* cloned into the unique AvaI restriction site, R6K *ori* , $Amp^r$ | This study |
| pMiniTn7T-CAT | R6K *ori* , $Cm^r$ $Amp^r$ | Beare *et_al* 2011 |
| pMiniTn7T-*proBA* | *$1169^P$ -proBA* cloned into pMiniTn7T-CAT, replacing *cat* , proline-based nutritional selection vector, R6K *ori* , $Amp^r$ | This study |
| pMiniTn7T-*proBA -tetRA* | *tetR* repressor and *tetA* promoter cloned into pMiniTn7T-*proBA* , R6K *ori* , $Amp^r$ | This study |
| pMiniTn7T-*proBA-tetRA-cbuA0028* | *cbuA0028* from NMII cloned into pMiniTn7T-*proBA-tetRA* , R6K *ori* , $Amp^r$ | This study |
| pMiniTn7T-*proBA-tetRA-cbuA0028* G | *cbuA0028* from the G Q212 strain cloned into pMiniTn7T-*proBA-tetRA* , R6K *ori* , $Amp^r$ | This study |
| *cbuA0027, cbuA0028 and cbu0665 cell-free expression vectors* | | |
| pET28a(+) | *E. coli* expression vector with C-terminal V5 and 6xHis tag, pMB1 *ori* , $Km^r$ | Millipore Sigma |
| pET28a(+)-*cbuA0028* | *cbuA0028* -V5 cloned into pET28a(+), pMB1 *ori* , $Km^r$ | This study |
| pDEST15 | N-terminal GST tagged vector, $Cm^r$, $Amp^r$ | Invitrogen |
| pDEST15-*cbuA0027* | *cbuA0027* cloned into pDEST15, $Amp^r$ | This study |
| pEXP1-DEST | N-terminal XpressT and 6xHis tagged vector, $Cm^r$, $Amp^r$ | Invitrogen |
| pEXP1-*cbu0665* | *cbu0665* cloned into pEXP1-DEST, $Amp^r$ | This study |
| *cbuA0028 and cbuA0027 promoter analysis vectors* | | |
| pMiniTn7T-*lysCA* | Lysine-based nutritional selection vector, R6K *ori* , $Amp^r$ | This study |
| pmScarlet-i_C1 | Mammalian expression vector of *mScarlet-i* , $Km^r$ | Bindels *et_al* 2016 |
| pMiniTn7T-*lysCA* -NoP-*mScarlet-i* | Promoterless *mScarlet-i* cloned into pMiniTn7T-*lysCA* , R6K *ori* , $Amp^r$ | This study |
| pMiniTn7T-*lysCA* -P*cbuA0027* -Frag-1-*mScarlet-i* | Frag-1 *cbuA0027* promoter fused to *mScarlet-i* and cloned into pMiniTn7T-*lysCA* , R6K *ori* , $Amp^r$ | This study |
| pMiniTn7T-*lysCA* -P*cbuA0027* -Frag-2-*mScarlet-i* | Frag-2 *cbuA0027* promoter fused to *mScarlet-i* and cloned into pMiniTn7T-*lysCA* , R6K *ori* , $Amp^r$ | This study |
| pMiniTn7T-*lysCA* -P*cbuA0027* -Frag-3-*mScarlet-i* | Frag-3 *cbuA0027* promoter fused to *mScarlet-i* and cloned into pMiniTn7T-*lysCA* , R6K *ori* , $Amp^r$ | This study |
| pMiniTn7T-*lysCA* -P*cbuA0027* -Frag-4-*mScarlet-i* | Frag-4 *cbuA0027* promoter fused to *mScarlet-i* and cloned into pMiniTn7T-*lysCA* , R6K *ori* , $Amp^r$ | This study |
| pMiniTn7T-*lysCA* -P*cbuA0028* -*mScarlet-i* | *cbuA0028* promoter fused to *mScarlet-i* and cloned into pMiniTn7T-*lysCA* , R6K *ori* , $Amp^r$ | This study |

| Primer name | Sequence (5' - 3') |
|---|---|
| *Coxiella shuttle vector construction* | |
| P1169-CAT-SacB-forQpH1plasmid-F | CTGATGCGGTATTTTGGACAATGTTTTTCTACCAAAGC |
| P1169-CAT-SacB-forQpH1plasmid-R | TTATTTGTTAACTGTTAATTGTCCTTG |
| P1169-TyrB-forQpH1plasmid-F | ACAGTTAACAAATAAATGGCTTCGTTTCGCAG |
| P1169-TyrB-forQpH1plasmid-R | TTACATCACCGCAGCAAAC |
| pBR322ori-forQpH1plasmid-F | GCTGCGGTGATGTAACCTGCAGGTTAACGTGAGTTTTCGTTCCA |
| pBR322ori-forQpH1plasmid-R | AAAATACCGCATCAGGCG |
| QpH1ori-forpBR322-F | TGCGGTGATGTAACCTGCAGGTTCTATAGAAGGCTTTGCTAAATC |
| QpH1ori-forpBR322-R | AAACTCACGTTAACCTATGGATAAAATCAAATCAATTC |
| *PCR analysis of ΔQpH1* | |
| CBUA0023-F | TTACTCAATGGAATTCAAGATATGGTGAAGGAAATTTAAC |
| CBUA0023-R | GCTTCTCGAGGAATTCAGATATTGAGGAGGATTTTAGATTC |
| CAT-detection-F | ATGGAGAAAAAAATCACTGGATATACCACC |
| CAT-detection-R | TTACGCCCCGCCCTGCCACTCATCGC |
| *CRISPRi primers* | |
| proBA-forpJMP-F | CTGTTGAACTCTCGAGATGGCTTCGTTTCGCAGC |
| proBA-BsaIfix-F | TTGTCTCAAAAGAATGTGATTATC |
| proBA-BsaIfix-R | ATTCTTTTGAGACAACGTCTCCACAAGTTCTGG |
| proBA-forpJMP-R1 | CTAAACAAAAAACCCTGTCACTTTTTACAGTGACAGGGTTTTTAATATATCAATCTCTAATTTGTCCTG |
| proBA-forpJMP-R2 | GAAGCTGATGCTCGAGCTAAACAAAAAACCCTGTCACTTTTTAC |
| CBU1624-sgRNA-3-F | tagtATAACACCAAAAGCGGACCT |
| CBU1624-sgRNA-3-R | aaacAGGTCCGCTTTTGGTGTTAT |
| CBU1624-sgRNA-5-F | tagtACTCCAGATAACACCAAAAG |
| CBU1624-sgRNA-5-R | aaacCTTTTGGTGTTATCTGGAGT |
| CBU1267a-sgRNA-9-F | tagtTTGGACATTTTGTCTTTCCA |
| CBU1267a-sgRNA-9-R | aaacTGGAAAGACAAAATGTCCAA |
| CBUA0001-sgRNA-1-F | tagtCCGCTGATCGAGCTGTTTTA |
| CBUA0001-sgRNA-1-R | aaacTAAAACAGCTCGATCAGCGG |
| CBUA0001-sgRNA-2-F | tagtATGGATGCCCTGTTCTTTGA |
| CBUA0001-sgRNA-2-R | aaacTCAAAGAACAGGGCATCCAT |
| CBUA0001-sgRNA-3-F | tagtAATGGATGCCCTGTTCTTTG |
| CBUA0001-sgRNA-3-R | aaacCAAAGAACAGGGCATCCATT |
| CBUA0003-sgRNA-4-F | tagtCACTTTTTTGACAAAATCAA |
| CBUA0003-sgRNA-4-R | aaacTTGATTTTGTCAAAAAAGTG |
| CBUA0003-sgRNA-5-F | tagtGCACTTTTTTGACAAAATCA |
| CBUA0003-sgRNA-5-R | aaacTGATTTTGTCAAAAAAGTGC |
| CBUA0003-sgRNA-6-F | tagtACGTTTTCCCGAACATTCGG |
| CBUA0003-sgRNA-6-R | aaacCCGAATGTTCGGGAAAACGT |
| CBUA0003a-sgRNA-7-F | tagtTTCATTTGCTGCCAATGCGT |
| CBUA0003a-sgRNA-7-R | aaacACGCATTGGCAGCAAATGAA |
| CBUA0003a-sgRNA-8-F | tagtCAGCCGCCTCATACAGTTTC |
| CBUA0003a-sgRNA-8-R | aaacGAAACTGTATGAGGCGGCTG |
| CBUA0003a-sgRNA-9-F | tagtAGAGTCCTCAATTGACAGGC |
| CBUA0003a-sgRNA-9-R | aaacGCCTGTCAATTGAGGACTCT |
| CBUA0005-sgRNA-10-F | tagtAACGTGTTCTTAGTAGCTAT |
| CBUA0005-sgRNA-10-R | aaacATAGCTACTAAGAACACGTT |
| CBUA0005-sgRNA-11-F | tagtTCGCGTGTTTTCACTGGTGG |
| CBUA0005-sgRNA-11-R | aaacCCACCAGTGAAAACACGCGA |
| CBUA0005-sgRNA-12-F | tagtCAATCGCGTGTTTTCACTGG |
| CBUA0005-sgRNA-12-R | aaacCCAGTGAAAACACGCGATTG |
| CBUA0006-sgRNA-13-F | tagtAAACCAGCTGTTTGGCTAGA |
| CBUA0006-sgRNA-13-R | aaacTCTAGCCAAACAGCTGGTTT |
| CBUA0006-sgRNA-14-F | tagtTTGCAAACAAACCAGCTGTT |
| CBUA0006-sgRNA-14-R | aaacAACAGCTGGTTTGTTTGCAA |

| | |
|---|---|
| CBUA0006-sgRNA-15-F | tagtATCATGATATCCTTTAAGGA |
| CBUA0006-sgRNA-15-R | aaacTCCTTAAAGGATATCATGAT |
| CBUA0007-sgRNA-16-F | tagtAATAGAAGTTAGAAAAGTAT |
| CBUA0007-sgRNA-16-R | aaacATACTTTTCTAACTTCTATT |
| CBUA0007-sgRNA-17-F | tagtCAATAGAAGTTAGAAAAGTA |
| CBUA0007-sgRNA-17-R | aaacTACTTTTCTAACTTCTATTG |
| CBUA0007-sgRNA-18-F | tagtGAATGATTCAGCACGCTGTC |
| CBUA0007-sgRNA-18-R | aaacGACAGCGTGCTGAATCATTC |
| CBUA0008-sgRNA-19-F | tagtGGTCCGCTATAAAAGGGGGG |
| CBUA0008-sgRNA-19-R | aaacCCCCCCTTTTATAGCGGACC |
| CBUA0008-sgRNA-20-F | tagtTGTGGTCCGCTATAAAAGGG |
| CBUA0008-sgRNA-20-R | aaacCCCTTTTATAGCGGACCACA |
| CBUA0008-sgRNA-21-F | tagtATGTGGTCCGCTATAAAAGG |
| CBUA0008-sgRNA-21-R | aaacCCTTTTATAGCGGACCACAT |
| CBUA0008a-sgRNA-22-F | tagtGCTTGGTTGGTAGGATGTCC |
| CBUA0008a-sgRNA-22-R | aaacGGACATCCTACCAACCAAGC |
| CBUA0008a-sgRNA-23-F | tagtTAAATCCTTGCTTGGTTGGT |
| CBUA0008a-sgRNA-23-R | aaacACCAACCAAGCAAGGATTTA |
| CBUA0008a-sgRNA-24-F | tagtAGCGTAAATCCTTGCTTGGT |
| CBUA0008a-sgRNA-24-R | aaacACCAAGCAAGGATTTACGCT |
| CBUA0008b-sgRNA-25-F | tagtAAATTTGTCGCTTGGGTATA |
| CBUA0008b-sgRNA-25-R | aaacTATACCCAAGCGACAAATTT |
| CBUA0008b-sgRNA-26-F | tagtACTGATAAAATTTGTCGCTT |
| CBUA0008b-sgRNA-26-R | aaacAAGCGACAAATTTTATCAGT |
| CBUA0008b-sgRNA-27-F | tagtCACTGATAAAATTTGTCGCT |
| CBUA0008b-sgRNA-27-R | aaacAGCGACAAATTTTATCAGTG |
| CBUA0010-sgRNA-28-F | tagtAGTGGTACTGTTTTCATCAA |
| CBUA0010-sgRNA-28-R | aaacTTGATGAAAACAGTACCACT |
| CBUA0010-sgRNA-29-F | tagtATAGAGTCAAATAGTGCTAG |
| CBUA0010-sgRNA-29-R | aaacCTAGCACTATTTGACTCTAT |
| CBUA0010-sgRNA-30-F | tagtAGTCCCTTGGCTACCTTTGT |
| CBUA0010-sgRNA-30-R | aaacACAAAGGTAGCCAAGGGACT |
| CBUA0011-sgRNA-31-F | tagtAGCGGTGCACCTGAGTCCAA |
| CBUA0011-sgRNA-31-R | aaacTTGGACTCAGGTGCACCGCT |
| CBUA0011-sgRNA-32-F | tagtTTTTCTTGGGCGACTTTAAG |
| CBUA0011-sgRNA-32-R | aaacCTTAAAGTCGCCCAAGAAAA |
| CBUA0011-sgRNA-33-F | tagtACTATGTCCAGCATTTTCTT |
| CBUA0011-sgRNA-33-R | aaacAAGAAAATGCTGGACATAGT |
| CBUA0012-sgRNA-34-F | tagtGCCCCACAGAAGTCTTGCCG |
| CBUA0012-sgRNA-34-R | aaacCGGCAAGACTTCTGTGGGGC |
| CBUA0012-sgRNA-35-F | tagtAAATAACTTCTCGGCATAAT |
| CBUA0012-sgRNA-35-R | aaacATTATGCCGAGAAGTTATTT |
| CBUA0012-sgRNA-36-F | tagtGCATTCCCAAAATAACTTCT |
| CBUA0012-sgRNA-36-R | aaacAGAAGTTATTTTGGGAATGC |
| CBUA0013-sgRNA-37-F | tagtTTCGGTAGTGTAAAAAAATA |
| CBUA0013-sgRNA-37-R | aaacTATTTTTTTACACTACCGAA |
| CBUA0013-sgRNA-38-F | tagtTTTTCTTTTATGTCAATTTT |
| CBUA0013-sgRNA-38-R | aaacAAAATTGACATAAAAGAAAA |
| CBUA0013-sgRNA-39-F | tagtTTAAAACTTTTTCTGGAGTT |
| CBUA0013-sgRNA-39-R | aaacAACTCCAGAAAAAGTTTTAA |
| CBUA0013a-sgRNA-40-F | tagtAGGCTGAATTCTACACGTCA |
| CBUA0013a-sgRNA-40-R | aaacTGACGTGTAGAATTCAGCCT |
| CBUA0013a-sgRNA-41-F | tagtCAGGCTGAATTCTACACGTC |
| CBUA0013a-sgRNA-41-R | aaacGACGTGTAGAATTCAGCCTG |
| CBUA0013a-sgRNA-42-F | tagtTGCTTGCGATCGAGATCCTC |
| CBUA0013a-sgRNA-42-R | aaacGAGGATCTCGATCGCAAGCA |
| CBUA0014-sgRNA-43-F | tagtGGACACATCTTCATTCGTTT |
| CBUA0014-sgRNA-43-R | aaacAAACGAATGAAGATGTGTCC |
| CBUA0014-sgRNA-44-F | tagtTAAATGATCCCTTTCAGAAG |
| CBUA0014-sgRNA-44-R | aaacCTTCTGAAAGGGATCATTTA |

| | |
|---|---|
| CBUA0014-sgRNA-45-F | tagtATAAATGATCCCTTTCAGAA |
| CBUA0014-sgRNA-45-R | aaacTTCTGAAAGGGATCATTTAT |
| CBUA0015-sgRNA-46-F | tagtACCGTGCAATTTTTAAATTC |
| CBUA0015-sgRNA-46-R | aaacGAATTTAAAAATTGCACGGT |
| CBUA0015-sgRNA-47-F | tagtAATGATTAAACCGGCCCGGT |
| CBUA0015-sgRNA-47-R | aaacACCGGGCCGGTTTAATCATT |
| CBUA0015-sgRNA-48-F | tagtCAATGATTAAACCGGCCCGG |
| CBUA0015-sgRNA-48-R | aaacCCGGGCCGGTTTAATCATTG |
| CBUA0016-sgRNA-49-F | tagtCTTCTTTTGAGTGAATTTCT |
| CBUA0016-sgRNA-49-R | aaacAGAAATTCACTCAAAAGAAG |
| CBUA0016-sgRNA-50-F | tagtTTGAATCTTGTGGTAAGCTA |
| CBUA0016-sgRNA-50-R | aaacTAGCTTACCACAAGATTCAA |
| CBUA0016-sgRNA-51-F | tagtTTTGAATCTTGTGGTAAGCT |
| CBUA0016-sgRNA-51-R | aaacAGCTTACCACAAGATTCAAA |
| CBUA0017-sgRNA-52-F | tagtCGCAGAACGCATAGAGGAAA |
| CBUA0017-sgRNA-52-R | aaacTTTCCTCTATGCGTTCTGCG |
| CBUA0017-sgRNA-53-F | tagtGCGCAGAACGCATAGAGGAA |
| CBUA0017-sgRNA-53-R | aaacTTCCTCTATGCGTTCTGCGC |
| CBUA0017-sgRNA-54-F | tagtACAGCGCGCAGAACGCATAG |
| CBUA0017-sgRNA-54-R | aaacCTATGCGTTCTGCGCGCTGT |
| CBUA0018-sgRNA-55-F | tagtAACCATCGTATCCTTTTTTA |
| CBUA0018-sgRNA-55-R | aaacTAAAAAAGGATACGATGGTT |
| CBUA0018-sgRNA-56-F | tagtGATCTTCATAGGGGCGAGGG |
| CBUA0018-sgRNA-56-R | aaacCCCTCGCCCCTATGAAGATC |
| CBUA0018-sgRNA-57-F | tagtATTGATCTTCATAGGGGCGA |
| CBUA0018-sgRNA-57-R | aaacTCGCCCCTATGAAGATCAAT |
| CBUA0020-sgRNA-58-F | tagtGCTCAATCAATACTGCATTT |
| CBUA0020-sgRNA-58-R | aaacAAATGCAGTATTGATTGAGC |
| CBUA0020-sgRNA-59-F | tagtATGGGGGGTAATTCTTTTGG |
| CBUA0020-sgRNA-59-R | aaacCCAAAAGAATTACCCCCCAT |
| CBUA0020-sgRNA-60-F | tagtAAAATGGGGGGTAATTCTTT |
| CBUA0020-sgRNA-60-R | aaacAAAGAATTACCCCCCATTTT |
| CBUA0021-sgRNA-61-F | tagtACAGTAGGATTATTAGGAGG |
| CBUA0021-sgRNA-61-R | aaacCCTCCTAATAATCCTACTGT |
| CBUA0021-sgRNA-62-F | tagtCTGACAGTAGGATTATTAGG |
| CBUA0021-sgRNA-62-R | aaacCCTAATAATCCTACTGTCAG |
| CBUA0021-sgRNA-63-F | tagtCTACTGACAGTAGGATTATT |
| CBUA0021-sgRNA-63-R | aaacAATAATCCTACTGTCAGTAG |
| CBUA0022-sgRNA-64-F | tagtAGGCGCATTGTAGGCAAAAA |
| CBUA0022-sgRNA-64-R | aaacTTTTTGCCTACAATGCGCCT |
| CBUA0022-sgRNA-65-F | tagtTAGGCGCATTGTAGGCAAAA |
| CBUA0022-sgRNA-65-R | aaacTTTTGCCTACAATGCGCCTA |
| CBUA0022-sgRNA-66-F | tagtTAAGTCCATAGGCGCATTGT |
| CBUA0022-sgRNA-66-R | aaacACAATGCGCCTATGGACTTA |
| CBUA0023-sgRNA-67-F | tagtATGTGCTTTTTCAACACTAT |
| CBUA0023-sgRNA-67-R | aaacATAGTGTTGAAAAAGCACAT |
| CBUA0023-sgRNA-68-F | tagtAGGGAAGGATTGGTTTTCCG |
| CBUA0023-sgRNA-68-R | aaacCGGAAAACCAATCCTTCCCT |
| CBUA0023-sgRNA-69-F | tagtTAGGGAAGGATTGGTTTTCC |
| CBUA0023-sgRNA-69-R | aaacGGAAAACCAATCCTTCCCTA |
| CBUA0023a-sgRNA-70-F | tagtTTATACTGAGAACCAAGGTT |
| CBUA0023a-sgRNA-70-R | aaacAACCTTGGTTCTCAGTATAA |
| CBUA0023a-sgRNA-71-F | tagtGCTTTTTATACTGAGAACCA |
| CBUA0023a-sgRNA-71-R | aaacTGGTTCTCAGTATAAAAAGC |
| CBUA0024-sgRNA-72-F | tagtCGATAGGGCCAATGGGGATT |
| CBUA0024-sgRNA-72-R | aaacAATCCCCATTGGCCCTATCG |
| CBUA0024-sgRNA-73-F | tagtCGGGATCGATAGGGCCAATG |
| CBUA0024-sgRNA-73-R | aaacCATTGGCCCTATCGATCCCG |
| CBUA0024-sgRNA-74-F | tagtTCGGGATCGATAGGGCCAAT |
| CBUA0024-sgRNA-74-R | aaacATTGGCCCTATCGATCCCGA |

| | |
|---|---|
| CBUA0025-sgRNA-75-F | tagtAAGAATTTCACCCCATCATT |
| CBUA0025-sgRNA-75-R | aaacAATGATGGGGTGAAATTCTT |
| CBUA0025-sgRNA-76-F | tagtTTTTTGGTTTTTTACCACCT |
| CBUA0025-sgRNA-76-R | aaacAGGTGGTAAAAAACCAAAAA |
| CBUA0025-sgRNA-77-F | tagtTTTTTTGGTTTTTTACCACC |
| CBUA0025-sgRNA-77-R | aaacGGTGGTAAAAAACCAAAAAA |
| CBUA0026-sgRNA-78-F | tagtGGATTTCAGAGGTTAAAACT |
| CBUA0026-sgRNA-78-R | aaacAGTTTTAACCTCTGAAATCC |
| CBUA0026-sgRNA-79-F | tagtTAGGCATACAGGGATTTCAG |
| CBUA0026-sgRNA-79-R | aaacCTGAAATCCCTGTATGCCTA |
| CBUA0026-sgRNA-80-F | tagtGTTTAAGTTTTAGGCATACA |
| CBUA0026-sgRNA-80-R | aaacTGTATGCCTAAAACTTAAAC |
| CBUA0026a-sgRNA-81-F | tagtCATATACACACATCAACACT |
| CBUA0026a-sgRNA-81-R | aaacAGTGTTGATGTGTGTATATG |
| CBUA0026a-sgRNA-82-F | tagtTTTTGCCGACTCAGACTCAA |
| CBUA0026a-sgRNA-82-R | aaacTTGAGTCTGAGTCGGCAAAA |
| CBUA0026a-sgRNA-83-F | tagtCAGATTGTTTAATTTTTTCT |
| CBUA0026a-sgRNA-83-R | aaacAGAAAAAATTAAACAATCTG |
| CBUA0027-sgRNA-84-F | tagtACGGCCTCAATGGGCGATAA |
| CBUA0027-sgRNA-84-R | aaacTTATCGCCCATTGAGGCCGT |
| CBUA0027-sgRNA-85-F | tagtGATTCATGAACGGCCTCAAT |
| CBUA0027-sgRNA-85-R | aaacATTGAGGCCGTTCATGAATC |
| CBUA0027-sgRNA-86-F | tagtAGATTCATGAACGGCCTCAA |
| CBUA0027-sgRNA-86-R | aaacTTGAGGCCGTTCATGAATCT |
| CBUA0028-sgRNA-87-F | tagtTTTCTGATTTTGCCCATCGA |
| CBUA0028-sgRNA-87-R | aaacTCGATGGGCAAAATCAGAAA |
| CBUA0028-sgRNA-88-F | tagtGGCCGTTTTCAGTTGGCTAT |
| CBUA0028-sgRNA-88-R | aaacATAGCCAACTGAAAACGGCC |
| CBUA0028-sgRNA-89-F | tagtCACTAATGGCCGTTTTCAGT |
| CBUA0028-sgRNA-89-R | aaacACTGAAAACGGCCATTAGTG |
| CBUA0029-sgRNA-90-F | tagtGACCATTTAAGAGTGTGGGC |
| CBUA0029-sgRNA-90-R | aaacGCCCACACTCTTAAATGGTC |
| CBUA0029-sgRNA-91-F | tagtTTTTGACCATTTAAGAGTGT |
| CBUA0029-sgRNA-91-R | aaacACACTCTTAAATGGTCAAAA |
| CBUA0029-sgRNA-92-F | tagtTTTTTGACCATTTAAGAGTG |
| CBUA0029-sgRNA-92-R | aaacCACTCTTAAATGGTCAAAAA |
| CBUA0029a-sgRNA-93-F | tagtTGCTTCACGTGAGGCAATCA |
| CBUA0029a-sgRNA-93-R | aaacTGATTGCCTCACGTGAAGCA |
| CBUA0029a-sgRNA-94-F | tagtGGTAAATGTTGCTTCACGTG |
| CBUA0029a-sgRNA-94-R | aaacCACGTGAAGCAACATTTACC |
| CBUA0029a-sgRNA-95-F | tagtTTTTTCAACGACGGATTGAT |
| CBUA0029a-sgRNA-95-R | aaacATCAATCCGTCGTTGAAAAA |
| CBUA0031-sgRNA-96-F | tagtTAAAAGAAAATGGAAATAAC |
| CBUA0031-sgRNA-96-R | aaacGTTATTTCCATTTTCTTTTA |
| CBUA0031-sgRNA-97-F | tagtATGAGTAATATAAAAGAAAA |
| CBUA0031-sgRNA-97-R | aaacTTTTCTTTTATATTACTCAT |
| CBUA0031-sgRNA-98-F | tagtTGGTGGAGAAGGGAGGCGTG |
| CBUA0031-sgRNA-98-R | aaacCACGCCTCCCTTCTCCACCA |
| CBUA0032-sgRNA-99-F | tagtTAAGTGCAAATCACTGACTT |
| CBUA0032-sgRNA-99-R | aaacAAGTCAGTGATTTGCACTTA |
| CBUA0032-sgRNA-100-F | tagtTTAAGTGCAAATCACTGACT |
| CBUA0032-sgRNA-100-R | aaacAGTCAGTGATTTGCACTTAA |
| CBUA0032-sgRNA-101-F | tagtATTGCTGTATCTGCCTCTAG |
| CBUA0032-sgRNA-101-R | aaacCTAGAGGCAGATACAGCAAT |
| CBUA0033-sgRNA-102-F | tagtTTAAGATAGAGTAAATCTTT |
| CBUA0033-sgRNA-102-R | aaacAAAGATTTACTCTATCTTAA |
| CBUA0033-sgRNA-103-F | tagtAACTCGAACCCGGTATCTTT |
| CBUA0033-sgRNA-103-R | aaacAAAGATACCGGGTTCGAGTT |
| CBUA0033-sgRNA-104-F | tagtTCCGCGCATAAACTCGAACC |
| CBUA0033-sgRNA-104-R | aaacGGTTCGAGTTTATGCGCGGA |

| | |
|---|---|
| CBUA0034-sgRNA-105-F | tagtGCTACGATTTTTGTATTTGA |
| CBUA0034-sgRNA-105-R | aaacTCAAATACAAAAATCGTAGC |
| CBUA0034-sgRNA-106-F | tagtTGCTACGATTTTTGTATTTG |
| CBUA0034-sgRNA-106-R | aaacCAAATACAAAAATCGTAGCA |
| CBUA0034-sgRNA-107-F | tagtTATCATCAGAGGATTAGTGT |
| CBUA0034-sgRNA-107-R | aaacACACTAATCCTCTGATGATA |
| CBUA0034a-sgRNA-108-F | tagtAATTTGCCCACTTTAACCTT |
| CBUA0034a-sgRNA-108-R | aaacAAGGTTAAAGTGGGCAAATT |
| CBUA0034a-sgRNA-109-F | tagtAAGCATTTGGAATTGCTCTA |
| CBUA0034a-sgRNA-109-R | aaacTAGAGCAATTCCAAATGCTT |
| CBUA0034a-sgRNA-110-F | tagtCTTCAAAAGCGTAAAGCATT |
| CBUA0034a-sgRNA-110-R | aaacAATGCTTTACGCTTTTGAAG |
| CBUA0036-sgRNA-111-F | tagtAACGAAATCGTTTGTTTTAG |
| CBUA0036-sgRNA-111-R | aaacCTAAAACAAACGATTTCGTT |
| CBUA0036-sgRNA-112-F | tagtACCCGTCTGTAGTAAATCAA |
| CBUA0036-sgRNA-112-R | aaacTTGATTTACTACAGACGGGT |
| CBUA0036-sgRNA-113-F | tagtATTCTTTTATCCAACCCTTT |
| CBUA0036-sgRNA-113-R | aaacCGAAAGGGTTGGATAAAAGA |
| CBUA0036-sgRNA-114-F | tagtTTCTTTTATCCAACCCTTTC |
| CBUA0036-sgRNA-114-R | aaacGAAAGGGTTGGATAAAAGAA |
| CBUA0036-sgRNA-115-F | tagtTCTTTTATCCAACCCTTTCG |
| CBUA0036-sgRNA-115-R | aaacATTCTTTTATCCAACCCTTT |
| CBUA0037-sgRNA-116-F | tagtGGGGGTTTCTGTACCGTAGG |
| CBUA0037-sgRNA-116-R | aaacCCTACGGTACAGAAACCCCC |
| CBUA0037-sgRNA-117-F | tagtCGGGGGTTTCTGTACCGTAG |
| CBUA0037-sgRNA-117-R | aaacCTACGGTACAGAAACCCCCG |
| CBUA0037-sgRNA-118-F | tagtTCGGGGGTTTCTGTACCGTA |
| CBUA0037-sgRNA-118-R | aaacTACGGTACAGAAACCCCCGA |
| CBUA0038-sgRNA-119-F | tagtTTTTTCATTAACATACCTAA |
| CBUA0038-sgRNA-119-R | aaacTTAGGTATGTTAATGAAAAA |
| CBUA0038-sgRNA-120-F | tagtAATTTTCAATTTTTTTAATC |
| CBUA0038-sgRNA-120-R | aaacGATTAAAAAAATTGAAAATT |
| CBUA0038-sgRNA-121-F | tagtTGCTGCTTTATTTAATACGA |
| CBUA0038-sgRNA-121-R | aaacTCGTATTAAATAAAGCAGCA |
| CBUA0039-sgRNA-122-F | tagtCAATCCGGGAGCTCCGGATT |
| CBUA0039-sgRNA-122-R | aaacAATCCGGAGCTCCCGGATTG |
| CBUA0039-sgRNA-123-F | tagtCCAATCCGGGAGCTCCGGAT |
| CBUA0039-sgRNA-123-R | aaacATCCGGAGCTCCCGGATTGG |
| CBUA0039-sgRNA-124-F | tagtAAGCTCCAATCCGGGAGCTC |
| CBUA0039-sgRNA-124-R | aaacGAGCTCCCGGATTGGAGCTT |
| CBUA0039-sgRNA-125-F | tagtTTTGGCTCAAGCTCCAATCC |
| CBUA0039-sgRNA-125-R | aaacGGATTGGAGCTTGAGCCAAA |
| CBUA0039-sgRNA-126-F | tagtATTTGGCTCAAGCTCCAATC |
| CBUA0039-sgRNA-126-R | aaacGATTGGAGCTTGAGCCAAAT |
| CBUA0039a-sgRNA-127-F | tagtGCATTGGGTTTGTGGCGCGT |
| CBUA0039a-sgRNA-127-R | aaacACGCGCCACAAACCCAATGC |
| CBUA0039a-sgRNA-128-F | tagtTTTTTTGGGCATTGGGTTTG |
| CBUA0039a-sgRNA-128-R | aaacCAAACCCAATGCCCAAAAAA |
| CBUA0039a-sgRNA-129-F | tagtCGGTATTTTTTTTGGGCATT |
| CBUA0039a-sgRNA-129-R | aaacAATGCCCAAAAAAAATACCG |
| *CRISPRi complementation vector primers* | |
| TetRA-forpJB-F | ACTGACGCGTGAATTCTTAAGACCCACTTTCACATTTAA |
| TetRA-forpJB-R | TCGTATGGGTACATCTGCAGCTTTTCTCTATCACTGATAGGG |
| CBU1624-CO-F1 | AATACTCTTGGGCCCATTGCTCGTTCTCAGCGGCGTCATCGGCGTATTATTTGCTGGAAC |
| CBU1624-CO-F2 | GTGATAGAGAAAAGCTGCAGATGCAGACTAAAACAATGCGCTTATTTCGAATACTCTTGGGCCCATTGC |
| CBU1624-CO-R | GCATGCCTCAGTCGACTTATGACGAACCAATAATACTGTGG |

| | |
|---|---|
| CBUA0027-CO-F1 | GTGATAGAGAAAAGCTGCAGGTGATGAAAGATAAAAAAAAGAAAAGTAAAACTCGTTTTAAACCTCTGAGCCCAATCG |
| CBUA0027-CO-F2 | TTAAACCTCTGAGCCCAATCGAAGCTGTGCACGAGAGCGCAAAAGGATTATATGATGCCG |
| CBUA0027-CO-R | CCTTCAAGCCATCGCTTAGGTCGACTGAGGCATGC |
| *qRT-PCR probes and primers* | |
| groEL-F | ATGGCTTACCCGCTTTCTC |
| groEL-R | AACGTCTTTATGGTTCAAGACTTT |
| groEL-probe | CalFluorGold-TGTTCAAGCAGCCGTTGTCGC-BHQ1 |
| CBUA0006-F | CGCCAGTGCTCAACACATTC |
| CBUA0006-R | CACTGATTGATGCCTTACAAACCT |
| CBUA0006-probe | FAM-TCAAAGCTCCAACGCTCCACATTAGTTACG-BHQ1 |
| CBUA0013-F | TCATTTTGGTGGTATTAACGGGTAT |
| CBUA0013-R | CTCCACTTCACCCTTGTTTTTACTC |
| CBUA0013-probe | FAM-ATCCTCCAGAATGGAAAATCGCCGC-BHQ1 |
| CBUA0014-F | CCTTCTGAAAGGGATCATTTATCC |
| CBUA0014-R | TGCTCAAATTCCGCACCTACTT |
| CBUA0014-probe | FAM-ATCTTTCTGCGCGGGATTCGTATTCTCT-BHQ1 |
| CBUA0015-F | CCGGGCCGGTTTAATCAT |
| CBUA0015-R | GCTTTTACCACCCCGAAATTC |
| CBUA0015-probe | FAM-CGGAATTGACTGACTGCTCTCCCAGTG-BHQ1 |
| CBUA0016-F | GGATAAAAACCAAACTCTCCTCCTT |
| CBUA0016-R | CCTCATTAGTGGATTCATCTCGATCT |
| CBUA0016-probe | FAM-TCGACCCTTGGAGCCAGGACAATAAA-BHQ1 |
| CBUA0021-F | CTTGCCGGTTTTGATTACTCCTT |
| CBUA0021-R | GCTCGATCACGAAGAGAATATTTCT |
| CBUA0021-probe | FAM-TTTCTAAACACGTTCAATGCAGCCAGCA-BHQ1 |
| CBUA0023-F | TTTTTATGGTTGCGAGGTTGTAT |
| CBUA0023-R | GGTTGAAGTCAGAGCCGGTTAT |
| CBUA0023-probe | FAM-CTTCTCCAACCGAGGAATACCCATCAA-BHQ1 |
| CBUA0025-F | TCACACTCGACTCTCAGCCATT |
| CBUA0025-R | TGACTGACGAAGAAGCAGCATT |
| CBUA0025-probe | FAM-CGATGCGCCAATTTTTTGGTTTTTTACC-BHQ1 |
| CBUA0027-F | GTGCCTCGAGGGTGTTTCTC |
| CBUA0027-R | CCTCGTGAAATTAAGCGCATTC |
| CBUA0027-probe | FAM-TTTGCAAAAACGGCCTGGCTCACT-BHQ1 |
| CBUA0034-F | CGTTCAGTAAACGACCTTGTAAACT |
| CBUA0034-R | CCTCAAATACAAAAATCGTAGCACT |
| CBUA0034-probe | FAM-TTTATCGACATCCAAATTCCCACAGCC-BHQ1 |
| CBU0334-F | AAAAGGCGGTCACTTAATTGGTTC |
| CBU0334-R | GCGCTACTGATTGCTGAGCTT |
| CBU0334-probe | FAM-CATACAAAAATACCCAAAACACGGGTGCG-BHQ1 |
| *cbuA0027, cbuA0028 and cbuA0028G C. burnetii expression vector primers* | |
| ArgGH-forminiTn7-F | GGGGTTCGAGGTCGACTTAGCCTCCTTTTAATAATTCATTCAGTG |
| ArgGH-forminiTn7-R | TATCGATACCGTCGACATGGCTTCGTTTCGCAGCGAAC |
| LacIO-forminiTn7-F | AATGGAATTCCTCGAGCACTGTCAGGTGCGGG |
| LacIO-forminiTn7-R | TGCAAGGCCTCCCGGGTTTCCTGTGTGAAACCTGCTG |
| CBUA0027-forminiTn7-F | ACACAGGAAACCCGGGGTGATGAAAGATAAAAAAAAGAAAAG |
| CBUA0027-forminiTn7-R | TGCAAGGCCTCCCGGGCTAAGCGATGGCTTGAAG |
| ProBA-forminiTn7-F | GGGGTTCGAGGTCGACTCAATCTCTAATTTGTCCCG |
| ProBA-forminiTn7-R | CGCTGCGAAACGAAGCCATGTCGACGGTATCGATA |
| TetRA-forminiTn7-F | AATGGAATTCCTCGAGTTAAGACCCACTTTCACATTTAAG |
| TetRA-forminiTn7-R | GGTTGGCCTGCAAGGCCTCCCGGGCTTTTCTCTATCACTGATAGGG |
| CBUA0028-forminiTn7-F | TAGAGAAAAGCCCGGGATGAGGATATTTAAAACACGCTATTTC |
| CBUA0028-forminiTn7-R | CTGATAGAGGTGCTGTGACCCGGGAGGCCTTGCA |
| CBUA0028G-forminiTn7-F | TAGAGAAAAGCCCGGGATTCTGGGATATTCCTTTATCTATAAAAAAC |

| | |
|---|---|
| CBUA0028G-forminiTn7-R | TGCAAGGCCTCCCGGGTCACAGCACCTCTATCAGTTC |
| *cbuA0027, cbuA0028 and cbu0665 cell-free expression vectors* | |
| CBUA0028-forpET-F | AGGAGATATACCATGAGGATATTTAAAACACGCTATTTC |
| CBUA0028-forpET-R1 | GGCTTACCTTCAAGCTTCGAATTATCAACCACTTTGTACAACAGCACCTCTATCAGTTCG |
| CBUA0028-forpET-R2 | GGTGGTGGTGCTCGAGACGCGTAGAATCGAGACCGAGGAGAGGGTTAGGGATAGGCTTACCTTCAAGCTTCG |
| CBUA0027-forpDEST15-F | CAAACAAGTTTGTACAAAATGAAAGATAAAAAAAAGAAAAGTAAAACTC |
| CBUA0027-forpDEST15-R | CAAACCACTTTGTACACTAAGCGATGGCTTGAAGG |
| CBU0665-forpEXP1-F | CAAACAAGTTTGTACAAAATGCTTTTAAGAGGAATAATAGTG |
| CBU0665-forpEXP1-R | CAGCCGGATCAAGCTTCTATTGAAGTCTGGTTAAACC |
| *EMSA promoter primers* | |
| PCBUA0028-EMSA-F | ATGATCTTGCCAAAGCGGG |
| PCBUA0028-EMSA-R | TTTTCTGATTTTGCCCATCGATG |
| PCBUA0029-EMSA-F | GATCTTAACCAACCTCAGAATC |
| PCBUA0029-EMSA-R | TTTAAGAGTGTGGGCTGG |
| PGroES-EMSA-F | TTTTGCCACCAGCCGTTAATTC |
| PGroES-EMSA-R | TCTTCAAGGCGACGGAC |
| *cbuA0028 and cbuA0027 promoter analysis vectors* | |
| LysCA-forminiTn7-F | GGGGTTCGAGGTCGACTTATTCCAGAACTATTTCCTGAG |
| LysCA-forminiTn7-R | TATCGATACCGTCGACATGGCTTCGTTTCGCAGCG |
| NoP-mScarlet-i-F | TTACTCAATGGAATTCATGGTGAGCAAGGGCGAG |
| NoP-mScarlet-i-R | GCCCAAGCTTCTCGAGCTAAACAAAAAACCCTGTCACTTTTTAC |
| PCBUA0027-Frag1-F | TTACTCAATGGAATTCATGAGGATATTTAAAACACGCTATTTC |
| PCBUA0027-Frag1-R | TGCTCACCATAAGCTTCCTCTATCAGTTCGCCTATTTTTAATAATG |
| PCBUA0027-Frag2-F | TTACTCAATGGAATTCTACACGATGGAGATTTAGGTTC |
| PCBUA0027-Frag2-R | TGCTCACCATAAGCTTCCTCTATCAGTTCGCCTATTTTTAATAATG |
| PCBUA0027-Frag3-F | TTACTCAATGGAATTCAGAAGATAAAGCTTTTTTTGTATATGGTTATG |
| PCBUA0027-Frag3-R | TGCTCACCATAAGCTTCCTCTATCAGTTCGCCTATTTTTAATAATG |
| mScarlet-i-forPCBUA0027-F | AAGCTTATGGTGAGCAAGGGCGAG |
| mScarlet-i-forPCBUA0027-F | GCCCAAGCTTCTCGAGCTAAACAAAAAACCCTGTCACTTTTTAC |
| PCBUA0027-Frag4-mScarlet-i-F1 | GATATGAAAGAATTAGAGCTTCAATCATTATTAAAAATAGGCGAACTGATAGAGGAAGCTTATGGTGAGCAAGGGCGAG |
| PCBUA0027-Frag4-mScarlet-i-F2 | TTACTCAATGGAATTCGTACAAAAAGTTGTCAAAAACTTATTTTGATATGAAAGAATTAGAGCTTCAATC |
| PCBUA0027-Frag3-mScarlet-i-R | GCCCAAGCTTCTCGAGCTAAACAAAAAACCCTGTCACTTTTTAC |
| PCBUA0028-mScarlet-i-F1 | GTACAGATTGACAAATAGTTCACTTAGTGATATAGTCTTAACAATAAGGGAAAGCTTATGGTGAGCAAGGGCGAG |
| PCBUA0028-mScarlet-i-F2 | TTACTCAATGGAATTCCTGATGGAAATCTGCTCGAGATAGGCAATGAGTACAGATTGACAAATAGTTCACTTAG |
| PCBUA0028-mScarlet-i-R | GCCCAAGCTTCTCGAGCTAAACAAAAAACCCTGTCACTTTTTAC |
