## Supplemental File 1 for "A toxin-antitoxin system ensures plasmid stability in *Coxiella burnetii*"

**Supplemental file 1** – Construction of plasmids used in this study.

Construction of pB-TyrB-QpH1ori

PCR was carried out to amplify the pBR322 origin of replication, *tyrB* gene from *E. coli*, and *1169*$^{P}$-CAT-*sacB* fragment. These fragments were cloned using In-Fusion HD to create pB-TyrB. The QpH1 replication machinery and origin of replication (*cbuA00036*-*cbuA0039a*) was amplified from the QpH1 plasmid and cloned into pB-TyrB digested with SbfI using In-Fusion HD to create pB-TyrB-QpH1ori.

Construction of CRISPRi plasmids

The *proBA* genes were amplified from *Legionella pneumophilia* JR32 genomic DNA (gDNA) and fused to the *cbu1169* promoter by PCR. The *1169*$^{P}$*-proBA* fragment was cloned into AvaI digested pJMP1356 by In-Fusion HD to create pB-CRISPRi. Single guide RNA (sgRNA) targeting sequences were generated by annealing two complementary 24 bp oligonucleotides, resulting in a 20 bp double stranded DNA (dsDNA) fragment with a 4 bp BsaI-compatible overhang on the 5′ ends. The sgRNA target dsDNA sequences were cloned into pB-CRISPRi digested with BsaI using the NEBridge Golden Gate Assembly kit (BsaI-HF®v2) kit.

Construction of CRISPRi complementation vectors

The targeting regions chosen for CRISPRi knockdown of *cbu1624* and *cbuA0027* were codon optimized using overlapping oligonucleotides to add silent mutations to the *cbu1624* (*cbu1624*-CO) and *cbuA0027* (*cbuA0027*-CO) sequences. The *cbu1624*-CO and *cbuA0027*-CO codon optimized genes were amplified by PCR and cloned into pJB-*lysCA-tetRA* digested with PstI/SalI to create pJB-*lysCA-tetRA-cbu1624*-CO and pJB-*tetRA-cbuA0027*-CO, respectively, using In-Fusion HD.

Construction of *cbuA0027* and *cbuA0028 C. burnetii* expression vectors

The *cbuA0027* and *cbuA0028* or *cbuA0028G* genes were amplified by PCR from *C. burnetii* NMII or G Q212 gDNA, respectively. The *cbuA0027* PCR fragment was then cloned into pMiniTn7T-*argGH-lacIO* digested with SmaI using In-Fusion HD to create pMiniTn7T-*argGH-lacIO-cbuA0027*. The *cbuA0028* and *cbuA0028G* PCR products were cloned into pMiniTn7T-proBA-TetRA digested with SmaI to create pMiniTn7T-*proBA-tetRA-cbuA0028* and pMiniTn7T-*proBA-tetRA-cbuA0028*G.

Construction of *cbuA0027*, *cbuA0028* and *cbu0665* cell-free expression vectors

The *cbuA0027* and *cbu0665* genes were amplified by PCR from NMII gDNA and the resulting PCR products were cloned into BsrGI-digested pDEST15 or pEXP1-DEST to create pDEST15-*cbuA0027* and pEXP1-*cbu0665*, respectively. The *cbuA0028* gene was amplified by PCR from NMII gDNA using oligonucleotides that incorporate a C-terminal V5 tag. The resulting *cbuA0028*-V5 PCR fragment was cloned into pET28a(+) digested with NcoI/XhoI using In-Fusion HD to create pET28a(+)-*cbuA0028*.

*cbuA0028* and *cbuA0027* promoter analysis vectors

The promoterless *mScarlet-i* gene was amplified by PCR from pmScarlet-i_C1 (Addgene #85044) and the resulting fragment cloned into EcoRI/XhoI-digested pMiniTn7T-*lysCA* using In-Fusion HD to create pMiniTn7T-*lysCA*-NoP-*mScarlet-i*. The *cbuA0027* promoter fragments 1-3 were amplified by PCR from QpH1 plasmid DNA. The *mScarlet-i* gene was amplified from pmScarlet-i_C1 and cloned with the *cbuA0027* promoter fragments 1-3 into EcoRI/XhoI-digested pMiniTn7T-lysCA using In-Fusion HD to create pMiniTn7T-*lysCA*-P*cbuA0027*-Frag-1-*mScarlet-i*, pMiniTn7T-*lysCA*-P*cbuA0027*-Frag-2-*mScarlet-i* and pMiniTn7T-*lysCA*-P*cbuA0027*-Frag-3-*mScarlet-i*. The *mScarlet-i* gene was fused by PCR to the *cbuA0027*-Frag-4 and *cbuA0028* promoters using overlapping oligonucleotides. The resulting *cbuA0027*-Frag-4-mScarlet-i and *cbuA0028*-mScarleti-I PCR products were cloned into EcoRI/XhoI-digested pMiniTn7T-lysCA using In-Fusion HD to create pMiniTn7T-*lysCA*-P*cbuA0027*-Frag-4-*mScarlet-i* and pMiniTn7T-*lysCA*-P*cbuA0028*-*mScarlet-i*, respectively.
